# Mathematical Modeling of the Canonical Aryl Hydrocarbon Receptor Pathway in murine bone-marrow-derived macrophages

**DOI:** 10.64898/2026.05.05.722708

**Authors:** Vincent Wieland, Tobias Blum, Irina Iriady, Laia Reverte-Salisa, Dilan Pathirana, Irmgard Förster, Heike Weighardt, Jan Hasenauer

**Affiliations:** Bonn Center for Mathematical Life Sciences, University of Bonn, Bonn, Germany; Life and Medical Science Institute, University of Bonn, Bonn, Germany

## Abstract

The aryl hydrocarbon receptor (AhR) is a ligand-activated transcription factor involved in xenobiotic sensing, as well as development, immunity, and tissue homeostasis. AhR signaling proceeds through a canonical and a non-canonical branch. It remains unclear which biochemical steps of the canonical pathway are responsible for distinct ligand specific transcriptional responses. Here, we developed a mechanistic ordinary differential equation model of the canonical AhR pathway in murine bone-marrow-derived macrophages. We calibrated it to time-resolved qPCR measurements of *Cyp1a1* and *Ahrr* mRNA in mouse bone-marrow-derived macrophages exposed to three structurally diverse, environmentally relevant ligands with known immunomodulatory activity: 3-methylcholanthrene, indolo[3,2-b]carbazole and bisphenol A. Calibration used global optimization under a heteroskedastic likelihood. To dissect ligand specificity, we evaluated 528 distinct candidate models that allow one or two ligand-involving reaction rate constants to vary. Model selection revealed a dominant regime governed by faster promoter binding. Furthermore, it supported that ICZ requires no ligand-specific parameter relative to 3MC and most differences are visible in the expression levels of *Ahrr* mRNA. In addition, the model revealed that differences at the promoter binding and transcriptional levels were better supported than upstream signaling events. The resulting model is made available in SBML and PEtab formats for reproducibility, and to enable further research into whether ligand-specific effects are conserved or rewired across cell types.

## Introduction

Cells are routinely exposed to a wide range of environmental compounds that can perturb cellular function and contribute to adverse outcomes, including carcinogenic effects. Many enzymes have evolved to biotransform and eliminate such compounds [1]. Their expression is frequently induced by xenobiotic- and stress-sensing transcriptional regulators, including the aryl hydrocarbon receptor (AhR) [2], the pregnane X receptor (PXR) [3], the constitutive androstane receptor (CAR) [4], and nuclear factor erythroid 2-related factor 2 (NRF2) [5]. Together, these regulators form a connected response system that couples chemical sensing to transcriptional reprogramming of metabolism, transport, and cellular stress defenses. Among these sensors, AhR is often considered to be of particular importance because it integrates signals from a chemically diverse ligand space and controls hallmark detoxification genes while also impacting broader physiological processes such as development and immunity [6, 7].

AhR is a ligand-activated transcription factor of the basic helix-loop-helix/per-ARNT-Sim family with central roles in xenobiotic metabolism, development, and immunity [6, 7]. In its inactive state, AhR resides in the cytosol as part of a multiprotein chaperone complex, primarily consisting of a heat shock protein 90 (HSP90) dimer, the co-chaperone protein p23, and the AhR-interacting protein (AIP). Upon ligand binding, the receptor undergoes a rapid conformational change that exposes its nuclear localization signal, triggering translocation into the nucleus. Current models suggest that the tyrosine kinase c-Src dissociates from the AhR complex in the cytoplasm, while HSP90, p23, and AIP remain associated with AhR during nuclear import and are released in the nucleus upon, or during, the formation of a heterodimer with the AhR nuclear translocator (ARNT) [8, 9]. This heterodimer then binds to xenobiotic response elements (XREs) in the promoter regions of target genes. This process facilitates the robust transcriptional regulation of the ‘AhR gene battery’, including cytochrome P450 enzymes (e.g. *Cyp1a1* ) and the AhR repressor (AhRR) [2, 10]. This DNA-binding transcriptional arm is commonly referred to as the canonical or genomic AhR pathway. Beyond the canonical route, AhR signalling encompasses additional regulatory branches and molecular interactions that shape downstream responses in a context-dependent manner, the so-called non-canonical pathway [2, 6, 7]. After transcriptional activation, AhR is exported from the nucleus and degraded through the ubiquitin–proteasome pathway [11, 12], and the target gene TiPARP/PARP7 provides a further, transcriptionally encoded negative feedback on receptor abundance [13].

AhR ligands span a broad chemical spectrum, ranging from endogenous and dietary compounds to environmental pollutants and synthetic chemicals. Dietary precursors such as indole-3-carbinol (generated from glucosinolates in cruciferous vegetables) can be converted into high-affinity AhR agonists, including indolo[3,2-b]carbazole (ICZ) [14]. For experimental and quantitative studies, in particular, ligands with distinct origins and activities are informative: ICZ represents a diet-derived, high-affinity agonist [14]. Bisphenol A (BpA), an industrial co-monomer used in polycarbonate plastics, is not a classical AhR agonist but has been reported to have endocrine-disrupting and modulatory effects in AhR-related assays [15, 16]. Finally, the polycyclic aromatic hydrocarbon (PAH) 3-methylcholanthrene (3MC) is a commonly used synthetic AhR ligand and carcinogenic xenobiotic [17, 18]; as a high-affinity yet metabolically sensitive ligand, it is often used to probe early canonical AhR dynamics without the long-lasting stress responses associated with highly persistent ligands such as 2,3,7,8-tetrachlorodibenzo-*p*-dioxin (TCDD) [17–20]. Both 3MC and ICZ are high-affinity ligands of the murine AhR, with reported binding IC_50_ values in the low-nanomolar range — approximately 1.6 nM for 3MC and 0.2–3.6 nM for ICZ [21, 22] — whereas BpA acts substantially more weakly and in a context-dependent manner [23, 24] and is cleared predominantly by phase-II glucuronidation [25]. 3MC and ICZ are metabolically labile in comparison with persistent agonists such as TCDD, so that their cellular concentrations can decline appreciably over the course of an exposure [26, 27]. The three compounds therefore differ not only in the affinity with which they engage the receptor but also in how long they remain available to it.

Although the canonical AhR pathway follows a well-established sequence of events, it is well documented that different ligands can elicit distinct transcriptional response dynamics [28–30]. The mechanistic origin of these ligand-specific effects, however, remains unclear. Structurally diverse ligands display distinct interaction patterns with the AhR ligand binding domain and thereby may contribute to differential modulation of AhR functionality [31]. Classical high-affinity agonists are typically planar, hydrophobic and polycyclic, but the ligand spectrum of the receptor extends well beyond that structural class [32]. Recent structural work shows chemically diverse agonists occupying the ligand-binding pocket of the PAS-B domain within overall similar AhR–ARNT complexes. 3MC and ICZ were not among the ligands used for analysis of the ligated AhR–ARNT complex in that study, but mutating conserved residues of that pocket also altered AhR activation by 3MC and ICZ [8]. A common structural framework therefore accommodates the compounds used here, while ligand-specific contacts and conformational differences within the pocket can still influence receptor activation and may propagate into the transcriptional response [8, 32, 33]. Ligand-dependent differences could arise at multiple steps of the cascade, for instance through altered receptor complex stability, conformational effects that modulate co-factor recruitment, or changes in promoter binding and transcriptional efficacy. Disentangling which biochemical reaction steps are altered by a given ligand is challenging experimentally, because most measurements provide indirect, partial snapshots of the system and do not uniquely identify the causal reaction-level differences that generate the observed time courses.

In such settings, mechanistic mathematical models provide a principled framework for integrating heterogeneous data and testing hypotheses quantitatively. It has been shown that by formalizing pathway knowledge using mathematical models, one can connect reaction-level assumptions to measurable outputs, and evaluate which mechanistic variants are compatible with the data [34–37]. Mathematical models can even generate predictions that guide the design of informative follow-up experiments [38].

In this study, we employ mechanistic modeling to investigate ligand-specific regulation within the canonical AhR pathway by integrating time-resolved qPCR measurements of *Cyp1a1* and *Ahrr* mRNA from mouse bone-marrow derived macrophages (BMDM) exposed to 3MC, ICZ, and BpA. We first construct a baseline biochemical reaction network for canonical AhR signaling that includes promoter binding for the two target genes and dual negative feedback via CYP1A1-mediated ligand clearance and AhRR:ARNT repression, and we formalize this network as an ordinary differential equations (ODE) model under mass-action kinetics. Previous mechanistic AhR models represent explicit binding of AhR:ARNT complexes to DNA and recruitment of RNA polymerase [39]. The presented model combines this with dual-feedback, explicit promoter species and competition between activator and repressor complexes. We then generate a family of model variants from this baseline by allowing a small number of ligand-involving reaction rate constants to differ between compounds, calibrate each candidate to the experimental time courses using likelihood-based inference, and compare candidates. This model-selection framework identifies a constrained set of reaction-level mechanisms consistent with the data and suggests concrete directions for further research on the canonical AhR pathway.

The scope of this study is to model the canonical, DNA-binding arm of AhR signalling in murine BMDM. BMDM’s were chosen because AhRR is predominantly expressed in myeloid cells and because AhR/AhRR signalling shapes macrophage function under inflammatory and metabolic challenge [40–42]. Non-canonical AhR signalling, cross-talk with other ligand-activated receptors, and AhR functions in other cell types are not represented in the model and are therefore not addressed by our analysis. Restricting the pathway in this way keeps the number of competing mechanistic hypotheses small enough for the available transcriptional data to discriminate between them.

## Results

### Mechanistic model of the canonical AhR pathway

To assess how the response of the canonical AhR pathway to different ligands is regulated, we developed a dynamical mathematical model (Figure 1). The model tracks the life cycle of an AhR-ligand complex and its transcriptional consequences through the following core processes:

**Figure 1:**
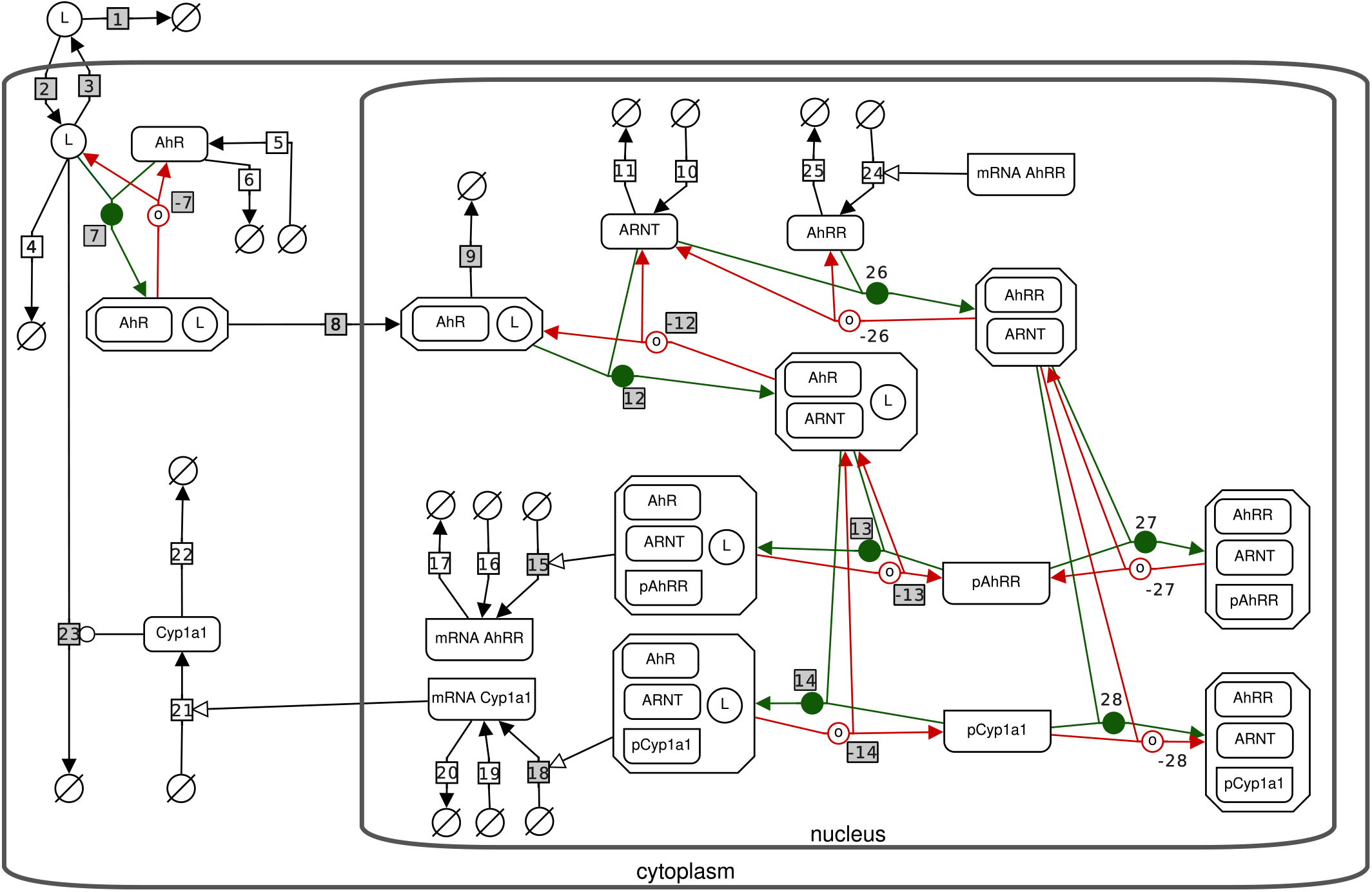
AhR signalling pathway model. Systems Biology Graphical Notation (SBGN) representation of the biochemical reaction network upon binding of the ligand, denoted by L. Reactions are numbered in order from 1 to 28, where the back reactions are indicated by a minus sign. All reactions that are allowed to be ligand-specific, are highlighted with a grey box.

#### Complex Formation and Translocation

Extracellular ligand (L) enters the cytoplasm where it binds to the AhR, forming the complex AhR:L. While AhR is known to reside in a multi-protein chaperone complex in the cytosol, we do not explicitly model these chaperones. Their high cellular abundance relative to AhR [43] and the reported rapid dissociation of the complex upon nuclear entry [44] suggest that chaperone dynamics are not rate-limiting.

#### Binding with ARNT

Following nuclear translocation, the multi-protein chaperone complex rapidly dissociates, providing ligand-receptor complex (AhR:L) in the nucleus, which enables binding to ARNT to form the active transcription factor complex (AhR:L:ARNT).

#### Promoter Binding

The active transcription factor complex binds to the promoter regions of target genes, forming activator–promoter complexes that drive the transcription of *Ahrr* and *Cyp1* family members. The transcription rate is defined as the sum of a basal production rate and a rate proportional to the concentration of the active promoter–activator complex. Each gene is represented by two promoter copies per diploid nucleus, corresponding to a total promoter concentration of 0.0332 nM in a nuclear volume of 10^*−*13^ L, and a single lumped response element is modelled per promoter copy. Because basal transcription enters as a constant, activator-independent flux, repressor occupancy suppresses the induced component of transcription but not the basal component.

Signal termination and adaptation are mediated by two negative feedback loops [45]:

#### Metabolic Feedback (via Cyp1a1)

The translated protein facilitates the metabolic clearance of various AhR ligands. While AhR regulates a suite of metabolic enzymes including Cyp1a2 and Cyp1b1, we utilize Cyp1a1 as a representative proxy for the metabolic feedback loop, as it is highly inducible in macrophages and exhibits overlapping substrate specificity with other *Cyp1* family members. For the classical agonists 3MC and ICZ this leads to a Cyp1a1-induced degradation of the cytoplasmic ligand. In contrast, for BpA, degradation is assumed to occur at a constant rate independent of Cyp1a1 activity, reflecting its status as a non-classical perturbant [16, 46].

#### Transcriptional Repression (via AhRR)

The AhR repressor protein (AhRR) acts as a competitive inhibitor. It binds to ARNT, forming a repressor complex (AhRR:ARNT) that competes with the active AhR heterodimer (AhR:ARNT:L) for the promoter sites [6]. These repressor–promoter complexes occupy the DNA without inducing transcription.

In total, the reaction network comprises 35 biochemical reactions. All reactions and their associated fluxes and rate constants are listed in Table S4 in the Supplementary Information S3. The fluxes are assumed to follow mass-action kinetics and account for the volume of the compartment in which they take place. Transport between the extracellular medium, the cytoplasm, and the nucleus therefore conserves amounts rather than concentrations.

From the network under consideration, a system of ordinary differential equations was derived, the so-called reaction rate equation [47]. This ODE system possesses 18 state variables, capturing the biochemical species in the different compartments across the considered cell population. The model does not possess single-cell resolution but rather has to be interpreted as a model of the population average. Hence, while a single promoter copy is at any instant in exactly one of three discrete states—free, bound by the activating AhR:L:ARNT complex, or bound by the repressing AhRR:ARNT complex—the state variables for the promoters are the expected concentrations of these states across the population of cells that contribute to one measurement. This coincides with the the quantity reported by the pooled-lysate qPCR readout. A fractional occupancy is therefore the fraction of promoter copies in a given state across the population, and the simultaneous presence of activator- and repressor-bound promoter complexes represents coexistence across cells, not within a single cell. The model yields a unique solution, and for non-negative initial conditions, this solution remains non-negative (see Supplementary Information S1, subsection S1.4).

To simulate ligand-induced responses, we assume that the system is initially at a basal steady state, representing the unperturbed condition prior to exogenous ligand stimulation. This steady state was derived analytically as a parameter-dependent solution of the algebraic steady-state equations and is used to initialize all simulations. The full ODE system with 18 state variables and its unperturbed steady state are provided in the Supplementary Information S1 and as an SBML file.

### Parameterized model reproduces ligand-specific transcriptional dynamics

To assess whether the proposed mechanistic ODE model can quantitatively describe ligand-specific transcriptional responses of the canonical AhR pathway, we evaluated its ability to describe time-resolved gene expression data. The experimental data were collected from BMDM stimulated with three chemically distinct compounds: the classical AhR agonists 3MC and ICZ, and the non-classical perturbant BpA. Cells were exposed to 1 *µ*M 3MC, 10 *µ*M ICZ, or 1 *µ*M BpA and dimethyl sulfoxide (DMSO) control. Relative mRNA abundance of the canonical AhR target genes *Cyp1a1* and *Ahrr* were measured by qPCR at six post-stimulation time points between 2 and 24 hours and normalized to DMSO control measurements. Each experimental condition comprised three biological replicates, each measured in technical triplicate and each paired with its own vehicle control (see Materials and Methods). The experimental schedule, molecular structures of the ligands, and corresponding measurement data on relative ΔΔ*C*_*T*_ value are shown in Figure 2. Upon stimulation with 3MC, *Cyp1a1* expression increased to about 15 fold as compared to the DMSO control over the complete time course. After stimulation with ICZ, *Cyp1a1* expression clearly peaked at 4 h after stimulation and declined after the 4 h incubation time point. *Ahrr* expression, however, was highest at the 3 h time point after activation with either 3MC or ICZ and also declined at later time points. Both *Cyp1a1* and *Ahrr* expression show lower responses over the observation period after stimulation with BpA compared to the other ligands. The measurements provide relative, but not absolute, mRNA abundances. Accordingly, scaling factors were introduced to map model-predicted mRNA concentrations to the relative qPCR readouts.

**Figure 2:**
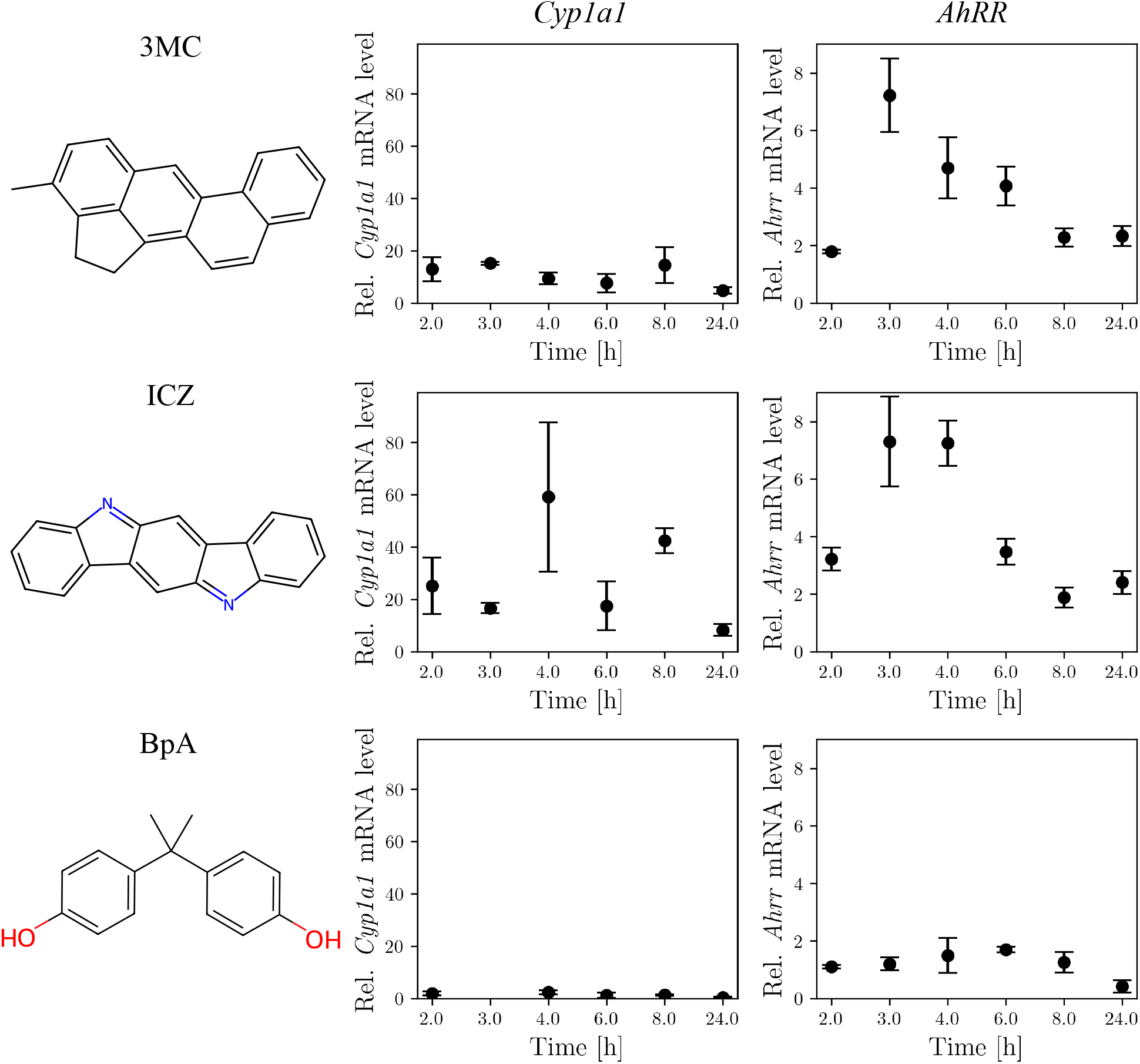
Ligands and corresponding qPCR-measurement for the two target genes. Lig- and structures and relative expression on ΔΔ*C*_*T*_ levels. Dots represent the mean over the replicates and bars depict the standard error of the mean (sem).

First, we assessed the suitability of our proposed model in capturing the dynamic response to each individual ligand. Therefore, we calibrated the model for each ligand-specific data set independently. Model calibration was performed by maximum likelihood estimation of kinetic, scaling, and noise parameters using global optimization (see Materials and Methods). Parameter estimation problems were specified using the PEtab standard [48] and solved with a self-adaptive cooperative enhanced scatter search algorithm combined with local, gradient-based optimization [49] (see Supplementary Information S1, S1.5). Parameters that are structurally non-identifiable due to the lack of absolute concentration measurements were fixed or reparameterized prior to optimization; details of these parameter reductions are provided in the Supplementary Information S3 (Table S5). Across all three datasets, the lowest recorded objective values form a flat group separated from the remainder (Figure 3 (a)–(c)). Such a group indicates an optimum that the search revisited repeatedly.

**Figure 3:**
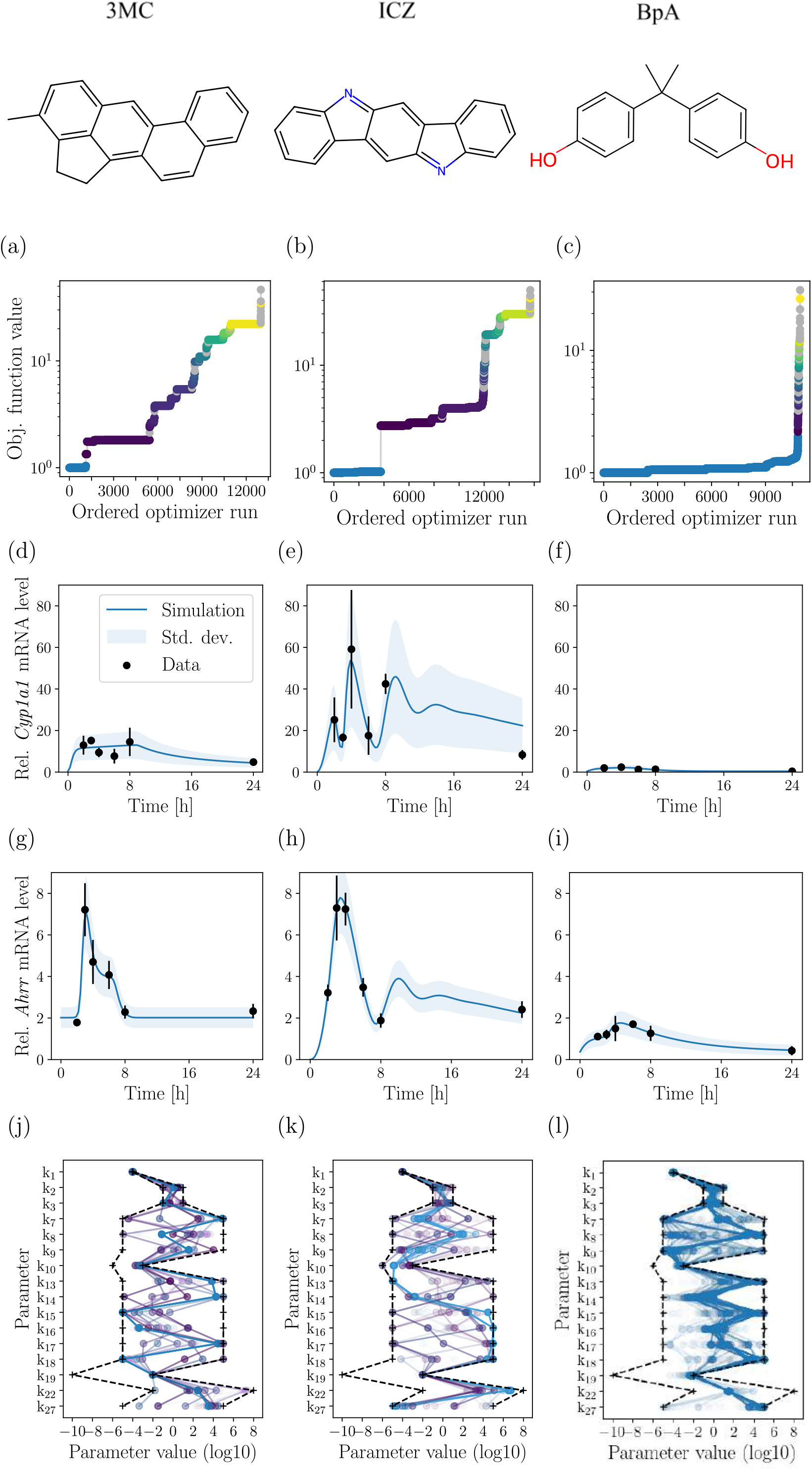
Results for fitting each ligand separately. (a)–(c) Recorded optimization end points, sorted by negative log-likelihood on its absolute scale; the best value is marked in red. (d)–(i) Visualization of the model fits by optimized model’s simulation (blue) and data (black) for each experimental condition separately. Shaded areas and error bars represent the standard deviation of the noise model or the data replicates, respectively. (j)–(l) Best-fit estimates (blue) of the 16 ligand-involving rate constants; the BpA panel includes the additional constant cytosolic ligand degradation parameter *k*_4_. Dashed black traces mark the parameter bounds. Colours in (a)–(c) encode objective values; the blue traces in (j)–(l) show the best estimates only.

Using the optimal parameter estimates, we simulated *Cyp1a1* and *Ahrr* mRNA trajectories and compared them to the experimental observations (Figure 3 (d)–(i)). For all three ligands, the model reproduces the main temporal features of the measured transcriptional responses. For ICZ, the model slightly underestimates the final *Cyp1a1* mRNA value; however, this data point remains within one standard deviation of the simulated distribution. The vehicle control against which these values are normalised is itself variable, and that variability propagates into every normalised measurement. The heteroskedastic form used here was selected from a comparison of seven observation-error models. The objective function and this comparison are defined in Materials and Methods.

To obtain lower bounds on parameter uncertainty, we constructed parameter ensembles by collecting parameter vectors from the optimizer history that fall within the 95% likelihood-based confidence region (see Materials and Methods). Simulations from this ensemble show good visual agreement with the experimental data (Supplementary Information S2, Figure S2). Analysis of the ensemble revealed pronounced differences between ligand-specific parameter estimates (Figure 3 (j)–(l)). At the same time, many parameters remain weakly constrained, indicating practical non-identifiability. A likely cause is the limited number of measured observables and time points relative to the number of unknown parameters.

Together, these results demonstrate that the proposed pathway model can robustly capture the observed transcriptional dynamics and provides a suitable foundation for subsequent model-selection and mechanistic analyses, while highlighting substantial remaining parameter uncertainty. The same holds when the three datasets are described jointly by a single parameter set with a small number of ligand-specific rate constants, as in the model selected in the following section (Figure 5): for all three ligands the simulated trajectories follow the induction of both target genes within the first hours, the earlier peak of *Ahrr* relative to *Cyp1a1*, the subsequent decline towards a plateau, and the markedly lower response amplitude under BpA than under the two classical agonists.

**Figure 4:**
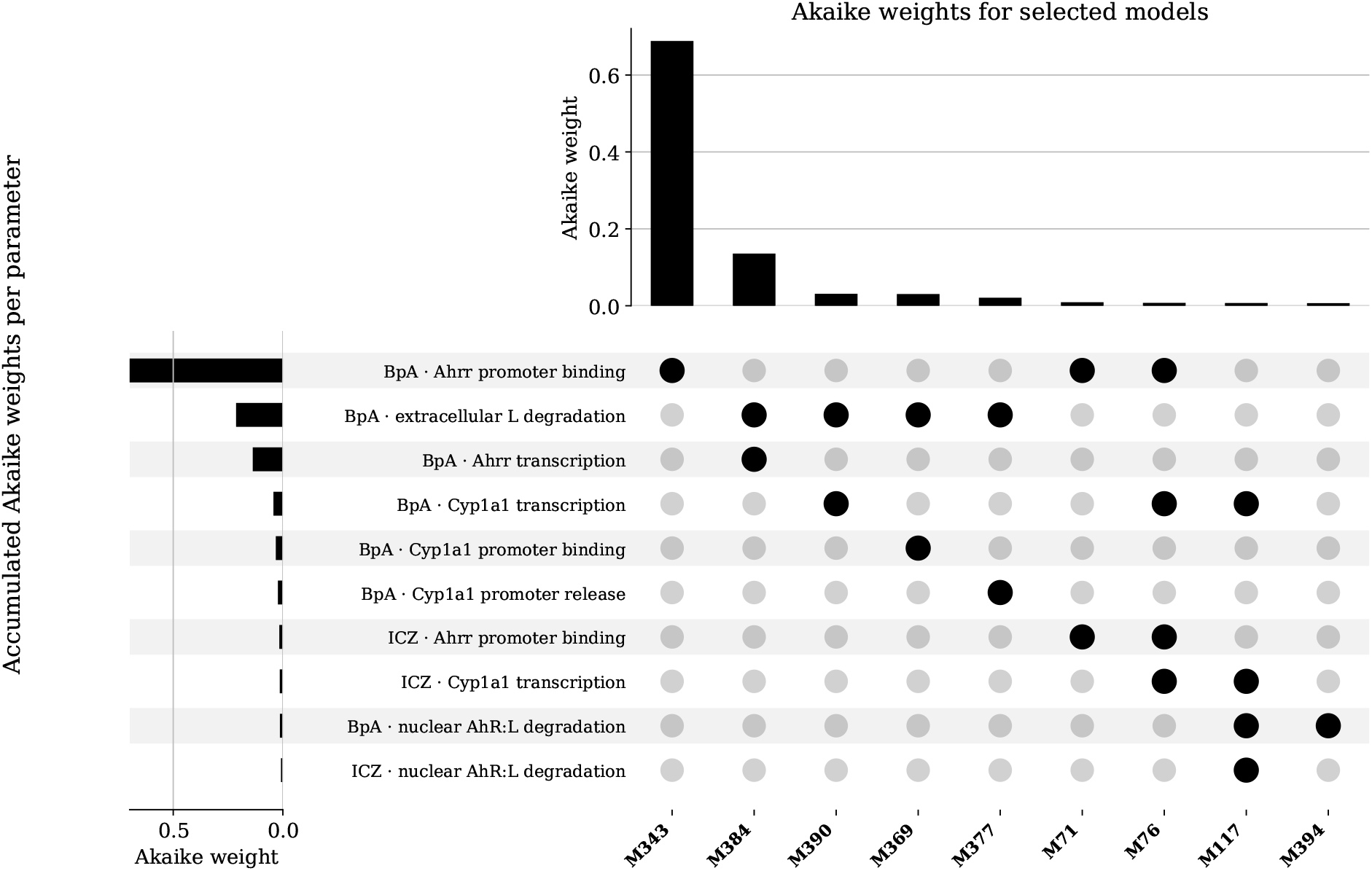
Model selection results. UpSet plot showing which combinations of ligand-specific reaction rate constants (rows) the calibrated models (columns) allow for. Each row is one rate constant together with the ligand whose fold-change is freed. Each column is one combination of such rate constants, labelled by its best-ranked model. The upper bar plot shows the Akaike weight of each combination, computed from the corrected Akaike Information Criterion and summed over all calibrated models that share the combination.

**Figure 5:**
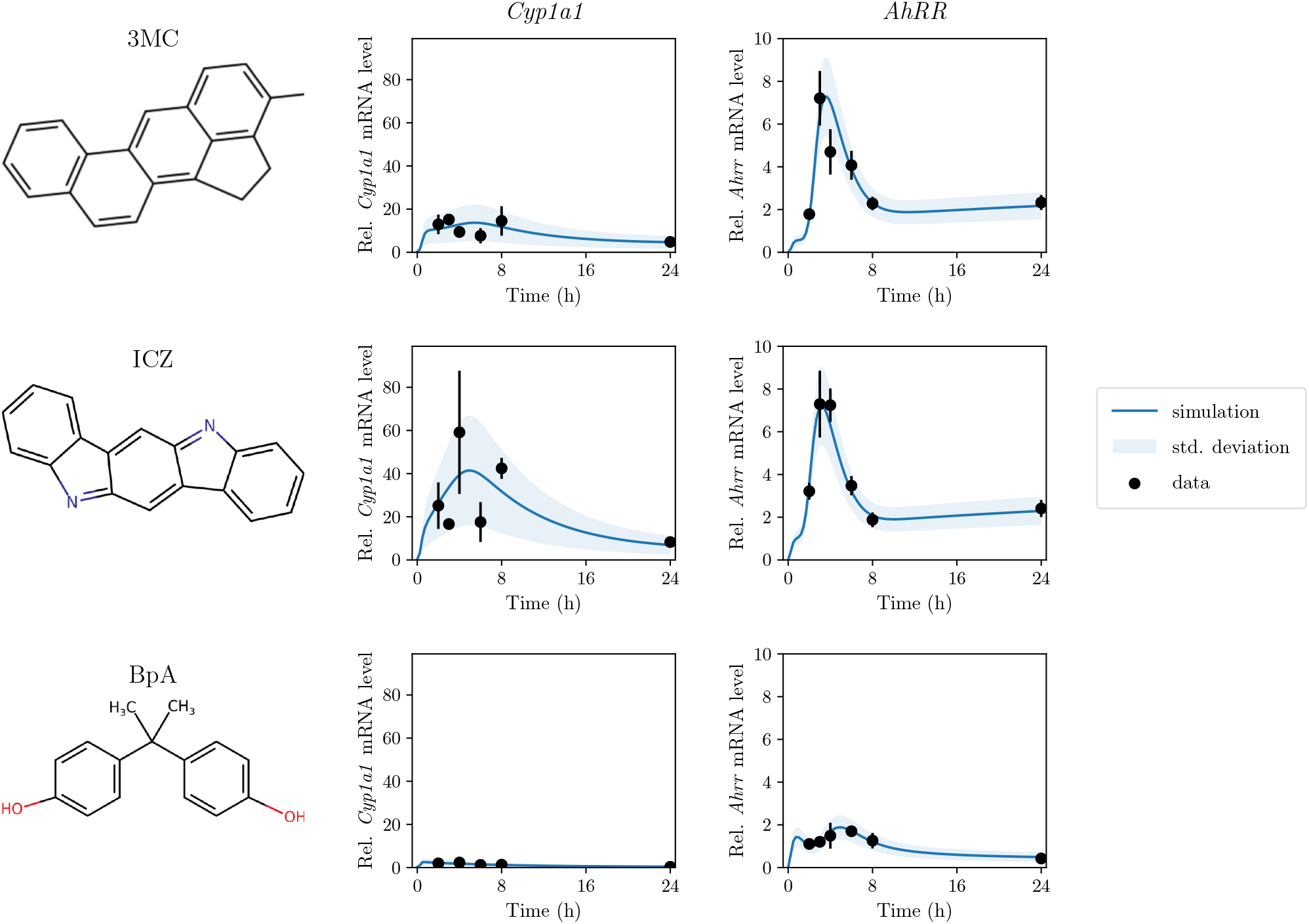
Model fit for the best model. Model fit given by the optimized parameter vector of M343, the best model. Rows show the three ligands (3MC, ICZ and BpA, with their chemical structures on the left), columns the two observables (*Cyp1a1* and *Ahrr* mRNA, relative ΔΔ*C*_*T*_ on a linear scale). Lines are the simulated observables. The shaded band around each line is the fitted observation-noise model evaluated along that trajectory, that is, the simulated observable plus and minus one standard deviation *σ*_*i*_(*y*_*i*_(*t*)) of the heteroskedastic error model at the optimized parameters. Dots are the mean over the three biological replicates at each measurement time, with error bars giving the standard error of that mean.

### Model selection implicates ligand-specific regulation at the promoter level

To pinpoint which biochemical reaction steps are most likely responsible for ligand-specific transcriptional responses, we performed a systematic model selection based on a shared mechanistic pathway model. Rather than fitting each ligand individually, the datasets for the three ligands (3MC, ICZ, and BpA) were fitted jointly, while allowing only a subset of reaction rate parameters to differ between ligands. This approach enables a direct comparison of alternative hypotheses regarding the steps of the pathway at which ligand-specific effects occur, while retaining a common mechanistic backbone.

We focused on reactions that directly involve the ligand or ligand-containing complexes, as these represent plausible points at which ligand-specific regulation may arise. In total, 16 such reactions are contained in the model (highlighted in Figure 1). Ligand-specific effects were encoded through fold-change parameters that modify baseline reaction rates, allowing either (i) the rate constant for one ligand to differ from the other two or (ii) the rate constant for all three ligands to differ. In addition, we allow ligand-specific effects in the degradation mechanisms, motivated by the aforementioned, well-established differences between BpA in comparison to 3MC and ICZ. The complete model space includes 262,124 models with different combinations of ligand-specific parametrizations. In a first step to constrain the model space, we applied the efficient strategy of forward–backward model-selection using the Akaike Information Criterion (AIC) [50], starting from the fully shared model and iteratively introducing or removing ligand-specific parameters. Across these runs, model complexity consistently collapsed, with selected candidate models containing at most one or two ligand-specific parameters. This result indicates that introducing more than two ligand-dependent parameters does not yield a relevant improvement in fit to justify the increased complexity.

Motivated by this result, we subsequently performed an exhaustive search over the space of models in which exactly one or two of the 16 ligand-involving reactions were allowed to differ between ligands. This restriction ensures comprehensive coverage of the most parsimonious mechanistic hypotheses while keeping the model-selection problem computationally tractable. The search enumerates 528 distinct candidate models, and all of them were calibrated using the same parameter estimation pipeline, parameter bounds, and noise assumptions as described in the previous section (Supplementary Information S1, Section S1.9). Each model was optimized in three phases: a gradient-based multistart, a self-adaptive cooperative enhanced scatter search, and a final local refinement (see Materials and Methods).

Model comparison using the AIC identified 9 models that were substantially better supported by the data than the remaining candidates (Figure 4). The best model, M343, accounts for 0.69 of the total Akaike weight and contains a single ligand-specific reaction rate: the association rate of the activated AhR:L:ARNT complex with the *Ahrr* promoter (R_13_ in Figure 1) is 9.08-fold higher for BpA than for the two classical agonists.

The calculation of the profile likelihood for the fold-change yielded a 95% confidence interval of [3.38, 36.1], which excludes unity, so the acceleration is supported by the data. The second-best model, M384 (ΔAICc 3.27, Akaike weight 0.13), instead makes the *Ahrr* transcription rate and the extracellular half-life of the ligand BpA-specific. Both leading models therefore place ligand specificity on the *Ahrr* arm of the pathway, and attribute it to BpA alone. Simulations of M343 provide a good quantitative description of the experimental data for all three ligands simultaneously (Figure 5).

This pattern is also supported by the analysis of the remaining models. Accumulating Akaike weights over all candidates that contain a given ligand-specific reaction (Figure 4), the binding of the activated complex to the *Ahrr* promoter carries by far the largest weight (0.70), followed by the extracellular ligand half-life (0.21), the *Ahrr* transcription rate (0.14), the combined binding to and release from the *Cyp1a1* promoter (0.076) and the *Cyp1a1* transcription rate (0.050). Grouping reactions by the level at which they act, an accumulated weight of 0.73 falls on models whose ligand-specific reactions are confined to the promoter and transcription level. Models with ligand-specific parameters for any upstream reaction carry 0.27, while the respective subset with ligand-specific parameters only for upstream reactions carries 0.027.

The same accumulation over ligands shows that the ligand-specific effect is confined to BpA. Models with at least one BpA-specific parameter carry an accumulated weight of 0.97, and models in which BpA is the only deviating ligand carry 0.93. Within the resolution of these data, ICZ and 3MC are thus described by one common parameter set and differ only in the applied ligand concentration, whereas BpA requires a distinct rate at the *Ahrr* promoter. Mechanistically, this places the ligand-dependent difference at the step at which the activated receptor complex engages the promoter, that is, in how efficiently a given nuclear signal is converted into promoter occupancy, rather than in ligand uptake, receptor binding, nuclear import or heterodimerisation.

As the optimization problems are challenging, even when using advanced methods such as self-adaptive cooperative enhanced scatter search combined with gradient-based refinement, we assessed the robustness of the model-selection results by rerunning the optimization of all selected models from twelve independent starting points with different random seeds. For all 9 models, the optimal objective functions found during the model-selection procedure were on average comparable to the objective functions found in the confirmatory optimization runs (Table S12). This confirms that the optimization during the selection procedure was successful and therefore the reliability of the model selection results.

To further confirm that allowing at most two ligand-specific reactions was sufficient, we assessed the models with three ligand specific reactions neighboring the selected models. Specifically, we calibrated 29 candidate models, each obtained by adding one ligand-specific reaction to one of the two best models, using the same optimization protocol. All 29 models performed worse than their respective parent models, with an increase in AIC of at least 10.14.

Taken together, this model selection narrows down the space of relevant ligand-specific mechanisms and identifies the engagement of the *Ahrr* promoter by the activated receptor complex as the best-supported ligand-dependent step. The two classical agonists do not require any ligand-specific rate constant relative to one another, so that within the canonical pathway, and at the concentrations used here, their different transcriptional responses are accounted for by the applied ligand dose alone, while the non-classical perturbant BpA is not.

### Clustering reveals a dominant dynamical regime across models

As our model selection identified several parameterizations that explain the observed ligand-specific transcriptional responses comparably well, we assessed whether these alternative mechanisms share common dynamical characteristics. To this end, we analyzed both observable and state trajectories of the 9 selected models. Rather than restricting the analysis to point estimates, we accounted for parameter uncertainty by constructing ensembles of parameter vectors from the optimization histories of all 9 calibrations of each of these models. For each retained combination of model and parameter vector, we simulated the three ligand conditions and subjected the resulting *Cyp1a1* and *Ahrr* mRNA trajectories to hierarchical agglomerative clustering based on pairwise Euclidean distances (see Materials and Methods). The number of clusters was determined as the largest *k* for which no cluster contained fewer than 1% of all simulated trajectories, yielding 2 clusters (Figure 6(a)). This choice ensures that the identified clusters correspond to robust dynamical regimes rather than isolated outlier simulations.

**Figure 6:**
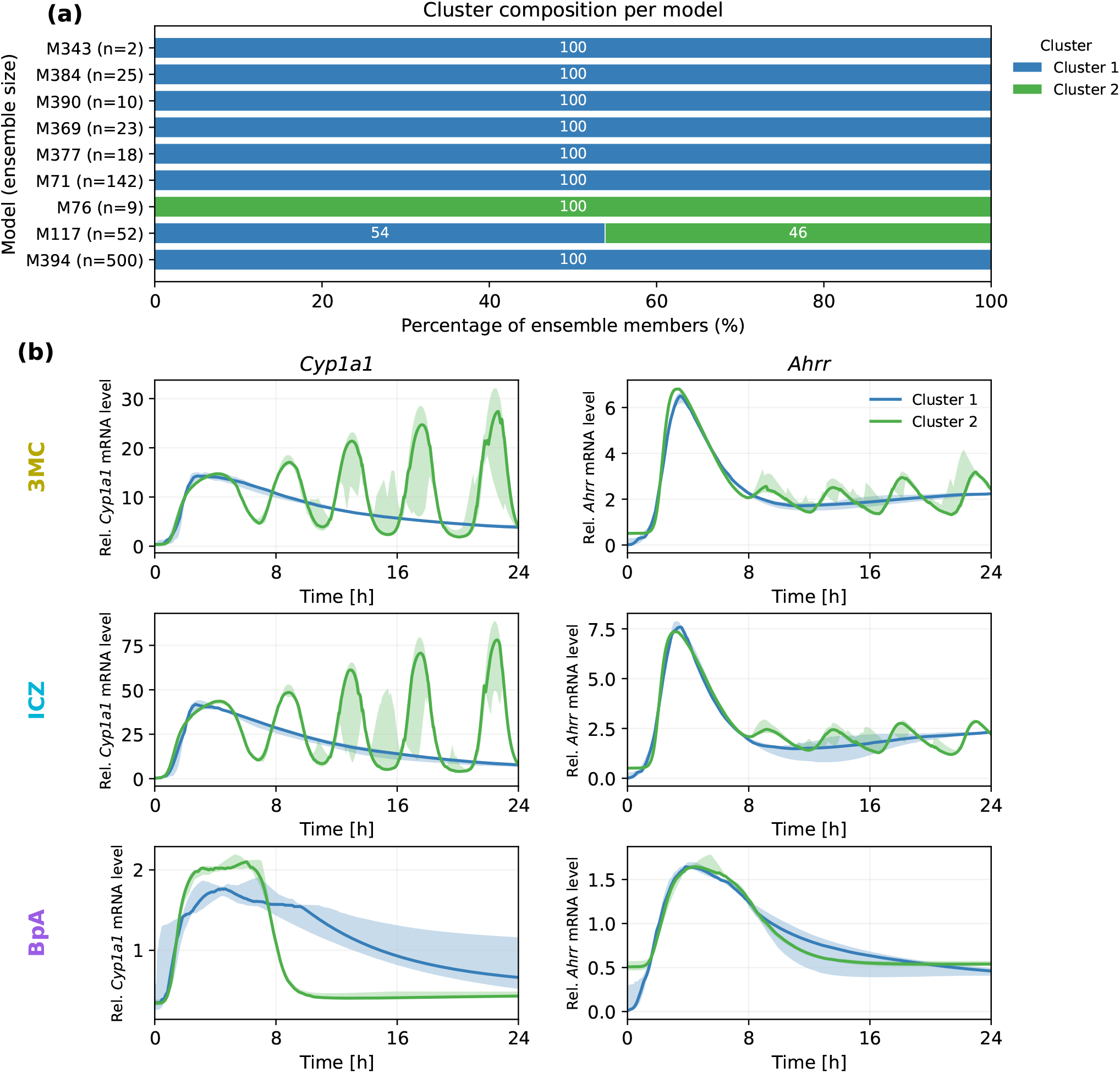
Clustering of the ensemble trajectories of the retained models and M360. (a) Contribution of each of the 9 models — the 9 retained models and M360 — to the 2 clusters, as a percentage of that model’s own ensemble; the number *n* in each model label counts the ensemble members of that model, i.e. the distinct recorded optima of its optimization history inside the 95% likelihood-based confidence region. Clusters are numbered by Akaike weight, not by member count. (b) Observable trajectories per cluster, pointwise median with the 25th and 75th percentiles as shaded areas; rows are the three ligands, columns the two observables.

To assess the overall importance of the two clusters, we constructed a contribution-metric based on accumulating the Akaike weights of all models in a cluster normalized with respect to the shares of the models that contribute to the cluster (see Supplementary Information S1, S1.11). This metric showed that main cluster (cluster 1) carries 99.1%, while holding 748 of the 781 simulated trajectories, i.e. 95.8% of them. It contains members of 8 of the 9 models (M117, M343, M369, M377, M384, M390, M394, M71), and for 7 of these (M343, M369, M377, M384, M390, M394, M71) it holds the entire ensemble (Figure 6(a)). The secondary cluster (cluster 2) carries 0.9%, so that the statistical support and the great majority of the individual trajectories rest on the same single dynamical regime.

The secondary regime (cluster 2), which is formed by members of M117, M76, shows sustained oscillations of *Cyp1a1* and *Ahrr* mRNA that set in after about 8 h, with a period of roughly four to five hours and a growing amplitude, where the dominant regime declines monotonically (Figure 6(b), Supplementary Information S3, Figure S4). These oscillations show good agreement with the data points such that the models that produce them survive the model selection at all. Nor are their parameter vectors a poorly fitting parts of the ensemble — every one of them attains a negative log-likelihood at or below that of M343 itself (212.75). The deviations between the clusters are thus concentrated where no data constrain them. Not one of the 6 comparisons of the two clusters in one observable under one ligand attains its largest separation at a measured time point, but all of them peak in a gap between measurements, 67% inside the 8–24 h gap alone. The largest separation of the secondary from the main regime amounts to 6.3 within-cluster standard deviations, in Cyp1a1 under ICZ at 22.6 h, and across all comparisons a median factor of 4.49 in discriminating power is lost by sampling at the six measured times rather than at the most informative ones.

As there are no reports in the literature on a prolonged oscillatory or otherwise modal activation of the canonical pathway, cluster 1 seems to provide the more plausible dynamics of the canonical pathway. Inspection of the models’ contribution to this cluster shows which ligand-specific reaction rates it collectively possesses (highlighted in blue in Figure 7): 11 rate constants (BpA_const_L_degrad, BpA_c1_dna_ahrr, BpA_c_eL, BpA_c_AhRR, BpA_c_Cyp, BpA_c1_dna_cyp, BpA_c2_dna_cyp, BpA_c_nAhRL, ICZ_c1_dna_ahrr, ICZ_c_Cyp, ICZ_c_nAhRL), of which 8 are BpA-specific and 3 ICZ-specific. The only one that every model of the cluster frees is BpA_const_L_degrad, the constant cytoplasmic degradation of BpA, which every candidate model estimates by construction. The others are the extracellular half-life of BpA, the further BpA-specific promoter, transcription and nuclear-complex steps of the individual models, among them the *Ahrr* promoter binding of M343, and the 3 ICZ-specific rate constants, carried by the models at the low-weight end of the cluster.

**Figure 7:**
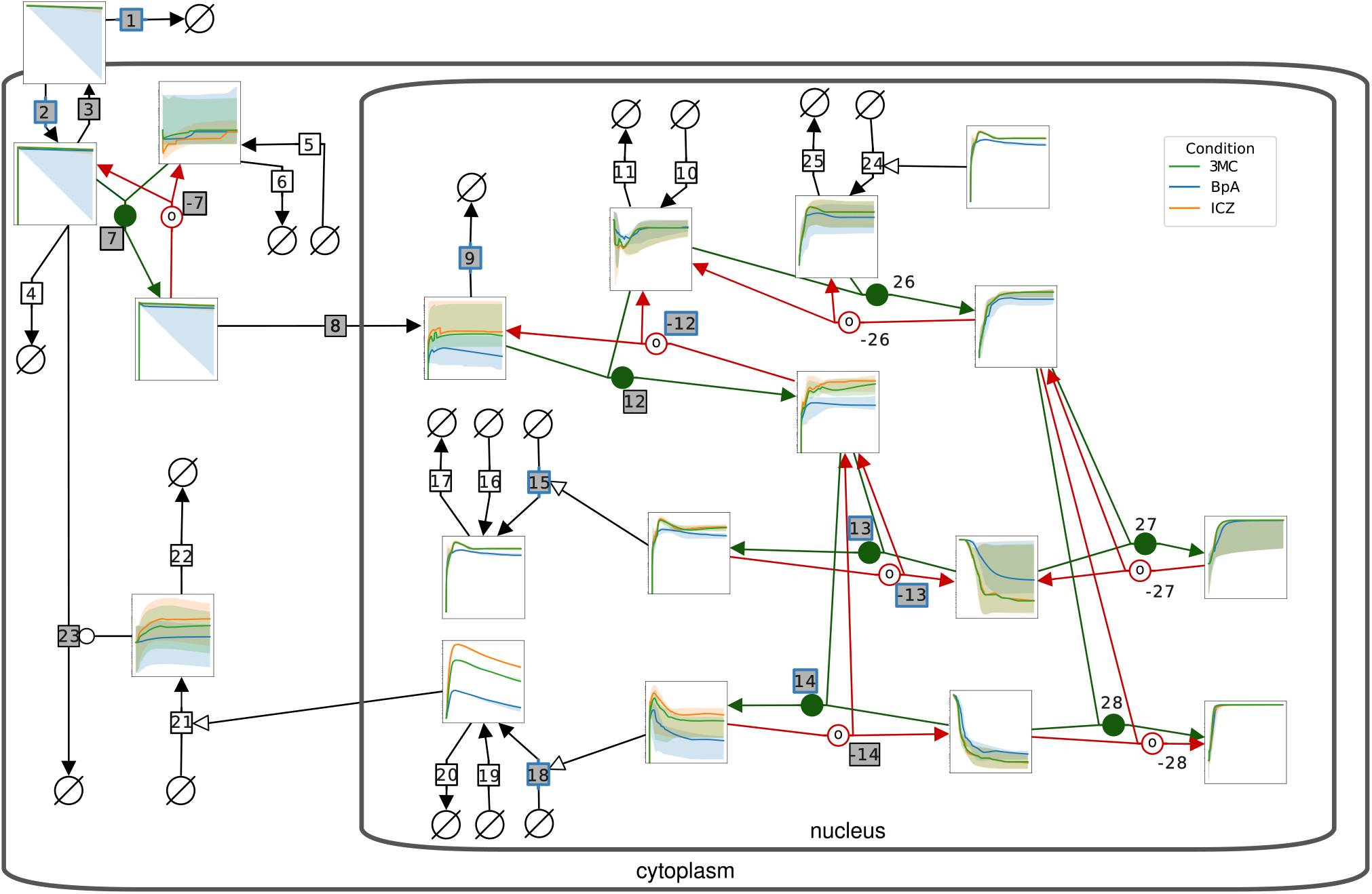
State dynamics for the selected cluster 1. SBGN model with state dynamics, median with (25%, 75%) percentiles, on log-scale for the main cluster. Reactions with grey background were part of reactions of interest and reactions with blue squares were selected as being different in the models represented by the main cluster.

To assess how these reaction-level differences translate into changes in unobserved biochemical species, we analyzed the simulated state trajectories for cluster 1. For each species and ligand, we computed pointwise ensemble intervals defined by the 25th and 75th percentiles of the simulations (Figure 7). Comparing the two clusters, 14 of the 18 species show a maximum separation exceeding 10% of their respective dynamic range. The largest relative difference is observed for nAhRR_ARNT_pCyp1a1, reaching 98% of its dynamic range, indicating that the clusters differ most strongly in the predicted occupancy of the *Cyp1a1* promoter by the repressor complex. When the separation is instead normalized by the variability within each cluster, the largest difference amounts to 14.6 within-cluster standard deviations and occurs for cAhR_L at 0.7h. Within cluster 1, the differences between BpA and the two classical agonists in the unobserved species are primarily associated with the ligand-specific *Ahrr* promoter binding in M343 and the BpA ligand half-life allowed to vary in a subset of the models, rather than with faster cytoplasmic ligand degradation alone.

Taken together, clustering of ensemble simulations reveals a dominant dynamical regime that carries almost all of the statistical support (99.1% of the Akaike-based contribution-metric) as well as 95.8% of the individual trajectories, and that holds 7 of the 9 models. The main dynamic is a single induction peak of both target genes within the first hours, followed by a monotone decline towards a plateau. The one alternative is a sustained oscillatory regime that the present measurement schedule cannot distinguish from it, because its swings fall between the sampling times and the two regimes coincide again at the last of them.

## Discussion

A central open problem in AhR biology is how chemically diverse ligands, ranging from xenobiotic ligands such as environmental pollutants, dietary components or drugs to endogenous ligands produced by the host or host microbiota, give rise to distinct transcriptional responses despite acting through a shared canonical signaling cascade. Disentangling where ligand specificity enters the path-way remains challenging, partially because experimental readouts are typically limited to a small number of target genes and time points. In this work, we addressed this problem by developing and quantitatively analyzing a mechanistic ODE model of the canonical AhR pathway that explicitly represents promoter binding and negative feedback, and by systematically comparing alternative mechanistic hypotheses using likelihood-based inference and model selection. All results refer to murine BMDM, and whether the same reactions carry ligand specificity in other AhR-responsive cell types remains open.

We show that the proposed ODE model can reproduce the main dynamical features of the transcriptional response across ligands using time-resolved qPCR measurements of two canonical AhR target genes in murine BMDM stimulated with ligands spanning a broad spectrum of chemical properties. The subsequent model selection identified candidate models that describe the available data for all three ligands with at most two ligand-specific reaction rates. The comparison of these candidates by AICc concentrated the majority of the evidence on one of them (Akaike weight 0.69), while the set of models that are statistically indistinguishable from the best one, an AICc difference of less than 10, comprises 9 models in total, which revealed that distinct parameterization can explain the data equally well.

The assumption that only one or two reactions differ between the ligands was justified from both directions. From above, the saturated model, in which every ligand-specific fold-change is estimated at once, improves on the best ranked model by 7.18 NLLH in the best fit we found. Since each further ligand-specific raises the AIC by 2, no candidate with one, two or three additional reactions can beat M343 by more than 12.35, 10.35 or 8.35 in the AIC. From below, a forward step calibrated every model that adds one further ligand-specific reaction to one of the three leading candidates. Every one of the 29 three-reaction models is worse than its parent, and none of them improved on the best found model. The restriction is therefore a property of the data underlying this work and further data may help to narrow the model selection even further.

In our analysis, the strongest supported source of ligand specificity is located at the level of nuclear signal processing rather than upstream signal transmission. The best-supported ligand-dependent step is the engagement of the *Ahrr* promoter by the activated AhR:L:ARNT complex, which proceeds faster for BpA than for the two classical agonists. Accumulated over the whole model space, reactions acting at the promoter and transcription level carry more evidence than models that locate ligand specificity exclusively upstream. Reactions governing ligand entry, receptor binding, nuclear import, and heterodimerisation do not require ligand-specific parametrization to reproduce the observed transcriptional responses. This suggests that the canonical AhR cascade up to nuclear localization is largely conserved across ligands in the presented setting for BMDM, whereas ligand-specific effects emerge primarily through differences in how the nuclear signal is integrated at the promoters of target genes. This is consistent with recent structural insights into ligand binding to the PAS–B ligand-binding domain [51]. Hence, our model-based analysis provides a mechanistic basis for how chemically distinct ligands may differentially influence receptor conformation and downstream engagement. Even in the presence of substantial parameter uncertainty, the model can therefore be used to assess pathway-level hypotheses and ligand-dependent regulatory mechanisms.

Notably, the model selection results provide no evidence that the two classical agonists, 3MC and ICZ, require distinct reaction rates within the canonical pathway at the resolution of the available data. ICZ requires no ligand-specific rate constant relative to 3MC. The accumulated Akaike weight of all models carrying any ICZ-specific parameter is 0.064, and that of all models carrying any 3MC-specific parameter is 0.006. The markedly different *Cyp1a1* and *Ahrr* time courses that the two compounds produce are thus accounted for by the applied concentration acting on a shared parameter set and a shared feedback structure. Ligand specificity in the sense of a distinct reaction rate is confined to BpA, with an accumulated weight of 0.93 on models in which BpA is the only deviating ligand. This is consistent with its status as a non-classical perturbant rather than a classical agonist, and with the different route of cytosolic clearance assumed for it.

Importantly, these conclusions must be interpreted in light of the model assumptions. While it is known that the promoters of AhR target genes such as *Cyp1a1* and *Ahrr* contain multiple AhR-responsive elements, we used a simple model for activation and inhibition. Structural studies of AhR:ARNT bound to DNA and studies of promoter architecture highlight the complexity of transcriptional regulation at response elements and underscore the relevance of our promoter-level modeling [52, 53]. The effective promoter representation used here captures the aggregated outcome of these interactions but cannot resolve site-specific binding or cooperative effects. Refining the promoter model is outside of the scope of this study, but highlights a key opportunity for future experimental designs: direct measurements of promoter occupancy or chromatin state would allow refinement of the promoter model and provide stronger constraints on transcriptional regulation. Additionally, transcription is modelled as a first-order mass-action process with no programmed dead time between promoter occupancy and mature transcript, an extension these data do not resolve (Supplementary Information S1, Section S1.8), and basal transcription enters as a constant, activator-independent flux, so repressor occupancy suppresses the induced component of transcription but not the basal one. Ligand binding is modelled as a single reversible step. This provides an effective description for the metabolically labile ligands considered here, whereas application of the model to highly persistent ligands such as TCDD may require a more detailed representation of ligand binding and clearance. Moreover, the model represents the canonical pathway, characterized by direct binding of AhR/ARNT to DNA, in murine BMDM only, and whether the same reactions carry ligand specificity in other AhR-responsive cell types remains open. Non-canonical AhR signalling and cross-talk with other ligand-activated receptors or with NF-*κ*B are not included. In particular, a modulation of *Cyp1a1* transcription mediated by oestrogen receptor *α*, which has been reported for AhR ligands, is not represented as a separate mechanism. Such an effect would act on the *Cyp1a1* transcription rate, and that reaction was part of the candidate space with an accumulated Akaike weight of only 0.050 over all models. The data therefore give little support to a ligand-specific modification of this particular step.

The results are further limited by data availability. For the specific target cells, we have only a single experimental design available in which each ligand at hand was applied at a single concentration. So, ligand-specific concentration–response behaviour cannot be separated from ligand-specific rate constants, and the reported fold-changes are conditional on the chosen doses and we cannot provide a validation against an independent dataset. To our knowledge, no published dataset matches this design closely enough in cell type, ligands, time resolution, and normalisation to serve as additional data or an external test set.

Parameter uncertainty is a further qualification, since the full saturated model is close to being overparametrized to resolve single fold-change estimates, 105 measurements for 71 parameters, and the obtained values should be interpreted with caution. Although, we performed an exhaustive literature search to constrain the parameter ranges using prior knowledge, many parameters remain weakly constrained by two mRNA observables at six time points, and the models of the retained set are separated by differences that are close to the AICc selection threshold. Profile likelihoods of M343 show that 13 parameters are identifiable, 26 are bounded on one side only and 2 are practically non-identifiable. However, the BpA-specific fold-change at the *Ahrr* promoter is identifiable.

To further pinpoint the parameters acting only through species the experiment never observes, which trade off against one another and against the scaling factors, requires observables of a different kind. For example, absolute or semi-quantitative protein measurements of AhR, AhRR and CYP1A1 to separate synthesis from degradation, direct promoter-occupancy readouts such as chromatin immunoprecipitation to constrain the binding and dissociation constants individually, and dose–response experiments to separate affinities from the rate constants with which they are currently confounded would help to resolve raised points.

Looking forward, our results suggest several concrete directions for both modeling and experimentation. On the modeling side, extending the framework to additional target genes, incorporating non-genomic AhR signaling, or explicitly modeling cross-talk with macrophage-relevant pathways could help to assess the generality of the inferred mechanisms. From an experimental design perspective, richer datasets, including absolute or semi-quantitative protein measurements, dose–response experiments, and direct promoter readouts, would substantially reduce uncertainty and enable sharper discrimination between competing mechanistic hypotheses. Furthermore, the clustering shows that the alternative to the dominant regime is a sustained oscillation that passes the data points closely, so it is invisible to the present schedule by construction (Supplementary Information S2, Figure S4). Two additional *Cyp1a1* measurements in the 8-24h gap, at about 9.6 and 22.7 h, are therefore the most discriminating experiment. The early phase before the first measurement remains worth sampling as well, since the unobserved species differ most strongly there. Joint inference across such datasets would also help to clarify whether ligand specificity is best described by differences in affinity, efficacy, or downstream transcriptional integration.

The feasibility of such future analyses is greatly enhanced by the availability of the present model and data for the canonical AhR pathway in standardized, interoperable formats. By providing the model in SBML and PEtab together with a fully reproducible calibration and model-selection pipeline, we aim to facilitate systematic model extension, data integration, and re-analysis as new experimental evidence becomes available. This framework enables *in silico* perturbation studies, exploration of combination treatments, and principled experimental design, and it supports long-term reuse and comparability across studies. As more quantitative data on AhR signalling accumulate, such integrative modelling approaches have the potential to deepen our understanding of how chemically diverse ligands shape immune and toxicological responses through a shared receptor system.

## Materials and Methods

### Experimental setup - data collection

The data was obtained from bone-marrow derived macrophages and the raw data is presented in the Supplementary Information (Supplementary Information S2, Figure S1; Supplementary Information S3, Table S3).

### Generation of bone-marrow derived macrophages

Bone marrow cells of 8–12 week old C57BL/6 JRCC mice were isolated and plated at a concentration of 5 *×* 105 cells/ml on RPMI 1640 (PAN Biotech), 10% FCS, L-glutamine, penicillin-streptomycin and *β*-mercaptoethanol. For differentiation into BMDM, 10% M-CSF from the L929 cell line was added. Cells were fed with the same medium on day 3 and harvested on day 6.

### Quantitative real-time PCR

BMDM (5×105 cells/ml) were stimulated with 1 *µ*M 3MC, 10 *µ*M ICZ, 1 *µ*M BPA or DMSO as solvent for 2, 3, 4, 6, 8 and 24 hours. The concentrations follow prior use of 1 *µ*M 3MC [54] and 10 *µ*M ICZ [26] in this system, both titrated in earlier experiments to give a comparably rapid induction of AhR signalling; 1 *µ*M BpA was taken from the literature as non-cytotoxic in reporter cell lines and human macrophages while still activating AhR [16, 24, 55]. RNA was isolated using the Zymo Quick-RNA Miniprep Kit (Zymo, Freiburg, Germany). First-strand cDNA was synthesized from 1 *µ*g of total RNA using a mixture of oligo(dT) and Revert Aid reverse transcriptase (Thermo Fisher Scientific Inc., Dreieich, Germany). mRNA expression was assessed by qRT-PCR using a Real-Time system CFX96 (BioRad) and Absolute qPCR SYBR Green ROX mix (Thermo Fisher Scientific) with primers given in Table 1.

**Table 1:**
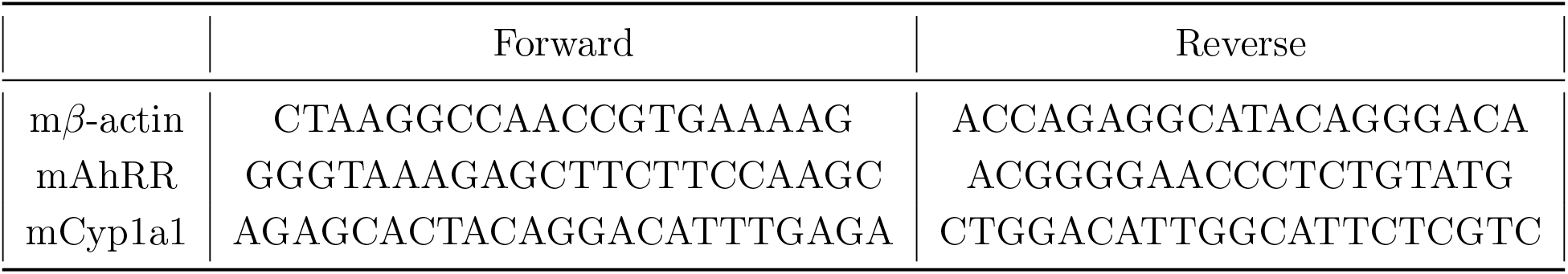
Primers used in qPCR.

The relative changes in gene expression are determined using the 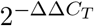 method [56]. This method compares a target gene in a treated sample normalized to a housekeeping gene and a control treatment. Using the housekeeping gene *β*-actin and as a control treatment dimethyl sulfoxide (DMSO), we got observations *y*_*i,k*_ for the *k*-th measurement timepoint *k* ∈ {1, …, 6} and the *i*-th target gene, *i* ∈ {1, 2}.

Three biological replicates were collected per condition and time point, each measured in technical triplicate, and each biological replicate comprised a treated sample together with its own paired DMSO vehicle control. The ΔΔ*C*_*T*_ values used for calibration were formed replicate-wise, dividing the 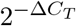 value of treated replicate *j* by that of control replicate *j*, rather than by the mean over the control replicates. Each normalised value therefore carries the full variability of a single control replicate rather than that of the control mean, which is the quantity the observation model has to account for.105 values were available for calibration, were three values were excluded, *Cyp1a1* under BpA at the 3 h time point, as documented in the Supplementary Information S1, Section S1.1.

### Model formalism

The biochemical reaction network was modeled using an ODE system of the form

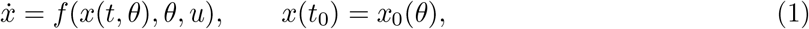

where 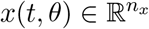 is the vector of concentrations of the biochemical species at time *t* and 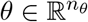 the parameter vector. The vector field 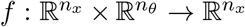 describes the temporal evolution of the concentrations and is constructed to be Lipschitz continuous in order to ensure uniqueness of the solution. We denote by *n*_*x*_ the number of species and by *n*_*θ*_ the number of parameters, e.g. rate constants. The initial condition *x*_0_(*θ*) is the state of the ODE at the time of the simulation start, i.e. the time of ligand treatment. To ensure that transient effects from the initial state of the system do not interfere with the signaling responses, i.e. the effects of interest, the initial condition needs to resemble the unperturbed state of the system before the ligand treatment. This unperturbed steady state *x*^*\**^ was derived analytically by solving the corresponding ODE system with the change of concentrations being 0,

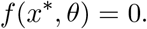

Additionally, it was assumed that all ligand-related compounds are at concentration 0 mol/L at the initial state and the first step in the model is the treatment with the ligand concentration corresponding to the experimental condition. The derived analytical solution can be found in Supplementary Information S1, S1.3.

### Modelling of the observation process

Biochemical species are assumed to be partially observed (using qPCR measurements) at discrete measurement timepoints {2h, 3h, 4h, 6h, 8h, 24h}. The link from unobserved states to the observables *y*(*t*) is represented by an observable mapping *h*,

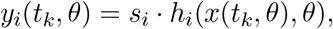

with *i* ∈ {1, 2} indexing the observables and *k* ∈ {1, …, 6} indexing the time points. Due to the relative nature of the measurements, it is necessary to introduce scaling factors *s*_*i*_ to scale the observables to the data. Measurement errors are captured using a Gaussian noise model with heteroskedastic variance

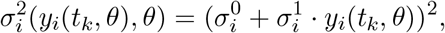

which represents a base measurement error via 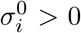 and a higher variance in the measurements for higher observed values with factor 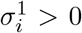. The observable mapping together with scaling and measurement error yields the following observation model for datapoints *y*_*i,k*_

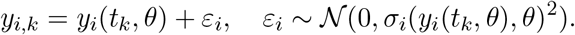

We selected this noise model after fitting seven observation-error models to the same measurements and compared them: the form given above, two further Gaussian forms with variance-additive and quadratic variance terms, a Laplace form, a log-normal form, and two variants in which the measured variability of the vehicle control is propagated explicitly into the likelihood. Because the log-scale variants describe a density over log_10_(*y*) rather than over *y*, all variants were placed on a common scale by the corresponding Jacobian correction before their information criteria were compared, and the comparison was accompanied by randomized quantile-residual diagnostics, which are standard normal for any correctly specified model regardless of its error distribution. The form given above ranked first. To confirm that this ranking is not an artefact of the particular ligand-specific parametrization it was established on, the comparison was repeated on four different model topologies, including the saturated one. The full comparison, the residual diagnostics, and the four-topology check are reported in the Supplementary Information S1, Section S1.7.

### Model parameterization

For reactions that directly involve the ligand or ligand-containing complexes, we introduce lig-and–specific fold-change parameters 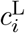 for each reaction *i* with rate *k*_*i*_ and ligand L, following Steiert *et al*. [57]. The rate for ligand L is thus given by 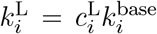, where 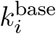 is a ligand-independent baseline rate. A fixed value of 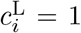 indicates that ligand L uses the baseline rate, whereas estimating 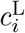 reflects a ligand–specific deviation (fold-change).

### Parameter estimation

Unknown model parameters (*θ, s, σ*) are inferred from the experimental data using maximum like-lihood estimation. Given a dataset 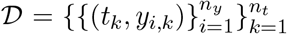 the likelihood function under the heteroskedastic Gaussian noise model is given by

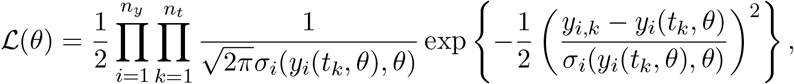

in which *x*(*t*_*k*_, *θ*) denotes the solution of the ODE system for the parameter vector *θ*.

For numerical stability and better convergence, we minimize the negative log-likelihood:

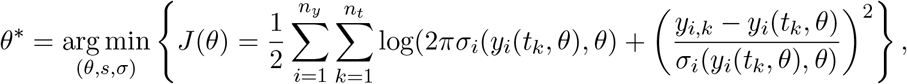

where *σ*_*i*_(*y*_*i*_(*t*_*k*_, *θ*), *θ*) denotes the heteroskedastic standard deviation.

The optimization was performed using a hybrid global–local optimization strategy implemented in pyPESTO [58]. Global exploration of the parameter space was conducted using the self-adaptive cooperative enhanced scatter search (saess) algorithm [49], a population-based metaheuristic that iteratively evolves a diverse set of candidate parameter vectors. In each global iteration, candidate solutions are combined and improved, and information is exchanged between cooperative subpopulations to enhance exploration while maintaining diversity and adaptively updating its internal search parameters balances exploration and exploitation. Promising solutions identified during the global phase are subsequently refined using the gradient-based interior trust-region reflective optimizer (FIDES) [59]. Gradients were computed using the adjoint sensitivities method for discrete-time measurements as outlined in [60]. Parameters were transformed to log_10_ scale to improve numerical stability during optimization [61–63]. Detailed information about the optimization options are provided in the Supplementary Information S1, S1.5 and all model parameters, with corresponding constraints are given in Supplementary Information S3, Table S5. Parameter bounds were set from the literature. Where a source was available, the bounds of a parameter were derived from it, and all bounds are reported per parameter in the Supplementary Information S1, Section S1.6).

Each calibration followed the same three-phase protocol. First, a gradient-based multistart from starting vectors sampled within the parameter bounds, second, a self-adaptive cooperative enhanced scatter search run on cooperating workers under a fixed wall-time budget, third, a local refinement of the best solutions found in either phase. A local optimization run is counted as converged when its final objective value lies within 10^*−*2^ of the best value found for that model. The objective *J* (*θ*) defined above is the quantity shown in all waterfall plots.

The number of converged start reported for the sweep are dominated by the records of the scatter search, and these are not independent draws. We therefore repeated the complete three-phase protocol 8 further times from independent seeds for each of the selected models giving 9 independent calibrations per model, and use these repetitions, for the convergence assessment of our model selection runs.

### Uncertainty analysis

The uncertainty in the model parameters and model fits were assessed by collecting all parameter vectors within a 95% confidence region of the MLE based on the likelihood-ratio test [64]. With the test statistic

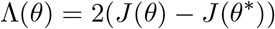

the confidence region of significance *α* is defined as

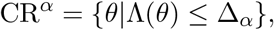

where Δ_*α*_ is the *α*-th percentile of the 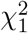 distribution. Based on this we created ensembles from the histories of the optimizers by collecting all parameter vectors lying in this confidence region and propagated this parameter uncertainty to model prediction.

The same definition was applied one coordinate at a time to obtain profile likelihoods [65]. Each selected parameter was fixed on a grid and the remaining parameters were re-optimised, so that the 95% confidence interval of that parameter is the range over which the re-optimised profile stays below Δ_0.95_ = 3.84. Profiling is expensive for this model, because each profile point re-optimises all remaining parameters at the solver tolerances used throughout. Therefore, we did run a full likelihood profile analyses only for the best model.

### Model selection

To determine which reactions are most likely account for ligand-specific dynamics, we fitted all three experimental datasets jointly while allowing 16 reaction rates to differ between the ligands as parametrized with the ligand-specific fold-change parameters 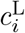.

For balancing goodness-of-fit against model complexity the Akaike Information Criterion (AIC) [66] was used, given by

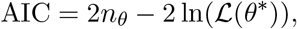

where *n*_*θ*_ is the number of estimated parameters and *L*(*θ*^*\**^) is the maximized likelihood. Since the number of measurements, *n* = 105, is not large relative to the number of estimated parameters, models were ranked by the corrected criterion

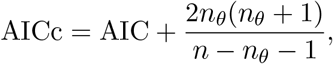

which adds a penalty that grows with *n*_*θ*_ and vanishes for *n*_*θ*_ ≪ *n*. Differences of 10 or more were considered substantial [67], and we also report the corresponding Akaike weights for comparing models. As an initial exploratory step, we applied the forward-backward model-selection strategy [50] without restricting the number of ligand-dependent reactions. Across these runs, models in which more than two reactions differed between ligands were never selected as good candidate models by the algorithm. This indicates that additional ligand-dependent reactions only increase complexity without yielding a substantial improvement in the fit. To make an exhaustive search computationally feasible, and without privileging any specific reaction or ligand, we therefore restricted the candidate model space to models in which exactly one or two of the 16 reactions were allowed to differ between ligands and constructed the candidate model space in a systematic way. For each of the 16 reactions, we considered it either ligand–independent (all 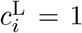) or ligand–dependent. In a ligand–dependent reaction differences were encoded by estimating either one or two fold-change parameters while keeping the remaining 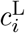 fixed at 1:

- If one fold-change parameter 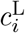 was estimated, exactly one ligand could differ from the other two (which share the baseline rate)
- If two fold-change parameters were estimated, all three ligands could in principle differ, because one ligand can be taken as baseline and the other two obtain independent fold changes.

For three ligands, this yields four qualitatively distinct patterns of ligand dependence for a given reaction. Importantly, we did not preselect any reaction-ligand combinations and every reaction was assigned any of these patterns, and all patterns were treated on equal footing. For a given model, we first chose the subset of reactions that could differ between ligands and then chose one of the four patterns of ligand dependence described above. The number of subsets with one or two reactions is therefore given by

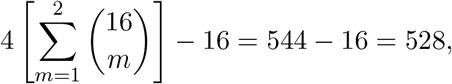

where we subtract 16 because we assumed a constant rate across ligands for the cytoplasmic degradation of BpA, i.e. the corresponding fold-change parameter was fixed to 1 in all models. This eliminates those model configurations in which this particular reaction is included in the set of lig- and–dependent reactions and BpA is the ligand allowed to deviate from the others. There are 16 such combinations.

All models within a AICc difference of 10 from the overall best model were considered statistically indistinguishable and were retained for further analysis, which yielded 9 candidate models. Those models were each given 8 independent repetitions of the complete three-phase protocol from different random seeds, so that 14 models have 9 calibrations each (Supplementary Information S3, Table S12). These repetitions assess the robustness and reproducibility of the calibration.

The restriction to at most two ligand-specific reactions is further justified in two ways additional to the exploratory forward–backward runs. First, the full saturated model, in which all ligand-specific fold-change parameters are estimated simultaneously (71 estimated parameters), is fitted directly. Comparing its maximised likelihood with that of the ranked models bounds the improvement that any additional ligand-specific reaction could possibly buy. It reaches a negative log-likelihood of 205.57, giving a gain of 14.35 AIC over the best ranked model and hence a bound of 14.35 *−* 2*k* AIC on any candidate with *k* additional ligand-specific fold-changes. Second, a forward step was constructed from the leading models after the model selection, in which 29 candidate models each add one further ligand-specific reaction to the three best models, each calibrated with the same protocol.

### Clustering

For each of the 9 retained models we collected the optimization histories of all calibrations and created ensembles of parameter vectors. The ensemble of a model consists of all recorded optimisation endpoints with 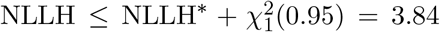 above the model’s own optimum, duplicates removed. The cutoff is the 95th percentile of the 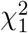 distribution, and it is applied per model, against that model’s own optimum, as in the uncertainty analysis above. Duplicate vectors are removed because identical parameter vectors produce identical trajectories and would only re-weight the clusters by how often the optimizer happened to record its incumbent, and ensembles are subsampled to at most 500 vectors, a limit that only M394 reaches. The number of clusters is chosen as the largest *k* for which no cluster holds less than 1% of the trajectories. We then simulated the model for each parameter vector and each experimental condition *c*, i.e. each ligand to obtain simulated observable trajectories, which were assembled into a 3-dimensional array 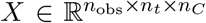, where *n*_obs_ = 2 is the number of observables, *n*_*t*_ = 6 the number of timepoints and *n*_*C*_ = 3 the number of ligands. To account for the different scaling factors *s*_*i*_ across ligands, trajectories were normalized separately for each experimental condition using an affine transformation based on the empirical minimum and maximum over all trajectories and time points in that condition, so that all variables contributed on a comparable scale to the distance computation [68].

Using the euclidean distance between trajectories we defined the pairwise distance matrix for a condition *c* as

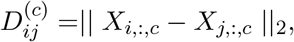

treating each trajectory as a point in 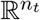 . Since the observable trajectories are aligned on a common time grid, we can then obtain a single overall dissimilarity matrix by averaging across conditions

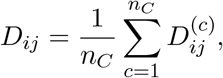

which captures similarity of temporal dynamics consistently across experimental conditions [68].

Clustering was performed on the distance matrix *D* using agglomerative hierarchical clustering with complete linkage, so the distance between two clusters is defined as the maximum pairwise distance between their members, which tends to produce compact, well-separated clusters.

### Implementation and availability

The model was formulated in the *SBML* format [69] and for specifying the parameter estimation problem we used *PEtab* (version 0.8.2) [48]. All code was executed with Python 3.12.10. The parameter estimation was done using the Python package *pyPESTO* (version 0.5.6) [58] with simulations carried out by *AMICI* (version 0.30.0) [70]. As optimizer choices we used enhanced scatter search [49] with inner optimization carried out by the Fides optimizer [59]. The model selection problem was built with the *PEtab Select* specification standard (version 0.4.0) [71]. Clustering was done using the Python package, *scikit-learn* (version 1.9.0) [72]. All SBML and PEtab files, and the instructions to reproduce the shown results, are available at https://doi.org/10.5281/zenodo.20035016 and on Github https://github.com/vwiela/Mathematical-Modeling-of-the-Canonical-Aryl-Hydrocarbon-Receptor-Pathway.git.

## Author contributions

Conceptualization, H.W., J.H. and I.F. Experiments, H.W., I.I., L.R.S. Methodology, V.W., T.B. Computations, V.W., T.B. Writing - Original Draft, V.W. and J.H. Writing - Review & Editing, all authors. Funding Acquisition, J.H and I.F.

