## Supplementary Information, Figures and Tables for "Mathematical Modeling of the Canonical Aryl Hydrocarbon Receptor Pathway in murine bone-marrow-derived macrophages"

**S2 Supplementary Figures** **22**

**S3 Supplementary Tables** **33**

### S1 Supplementary Information

#### S1.1 Measurement data description

The experimental design is 2 genes $\times$ 3 conditions $\times$ 6 time points $\times$ 3 replicates, = 108 values. The calibration dataset contains 105. The three values of BpA/*Cyp1a1*/3 h showed implausible values and discarded as failed measurements, leading to 105 measurements (Table S3). Additionally, two values are recorded as exactly zero: BpA, 24 h, replicate 3, for both genes. They are ordinary data points under the additive Gaussian error model but undefined under any log-scale likelihood, which is why the log-scale variants in the noise model comparison (Supplement S1.7) are fitted to 103 rather than 105 values.

Relative expression was quantified as $2^{-\Delta C_T}$ against $\beta$-actin for each sample, and the reported quantity is the ratio of the treated to the vehicle-control value. That ratio is formed replicate-wise: the $2^{-\Delta C_T}$ value of treated replicate $i$ is divided by the $2^{-\Delta C_T}$ value of control replicate $i$. Hence, each normalised measurement carries the full variance of a single control replicate rather than a reduced variance due to mean normalisation.

The vehicle control is not a constant reference (Figure S11a). For *Cyp1a1* the pooled within-time-point standard deviation is $1.17\,C_T$ and the geometric mean of the control drifts by $3.89\,C_T$ (14.8-fold) across the time course, between $7.50 \cdot 10^{-6}$ at 4 h and $5.07 \cdot 10^{-7}$ at 8 h. For *Ahrr* the corresponding values are $0.80\,C_T$ and $2.88\,C_T$ (7.4-fold), between $3.40 \cdot 10^{-3}$ at 4 h and $4.61 \cdot 10^{-4}$ at 3 h. Because the control sits in the denominator of every reported value, this is not separable from signal by inspection. Propagated into the normalised values, the control contributes a per-time-point relative standard deviation of 181/154/32/221/67/74 % for *Cyp1a1* and 81/161/21/83/4/55 % for *Ahrr* at 2, 3, 4, 6, 8 and 24 h. These are the values entering the explicit control-variance error models n4 and n7 of Section S1.7.

The ICZ $2^{-\Delta C_T}$ signal falls monotonically after its 4 h peak (Figure S11b). Between 6 h and 8 h the numerator falls 1.5-fold while the denominator falls 6.5-fold, while the mean of the replicate-wise ratios rises from 17.6 to 42.4. The two fold-changes are means of the replicates and the ratio is a mean of ratios, so the latter is not the quotient of the former. The apparent secondary peak at 8 h in the $\Delta\Delta C_T$ values is therefore an artefact of the normalisation, not a feature of the transcriptional response, and it is therefore not interpreted mechanistically in the manuscript.

#### S1.2 Model description

In the main manuscript we shortly described the canonical pathway. Here, we give a more detailed description including the 35 reactions together with its corresponding mass-action flux as listed in Table S4.

The model is divided into three compartments: the extracellular space, the cytoplasm and the nucleus, and the subscripts e, c, and n respectively designate the location of the species. All reactions are modeled with mass-action kinetics; we denote the reaction flux of reaction $R_i$ by $v_i$ and concentration of species $X$ by $[X]$.

Reaction fluxes are written below as intensive rates in the compartment to which the reaction is assigned. In the ODE system, the contribution of a reaction $\mathrm{R}_r$ located in a compartment of volume $V_r$ to the concentration of a species located in a compartment of volume $V_X$ is $(V_r/V_X)\,v_r$; for reactions whose participants share one compartment this factor equals one. The compartment volumes are $V_\mathrm{e} = 1\,\mathrm{L}$ (the medium, treated as a reservoir), $V_\mathrm{c} = 10^{-12}\,\mathrm{L}$ and $V_\mathrm{n} = 10^{-13}\,\mathrm{L}$. Four reactions connect compartments and therefore carry a volume factor: ligand exchange ($\mathrm{R}_2$, $\mathrm{R}_3$) and nuclear import ($\mathrm{R}_8$), both assigned to the cytoplasm, and translation of the nuclear *Cyp1a1* transcript into cytosolic protein ($\mathrm{R}_{21}$), assigned to the nucleus. The deposited SBML file declares the units of every compartment, species and parameter and passes the libSBML unit-consistency check with 0 errors and 0 warnings.

**Extracellular ligand dynamics and membrane exchange.** The only extracellular species is the ligand $\mathrm{L_e}$. We assumed that the cell encounters only one certain ligand, which in the case of the measured data is 3-methylcholanthrene (3MC), indolo[3,2-b]carbazole (ICZ) or the endocrine disruptor Bisphenol A (BpA) and no other endogenous or xenobiotic ligands. Furthermore, the ligand degradation does not generate other ligands of different potency. Extracellular ligand is removed by first-order degradation and can reversibly exchange with the cytoplasm via diffusion:

$$
\begin{aligned}
&(\mathrm{R}_1)\quad \mathrm{L_e} \to \varnothing, && v_1 = k_1\,[\mathrm{L_e}],\\
&(\mathrm{R}_2)\quad \mathrm{L_e} \to \mathrm{L_c}, && v_2 = k_2\,[\mathrm{L_e}],\\
&(\mathrm{R}_3)\quad \mathrm{L_c} \to \mathrm{L_e}, && v_3 = k_3\,[\mathrm{L_c}].
\end{aligned}
$$

In addition to enzyme-mediated ligand clearance (see below), we include a first-order cytosolic ligand degradation term with constant rate to capture unspecific clearance:

$$
(\mathrm{R}_4)\quad \mathrm{L_c} \to \varnothing, \qquad v_4 = k_4\,[\mathrm{L_c}].
$$

This unspecific degradation is condition-specific: $k_4$ is estimated for BpA and fixed to zero for 3MC and ICZ. With the compartment convention above, the ligand balances read

$$
\begin{aligned}
\frac{d[\mathrm{L_e}]}{dt} &= -v_1 - \frac{V_\mathrm{c}}{V_\mathrm{e}}\,(v_2 - v_3),\\
\frac{d[\mathrm{L_c}]}{dt} &= v_2 - v_3 - v_4 - v_7 + v_{-7} - v_{23},
\end{aligned}
$$

with the binding and Cyp1a1-mediated degradation fluxes $v_7$, $v_{-7}$ and $v_{23}$ defined below: the medium is depleted by the amount taken up by the cell, scaled by the volume ratio.

**Cytosolic AhR turnover and ligand binding.** Cytosolic AhR ($\mathrm{AhR_c}$) is constitutively produced and degraded. Ligand binding in the cytosol is modeled as a reversible mass-action process forming the activated complex $\mathrm{AhR_c{:}L_c}$:

$$\begin{aligned}
&(\mathrm{R}_5) && \varnothing \to \mathrm{AhR_c}, && v_5 = k_5,\\
&(\mathrm{R}_6) && \mathrm{AhR_c} \to \varnothing, && v_6 = k_6\,[\mathrm{AhR_c}],\\
&(\mathrm{R}_7) && \mathrm{AhR_c} + \mathrm{L_c} \to \mathrm{AhR_c{:}L_c}, && v_7 = k_7\,[\mathrm{AhR_c}]\,[\mathrm{L_c}],\\
&(\mathrm{R}_{-7}) && \mathrm{AhR_c{:}L_c} \to \mathrm{AhR_c} + \mathrm{L_c}, && v_{-7} = k_{-7}\,[\mathrm{AhR_c{:}L_c}].
\end{aligned}$$

We assume that aryl hydrocarbon receptor (AhR) as well as AhR nuclear translocator (ARNT) expression and degradation are not regulated by AhR signaling (even though AhR expression can be influenced by AhR signaling in some contexts). Additionally, the associated chaperones, which keep the unactivated AhR in the cytosol and enable non-genomic cross-talk to other pathways, are not modeled. These chaperones are not rate-limiting due to their high cellular abundance [1] and rapid dissociation of the AhR–chaperon complex upon nuclear entry [2].

**Nuclear import and nuclear AhR degradation.** Upon activation, the cytosolic complex is imported into the nucleus. We treat nuclear import as effectively non-reversible, consistent with rapid proteasomal degradation of AhR after nuclear export [3, 4]. Together with $\mathrm{R}_7/\mathrm{R}_{-7}$, this gives the two-stage mechanism commonly described for ligand-activated AhR: a fast, reversible cytosolic binding equilibrium ($\mathrm{R}_7/\mathrm{R}_{-7}$) followed by an effectively irreversible transformation and import step ($\mathrm{R}_8$). In the model, this is captured by (i) import and (ii) nuclear degradation of the ligand-bound nuclear AhR complex:

$$\begin{aligned}
&(\mathrm{R}_8) && \mathrm{AhR_c{:}L_c} \to \mathrm{AhR_n{:}L_n}, && v_8 = k_8\,[\mathrm{AhR_c{:}L_c}],\\
&(\mathrm{R}_9) && \mathrm{AhR_n{:}L_n} \to \varnothing, && v_9 = k_9\,[\mathrm{AhR_n{:}L_n}].
\end{aligned}$$

Because the import reaction is assigned to the cytoplasm while its product is nuclear, it enters the nuclear balance as $(V_\mathrm{c}/V_\mathrm{n})\,v_8$. Reaction ($\mathrm{R}_9$) represents degradation of nuclear AhR, effectively removing the bound ligand from the system.

**ARNT turnover and heterodimerization with nuclear AhR.** AhR's transcriptional co-activator in the nucleus ($\mathrm{ARNT_n}$) is also produced and degraded constitutively and reversibly forms a heterodimer with ligand-bound nuclear AhR:

$$\begin{aligned}
&(\mathrm{R}_{10}) && \varnothing \to \mathrm{ARNT_n}, && v_{10} = k_{10},\\
&(\mathrm{R}_{11}) && \mathrm{ARNT_n} \to \varnothing, && v_{11} = k_{11}\,[\mathrm{ARNT_n}],\\
&(\mathrm{R}_{12}) && \mathrm{AhR_n{:}L_n} + \mathrm{ARNT_n} \to \mathrm{AhR_n{:}L_n{:}ARNT_n}, && v_{12} = k_{12}\,[\mathrm{AhR_n{:}L_n}]\,[\mathrm{ARNT_n}],\\
&(\mathrm{R}_{-12}) && \mathrm{AhR_n{:}L_n{:}ARNT_n} \to \mathrm{AhR_n{:}L_n} + \mathrm{ARNT_n}, && v_{-12} = k_{-12}\,[\mathrm{AhR_n{:}L_n{:}ARNT_n}].
\end{aligned}$$

**Promoter binding of the activator complex.** The transcriptional activator complex $\mathrm{AhR_n{:}L_n{:}ARNT_n}$ can bind target gene promoters. Although multiple aryl hydrocarbon response elements (AhREs) may exist in a given promoter [5–7], we model promoter binding as a single reversible binding event per promoter. Binding to the AhRR promoter $\mathrm{p_{Ahrr}}$ and to the Cyp1a1 promoter $\mathrm{p_{Cyp1a1}}$ is described by:

$$\begin{aligned}
&(\mathrm{R}_{13}) && \mathrm{p_{Ahrr} + AhR_n{:}L_n{:}ARNT_n \rightarrow AhR_n{:}L_n{:}ARNT_n{:}p_{Ahrr}}, && v_{13} = k_{13}\,[\mathrm{p_{Ahrr}}]\,[\mathrm{AhR_n{:}L_n{:}ARNT_n}],\\
&(\mathrm{R}_{-13}) && \mathrm{AhR_n{:}L_n{:}ARNT_n{:}p_{Ahrr} \rightarrow p_{Ahrr} + AhR_n{:}L_n{:}ARNT_n}, && v_{-13} = k_{-13}\,[\mathrm{AhR_n{:}L_n{:}ARNT_n{:}p_{Ahrr}}],\\
&(\mathrm{R}_{14}) && \mathrm{p_{Cyp1a1} + AhR_n{:}L_n{:}ARNT_n \rightarrow AhR_n{:}L_n{:}ARNT_n{:}p_{Cyp1a1}}, && v_{14} = k_{14}\,[\mathrm{p_{Cyp1a1}}]\,[\mathrm{AhR_n{:}L_n{:}ARNT_n}],\\
&(\mathrm{R}_{-14}) && \mathrm{AhR_n{:}L_n{:}ARNT_n{:}p_{Cyp1a1} \rightarrow p_{Cyp1a1} + AhR_n{:}L_n{:}ARNT_n}, && v_{-14} = k_{-14}\,[\mathrm{AhR_n{:}L_n{:}ARNT_n{:}p_{Cyp1a1}}].
\end{aligned}$$

**Transcription, basal transcription, and mRNA degradation.** Transcription is modeled as synthesis of mRNA catalyzed by the corresponding promoter-bound activator complex (which is not consumed). Additionally, we include basal (activator-independent) transcription from free promoter:

$$\begin{aligned}
&(\mathrm{R}_{15}) && \mathrm{AhR_n{:}L_n{:}ARNT_n{:}p_{Ahrr} \rightarrow AhR_n{:}L_n{:}ARNT_n{:}p_{Ahrr} + mRNA_{AhRR}},\\
& && \qquad v_{15} = k_{15}\,[\mathrm{AhR_n{:}L_n{:}ARNT_n{:}p_{Ahrr}}],\\
&(\mathrm{R}_{16}) && \emptyset \rightarrow \mathrm{mRNA_{AhRR}}, && v_{16} = k_{16},\\
&(\mathrm{R}_{17}) && \mathrm{mRNA_{AhRR}} \rightarrow \varnothing, && v_{17} = k_{17}\,[\mathrm{mRNA_{AhRR}}],\\
&(\mathrm{R}_{18}) && \mathrm{AhR_n{:}L_n{:}ARNT_n{:}p_{Cyp1a1} \rightarrow AhR_n{:}L_n{:}ARNT_n{:}p_{Cyp1a1} + mRNA_{Cyp1a1}},\\
& && \qquad v_{18} = k_{18}\,[\mathrm{AhR_n{:}L_n{:}ARNT_n{:}p_{Cyp1a1}}],\\
&(\mathrm{R}_{19}) && \emptyset \rightarrow \mathrm{mRNA_{Cyp1a1}}, && v_{19} = k_{19},\\
&(\mathrm{R}_{20}) && \mathrm{mRNA_{Cyp1a1}} \rightarrow \varnothing, && v_{20} = k_{20}\,[\mathrm{mRNA_{Cyp1a1}}].
\end{aligned}$$

We do not account for epigenetic regulation of chromatin accessibility or transcript truncation/isoforms and model mRNA removal as first-order degradation. The transcription rate of each gene is the sum of its induced and its basal flux, $v_{15} + v_{16}$ for *Ahrr* and $v_{18} + v_{19}$ for *Cyp1a1*. Basal transcription ($\mathrm{R}_{16}$, $\mathrm{R}_{19}$) is a constant, activator-independent flux; consequently, occupancy of a promoter by the repressor complex (see below) suppresses the induced component of transcription but not the basal component.

**Translation, Cyp1a1-mediated ligand degradation, and Cyp1a1 turnover.** Cyp1a1 protein is translated from $\mathrm{mRNA_{Cyp1a1}}$ and degraded constitutively:

$$\begin{aligned}
&(\mathrm{R}_{21}) && \mathrm{mRNA_{Cyp1a1} \rightarrow mRNA_{Cyp1a1} + Cyp1a1_c}, && v_{21} = k_{21}\,[\mathrm{mRNA_{Cyp1a1}}],\\
&(\mathrm{R}_{22}) && \mathrm{Cyp1a1_c} \rightarrow \varnothing, && v_{22} = k_{22}\,[\mathrm{Cyp1a1_c}].
\end{aligned}$$

Since the transcript is nuclear and the protein cytosolic, translation enters the cytosolic balance as $(V_\mathrm{n}/V_\mathrm{c})\,v_{21}$. Cyp1a1 provides negative feedback by degrading cytosolic ligand. Although this is an enzyme-mediated process, we use a mass-action rate (instead of Michaelis–Menten kinetics) to simplify the mathematical complexity and support efficient gradient calculations:

$$(\mathrm{R}_{23}) \quad \mathrm{L_c + Cyp1a1_c \rightarrow Cyp1a1_c}, \qquad v_{23} = k_{23}\,[\mathrm{L_c}]\,[\mathrm{Cyp1a1_c}].$$

This route is condition-specific: it is active for 3MC and ICZ and switched off for BpA, whose cytosolic clearance is carried by $k_4$ instead (see the $R_4$ paragraph above). We assume the extracellular ligand $L_e$ is not degraded by cytosolic Cyp1a1.

**AhRR feedback: translation, turnover, ARNT sequestration, and promoter repression.** The mRNA of AhRR is translated to nuclear AhRR protein $AhRR_n$, which is degraded constitutively:

$$
\begin{aligned}
&(R_{24}) \quad \mathrm{mRNA_{AhRR}} \to \mathrm{mRNA_{AhRR}} + \mathrm{AhRR_n}, && v_{24} = k_{24}\,[\mathrm{mRNA_{AhRR}}],\\
&(R_{25}) \quad \mathrm{AhRR_n} \to \varnothing, && v_{25} = k_{25}\,[\mathrm{AhRR_n}].
\end{aligned}
$$

AhRR inhibits signaling by sequestering ARNT through reversible heterodimerization:

$$
\begin{aligned}
&(R_{26}) \quad \mathrm{AhRR_n} + \mathrm{ARNT_n} \to \mathrm{AhRR_n{:}ARNT_n}, && v_{26} = k_{26}\,[\mathrm{AhRR_n}]\,[\mathrm{ARNT_n}],\\
&(R_{-26}) \quad \mathrm{AhRR_n{:}ARNT_n} \to \mathrm{AhRR_n} + \mathrm{ARNT_n}, && v_{-26} = k_{-26}\,[\mathrm{AhRR_n{:}ARNT_n}].
\end{aligned}
$$

The resulting repressor complex can bind free promoter regions and thereby prevent activation-dependent transcription. Promoter repression and derepression are modeled as reversible binding events:

$$
\begin{aligned}
&(R_{27}) \quad \mathrm{p_{Ahrr}} + \mathrm{AhRR_n{:}ARNT_n} \to \mathrm{AhRR_n{:}ARNT_n{:}p_{Ahrr}}, && v_{27} = k_{27}\,[\mathrm{p_{Ahrr}}]\,[\mathrm{AhRR_n{:}ARNT_n}],\\
&(R_{-27}) \quad \mathrm{AhRR_n{:}ARNT_n{:}p_{Ahrr}} \to \mathrm{p_{Ahrr}} + \mathrm{AhRR_n{:}ARNT_n}, && v_{-27} = k_{-27}\,[\mathrm{AhRR_n{:}ARNT_n{:}p_{Ahrr}}],\\
&(R_{28}) \quad \mathrm{p_{Cyp1a1}} + \mathrm{AhRR_n{:}ARNT_n} \to \mathrm{AhRR_n{:}ARNT_n{:}p_{Cyp1a1}}, && v_{28} = k_{28}\,[\mathrm{p_{Cyp1a1}}]\,[\mathrm{AhRR_n{:}ARNT_n}],\\
&(R_{-28}) \quad \mathrm{AhRR_n{:}ARNT_n{:}p_{Cyp1a1}} \to \mathrm{p_{Cyp1a1}} + \mathrm{AhRR_n{:}ARNT_n}, && v_{-28} = k_{-28}\,[\mathrm{AhRR_n{:}ARNT_n{:}p_{Cyp1a1}}].
\end{aligned}
$$

In contrast to the promoter–activator complexes $\mathrm{AhR_n{:}L_n{:}ARNT_n{:}p_{Ahrr}}$ and $\mathrm{AhR_n{:}L_n{:}ARNT_n{:}p_{Cyp1a1}}$ (which drive transcription via $v_{15}$ and $v_{18}$), the promoter–repressor complexes in ($R_{27}$) and ($R_{28}$) do not induce transcription in the model.

Finally, we assume the modeled cell does not grow, divide, or die over the simulated time window. Cytoplasmic and nuclear volumes are constant, the total DNA amount is conserved, and concentrations are not affected by asymmetric division.

#### Observation and noise function

We measured the concentrations of two biochemical species, i.e. $\mathrm{mRNA_{Cyp1a1}}$ and $\mathrm{mRNA_{AhRR}}$. Due to the nature of the data being relative qPCR-measurements, we used a scaling factor to map the measurements to the real concentrations of the species. This leads to the following observables

$$
\begin{aligned}
y_1(t,\theta) &= s_{\mathrm{mRNA_{Cyp1a1}}} \cdot \mathrm{mRNA_{Cyp1a1}}(t)\\
y_2(t,\theta) &= s_{\mathrm{mRNA_{AhRR}}} \cdot \mathrm{mRNA_{AhRR}}(t).
\end{aligned}
$$

The heteroskedastic noise model introduces gaussian noise for each observable that accounts for a fixed noise part and for increasing measurement error with increasing data amplitude, $\sigma_i = \sigma_{0,i} + \sigma_{1,i}\, y_i$. The adequacy of this choice is assessed in Section S1.7, where it is compared against six alternatives, seven error models in total on a common AIC scale with randomized quantile-residual diagnostics.

##### Model parameterization

To achieve a more efficient parameterization of the model and enhance optimization performance, we followed the recommendations in [8]. For better numerical stability during optimization, we parameterized all association-dissociation reactions

$$\mathrm{R}_7/\mathrm{R}_{-7}; \mathrm{R}_{12}/\mathrm{R}_{-12}; \mathrm{R}_{13}/\mathrm{R}_{-13}; \mathrm{R}_{14}/\mathrm{R}_{-14}; \mathrm{R}_{26}/\mathrm{R}_{-26}; \mathrm{R}_{27}/\mathrm{R}_{-27}; \mathrm{R}_{28}/\mathrm{R}_{-28}$$

using, besides the association rate constant $k_i$, an equilibrium constant $d_i$ as the estimated parameter. The rate constant of the dissociation reaction is the ratio of the two, for example

$$k_{-7} := \frac{k_7}{d_7},$$

so that $d_i = k_i/k_{-i}$ is an *association* constant with unit $\mathrm{nM}^{-1}$; the conventional dissociation constant is $K_{D,i} = 1/d_i$ in nM. Additionally, we removed the scaling invariance inherent to the Cyp1a1 mediated feedback loop in the model by reparameterizing the transcription of $\mathrm{mRNA}_{\mathrm{Cyp1a1}}$. For this we fix the basal transcription rate to $k_{19} = 1.0\,\mathrm{nM\,s^{-1}}$; the induced transcription rate $k_{18}$ is then expressed relative to this basal flux and carries the unit $\mathrm{nM}^{-1}$ (Table S5).

The modeled cell is a somatic diploid cell carrying two copies of each target gene promoter in a nuclear volume of $V_\mathrm{n} = 10^{-13}\,\mathrm{L}$. We therefore fix the total promoter concentrations to $\mathrm{total}_{\mathrm{pCyp1a1}} = \mathrm{total}_{\mathrm{pAhrr}} = 2/(N_A V_\mathrm{n}) = 3.32 \cdot 10^{-2}\,\mathrm{nM}$.

In total, the reaction rates involve 35 parameters, of which 34 are estimated ($k_{19}$ is fixed; $k_4$ is estimated for BpA only). Additionally, we estimate 2 scaling parameters and 4 noise parameters, giving 40 parameters shared by all candidate models of the model selection; each candidate model adds the ligand-specific fold-change parameters of its subspace (Table S6). To improve numerical convergence, estimation is carried out on $\log_{10}$-scale and parameters are equipped with a uniform prior on that scale.

All model parameters, including steady-state constants, scaling and noise parameters, with corresponding bounds, units and estimation status are given in Table S5 below.

#### S1.3 Steady-state calculations

We assume that all states involving the ligand or a ligand-bound complex have an initial concentration of zero. At steady state, the concentrations of all states are constant; species are neither produced nor degraded. To compute steady-state concentrations, we therefore set all time derivatives to zero and solve the resulting algebraic equations.

**Cyp1a1-mRNA and protein.** We begin with the mRNA of the target gene *Cyp1a1*. Setting the rate of change to zero gives

$$\frac{d([\mathrm{mRNA}_{\mathrm{Cyp1a1}}])}{dt} = \big(k_{18}\,[\mathrm{AhR_n{:}L_n{:}ARNT_n{:}p_{Cyp1a1}}] + k_{19}\big) - \big(k_{20}\,[\mathrm{mRNA}_{\mathrm{Cyp1a1}}]\big).$$

Solving for the mRNA concentration yields

$$[\mathrm{mRNA}_{\mathrm{Cyp1a1}}] = \frac{k_{19}}{k_{20}}. \tag{1}$$

Next, we use (1) in the steady-state balance to compute cytosolic Cyp1a1 protein,

$$\frac{d([\mathrm{Cyp1a1_c}])}{dt} = -k_{22}\,[\mathrm{Cyp1a1_c}] + \frac{V_\mathrm{n}}{V_\mathrm{c}}\,k_{21}\,[\mathrm{mRNA}_{\mathrm{Cyp1a1}}],$$

which directly yields

$$[\mathrm{Cyp1a1_c}] = \frac{V_\mathrm{n}}{V_\mathrm{c}} \cdot \frac{k_{21}\,k_{19}}{k_{22}\,k_{20}}, \tag{2}$$

where the volume ratio arises because the transcript is nuclear and the protein cytosolic (Section S1.2).

**AhRR mRNA and cytosolic AhR.** For mRNA of *ahrr*, the steady-state condition applied to

$$\frac{d([\mathrm{mRNA}_{\mathrm{AhRR}}])}{dt} = \big(k_{15}\,[\mathrm{AhR_n{:}L_n{:}ARNT_n{:}p_{Ahrr}}] + k_{16}\big) - k_{17}\,[\mathrm{mRNA}_{\mathrm{AhRR}}]$$

gives

$$[\mathrm{mRNA}_{\mathrm{AhRR}}] = \frac{k_{16}}{k_{17}}. \tag{3}$$

The cytosolic AhR balance simplifies, because at steady state the ligand-binding fluxes cancel:

$$\frac{d([\mathrm{AhR_c}])}{dt} = k_5 - k_6\,[\mathrm{AhR_c}] - (k_7\,[\mathrm{L_c}]\,[\mathrm{AhR_c}] - \frac{k_7}{d_7}\,[\mathrm{AhR_c{:}L}]),$$

which reduces to

$$[\mathrm{AhR_c}] = \frac{k_5}{k_6} \tag{4}$$

**Promoter binding states (Cyp1a1 and AhRR).** We now analyse the binding equilibria at the two promoters. For the *Cyp1a1* promoter, the steady-state condition on

$$\frac{d([\mathrm{p_{Cyp1a1}}])}{dt} = -\left(k_{14}\,[\mathrm{AhR_n{:}L_n{:}ARNT_n}]\,[\mathrm{p_{Cyp1a1}}] - \frac{k_{14}}{d_{14}}\,[\mathrm{AhR_n{:}L_n{:}ARNT_n{:}p_{Cyp1a1}}]\right) - \left(k_{28}\,[\mathrm{AhRR_n{:}ARNT}]\,[\mathrm{p_{Cyp1a1}}] - \frac{k_{28}}{d_{28}}\,[\mathrm{AhRR_n{:}ARNT{:}p_{Cyp1a1}}]\right),$$

gives the equilibrium relation

$$[\mathrm{p_{Cyp1a1}}] = \frac{[\mathrm{AhRR_n{:}ARNT{:}p_{Cyp1a1}}]}{d_{28}\,[\mathrm{AhRR_n{:}ARNT}]}. \tag{5}$$

Repeating the same procedure for the *ahrr* promoter yields

$$[\mathrm{p_{Ahrr}}] = \frac{[\mathrm{AhRR_n{:}ARNT{:}p_{Ahrr}}]}{d_{27}\,[\mathrm{AhRR_n{:}ARNT}]}. \tag{6}$$

**Free nuclear ARNT and the AhRR:ARNT complex.** Next we consider nuclear ARNT. Setting the derivative to zero in

$$\begin{aligned}\frac{d([\mathrm{ARNT_n}])}{dt} = {}& k_{10} - k_{11}\,[\mathrm{ARNT_n}] \\ & - \left(k_{12}\,[\mathrm{AhR_n{:}L}]\,[\mathrm{ARNT_n}] - \frac{k_{12}}{d_{12}}\,[\mathrm{AhR_n{:}L_n{:}ARNT_n}]\right) \\ & - \left(k_{26}\,[\mathrm{AhRR_n}]\,[\mathrm{ARNT_n}] - \frac{k_{26}}{d_{26}}\,[\mathrm{AhRR_n{:}ARNT}]\right)\end{aligned}$$

leads to the expression

$$[\mathrm{AhRR_n{:}ARNT}] = \frac{-k_{10} + k_{11}\,[\mathrm{ARNT_n}] + k_{26}\,[\mathrm{AhRR_n}]\,[\mathrm{ARNT_n}]}{\frac{k_{26}}{d_{26}}}. \tag{7}$$

To proceed, we substitute (3) into the steady-state balance for nuclear AhRR,

$$\begin{aligned}\frac{d([\mathrm{AhRR_n}])}{dt} &= -\left(k_{26}\,[\mathrm{AhRR_n}]\,[\mathrm{ARNT_n}] - \frac{k_{26}}{d_{26}}\,[\mathrm{AhRR_n{:}ARNT}]\right) \\ &\quad + k_{24}\,[\mathrm{mRNA_{AhRR}}] - k_{25}\,[\mathrm{AhRR_n}] \\ &= -k_{10} + k_{11}\,[\mathrm{ARNT_n}] + k_{24}\,\frac{k_{16}}{k_{17}} - k_{25}\,[\mathrm{AhRR_n}],\end{aligned}$$

which simplifies to

$$[\mathrm{AhRR_n}] = \frac{-k_{10} + k_{11}\,[\mathrm{ARNT_n}] + k_{24}\,\dfrac{k_{16}}{k_{17}}}{k_{25}}. \tag{8}$$

We now use the balance for [$\mathrm{AhRR_n{:}ARNT}$] together with the promoter expressions (5) and (6) to eliminate the remaining terms. The resulting equation,

$$\begin{aligned}\frac{d([\mathrm{AhRR_n{:}ARNT}])}{dt} &= \left(k_{26}\,[\mathrm{AhRR_n}]\,[\mathrm{ARNT_n}] - \frac{k_{26}}{d_{26}}\,[\mathrm{AhRR_n{:}ARNT}]\right)\\ &\quad - \left(k_{27}\,[\mathrm{AhRR_n{:}ARNT}]\,[\mathrm{p_{Ahrr}}] - \frac{k_{27}}{d_{27}}\,[\mathrm{AhRR_n{:}ARNT{:}p_{Ahrr}}]\right)\\ &\quad - \left(k_{28}\,[\mathrm{AhRR_n{:}ARNT}]\,[\mathrm{p_{Cyp1a1}}] - \frac{k_{28}}{d_{28}}\,[\mathrm{AhRR_n{:}ARNT{:}p_{Cyp1a1}}]\right)\\ &= k_{10} - k_{11}\,[\mathrm{ARNT_n}],\end{aligned}$$

immediately yields

$$[\mathrm{ARNT_n}] = \frac{k_{10}}{k_{11}} \tag{9}$$

Inserting (9) into (8) gives a compact expression for nuclear AhRR:

$$[\mathrm{AhRR_n}] = \frac{-k_{10} + k_{11}\cdot\dfrac{k_{10}}{k_{11}} + k_{24}\,\dfrac{k_{16}}{k_{17}}}{k_{25}} = \frac{k_{24}\,k_{16}}{k_{25}\,k_{17}}. \tag{10}$$

Plugging (9) and (10) into (7) finally gives

$$[\mathrm{AhRR_n{:}ARNT}] = \frac{d_{26}\,k_{24}\,k_{16}\,k_{10}}{k_{25}\,k_{17}\,k_{11}} \tag{11}$$

**Cyp1a1 promoter conservation and occupancy.** The expression (5) determines $\mathrm{p_{Cyp1a1}}$ in terms of the complex [$\mathrm{AhRR_n{:}ARNT{:}p_{Cyp1a1}}$], but the latter is still unknown. Let $\mathrm{total_{p_{Cyp1a1}}}$ denote the total concentration of the promoter, which is assumed to be initially unbound. As the promoter can be unbound or bound to either complex $\mathrm{AhRR_n{:}ARNT}$ or $\mathrm{AhR_n{:}L_n{:}ARNT_n}$, but is neither produced nor degraded, the total concentration of all species with $\mathrm{p_{Cyp1a1}}$ is constant. Using

$$\mathrm{total_{p_{Cyp1a1}}} = [\mathrm{p_{Cyp1a1}}] + [\mathrm{AhR_n{:}L_n{:}ARNT_n{:}p_{Cyp1a1}}] + [\mathrm{AhRR_n{:}ARNT{:}p_{Cyp1a1}}]$$

we can solve for the unbound promoter:

$$[\mathrm{p_{Cyp1a1}}] = \mathrm{total_{p_{Cyp1a1}}} - [\mathrm{AhRR_n{:}ARNT{:}p_{Cyp1a1}}]. \tag{12}$$

Now we compute the bound state using the steady-state condition on

$$\begin{aligned}\frac{d([\mathrm{AhRR_n{:}ARNT{:}p_{Cyp1a1}}])}{dt} = \big(&k_{28}\,[\mathrm{AhRR_n{:}ARNT}]\,[\mathrm{p_{Cyp1a1}}]\\ &- \frac{k_{28}}{d_{28}}\,[\mathrm{AhRR_n{:}ARNT{:}p_{Cyp1a1}}]\big),\end{aligned}$$

together with (11). This gives

$$[\mathrm{AhRR_n{:}ARNT{:}p_{Cyp1a1}}] = \frac{k_{28}\,[\mathrm{AhRR_n{:}ARNT}]\,\mathrm{total}_{\mathrm{p_{Cyp1a1}}}}{k_{28}\,[\mathrm{AhRR_n{:}ARNT}] + \frac{k_{28}}{d_{28}}}. \tag{13}$$

Substituting (13) into (12) yields the final expression for the unbound promoter:

$$[\mathrm{p_{Cyp1a1}}] = \mathrm{total}_{\mathrm{p_{Cyp1a1}}} - \frac{k_{28}\,[\mathrm{AhRR_n{:}ARNT}]\,\mathrm{total}_{\mathrm{p_{Cyp1a1}}}}{k_{28}\,[\mathrm{AhRR_n{:}ARNT}] + \frac{k_{28}}{d_{28}}}. \tag{14}$$

The same derivations apply to the *ahrr* promoter using its corresponding reactions.

#### S1.4 Existence, uniqueness and non-negativity of the ODE solutions

Denoting the species of the reaction network by $X_1, ..., X_n$ and reactions from Table S4 by

$$y_r \to y'_r,$$

where $y_r, y'_r \in \mathbb{N}_0^n$ are the reactant/product complexes and $k_r \geq 0$ are rate constants for reaction $r$. Under mass-action kinetics

$$v_r(x) = k_r x^{y_r} = k_r \prod_{i=1}^{n} x_i^{(y_r)_i},$$

the ODE is given by

$$\dot{x} = f(x) = \sum_{r=1}^{R} (y'_r - y_r) v_r(x),$$

i.e. the species formation rate functions is a polynomial vector field.

**Proposition 1.** *Let $x(t) \in \mathbb{R}^n$ denote the concentrations of all species appearing in the reaction network. Assume $k_r \geq 0\,\forall r = 1, ..., R$ and that each reaction follows mass-action kinetics. Then:*

1. ***(Existence/Uniqueness)*** *For every initial condition $x(0) = x_0 \in \mathbb{R}^n$ there exists a unique maximal solution $x : [0, T_{max}) \to \mathbb{R}^n$.*
2. ***(Non-negativity)*** *If $x_0 \in \mathbb{R}^n_{\geq 0}$, then $x(t) \in \mathbb{R}^n_{\geq 0}$ for all $t \in [0, T_{max})$.*
3. ***(Global existence)*** *For the network given by the reactions in Table S4, $T_{max} = \infty$, i.e. solutions exist for all $t \geq 0$.*

**Sketch of proof:**

*1. Local existence/uniqueness and 2. non-negativity:* Under mass-action kinetics local well-posedness and non-negativity are standard properties of the induced chemical reaction network (CRN) ODE. In particular, this is given since the resulting ODE has a polynomial right-hand side, so $f$ is locally Lipschitz. Therefore, the ODE admits a unique local solution according to the Picard–Lindelöf theorem. Moreover, the non-negative orthant $\mathbb{R}^{n_x}_{\geq 0}$ is forward invariant, i.e. if $x_0 \geq 0$, then $x(t) \geq 0$ for all times in the interval of existence.

*3. Global existence:* Define $V(x) = \sum_{i=1}^{n} V_i x_i$, the total amount of material, where $V_i$ denotes the volume of the compartment of species $X_i$ (with concentrations instead of amounts, transport between compartments of different volume would not conserve the sum). For the reactions from Table S4 we observe:

- The zero-order production terms are of the form $\emptyset \rightarrow$, contributing a constant $\leq a$.
- The unimolecular reactions contribute at most linearly in $V$, since each term is bounded by $k_r V_j x_j \leq k_r V$.
- The transport reactions $\mathrm{R}_2$, $\mathrm{R}_3$ and $\mathrm{R}_8$ move amount between compartments and contribute exactly zero to $\dot{V}$.
- The bimolecular reactions are associations or catalytic degradation, which do not increase $V$; they contribute $\leq 0$ to $\dot{V}$.

Hence there exist constants $a, b \geq 0$ such that for all $x \geq 0$,

$$\dot{V}(t) \leq a + bV(t).$$

By Grönwall's inequality, $V(t)$ is uniformly bounded and therefore the local solution extends uniquely for all $t \geq 0$, i.e. $T_{\max} = \infty$. □

A detailed proof of the first two properties for general mass-action kinetic derived ODEs is given in Chellaboina *et al.* [9].

#### S1.5 Simulation and optimization settings

For the parameter optimization of the pathway model we used a self-adaptive cooperative enhanced scatter search algorithm combined with local, gradient-based optimization [10] with an interior trust region reflective (Fides) [11] as the inner optimizer as implemented in the Python package *pyPESTO* [12]. For all calibrated models the following tolerances for the outer and inner optimizer were used

- Outer Sacess walltime: 8 hours
- Inner Fides maximal iterations: 1000
- Inner Fides absolute tolerance: $10^{-8}$
- Inner Fides relative tolerance: $10^{-8}$
- Hessian update for Fides: L-BFGS

and the following tolerances were used for SUNDIALS *CVODES* solver accessed via *AMICI*

- Maximal number of integration steps: 10.000
- Relative Tolerance: $10^{-12}$
- Absolute Tolerance: $10^{-8}$

**Search protocol.** Every calibration ran the same protocol per problem: 20 Fides multistarts, warm-started from the best available parameter vector, followed by an 8 h saCeSS global search on 10 workers whose best candidates were locally refined. The multistart optimum is always included as a refinement start, so the reported value can never be worse than the multistart alone. For the model selection, each of the 544 subspaces was calibrated as one task of a Slurm array (10 CPU cores and 64 GB per task) and the median wall time was 8.4 h. Software versions: Python 3.12.10, AMICI 0.30.0, pyPESTO 0.5.6, PEtab 0.8.2, PEtab Select 0.4.0, Fides 0.8.0, scikit-learn 1.9.0. The complete calibration record (per-fit objective value, number of converged starts and at-bound counts) is deposited with the code. The construction of the likelihood-based parameter ensembles used for the prediction uncertainty of Figure S2 and for the trajectory clustering is given in Section S1.11.

#### S1.6 Parameter bounds: literature audit

The 37 numeric bound pairs of the model (rate constants, scaling and ligand fold-change parameters) were audited against available literature. Where a source could be found, the bound pair was set from it and the was recorded. Where none could be found, the bound was set from an order-of-magnitude physical argument. Thirty-three of the 37 bound pairs are set from the audit, four keep an order-of-magnitude default. The bounds used throughout are those of Table S5. Table S2 reports the audit itself.

Comparing the parameter bounds with the estimated parameters of the best model we observe that (M343, Section S1.9) twelve of its 41 estimates lie within 0.01 decades of a bound (Figure S7) and the literatue helps to constraint the search space. All of them are profiled, as is every other estimated parameter of M343 (Table S17, Figures S8 and S9). In every case the profile rises past the 95 % threshold on the side away from the bound, so each of these parameters is bounded by the data from one side and by the literature constraints from the other. The converse case appears as well. The *Cyp1a1* induction strength $k_{18}$ is interior in M343 and identifiable: $2.44 \times 10^6$ with the profile interval $[2.59 \times 10^4, 4.23 \times 10^7]$, 3.21 decades wide and inside its bounds $[1, 10^9]$, so the upper bound derived above is not binding. The bound used on the extracellular ligand degradation rate $k_1$ is $[10^{-7}, 10^{-3}]\,\mathrm{s}^{-1}$, is deliberately wide, since a narrow window on this parameter would fix the medium half-life by construction instead of estimating it. Inside that range the estimate drifts to the slow end. The medium is represented as a 1 L reservoir, so this parameter is the model's only route of ligand loss and must absorb abiotic decay, adsorption to plastic, protein binding and cellular uptake together. Over a 24 h experiment, half-lives of 6, 60 and 600 days are indistinguishable. The profile of $k_1$ is one-sided: the estimate is $7.24 \times 10^{-5}\,\mathrm{s}^{-1}$ and the interval is $[1.31 \times 10^{-5}$, at bound], so the data do place a lower limit of $1.31 \cdot 10^{-5}\,\mathrm{s}^{-1}$ on the rate while in the other direction the profile reaches the box bound $10^{-3}\,\mathrm{s}^{-1}$ (a half-life of 12 min) without ever rising by 1.92. Faster ligand loss is therefore not excluded by the data, but by the box constraint since this parameter hard to be measured by a 24 h experiment. The ligand-specific partner is $k_4$ (`BpA_const_L_degrad`, $R_4$): the BpA condition does not scale $k_1$ by a fold-change but carries its own ligand-degradation constant, estimated for BpA and fixed elsewhere, so the profile of $k_4$ is the profile of that combination. It is one-sided in the same direction, $[4.40 \times 10^{-4}$, at bound] around $0.00259\,\mathrm{s}^{-1}$, with the upper side at the box bound $10^{-2}\,\mathrm{s}^{-1}$.

#### S1.7 Noise-model comparison and residual diagnostics

The observation model assumes additive Gaussian noise whose standard deviation is affine in the predicted value. Because the vehicle control varies substantially across the time course (Section S1.1), the question whether the error model should absorb the control variability explicitly was treated as a model-selection question. We fitted seven observation models to the same measurements with the same model, the same bounds and the same optimisation protocol

- n1: Gaussian, $\sigma = \sigma_0 + \sigma_1\mu$ (the chosen form)
- n2: Gaussian, $\sigma = \sqrt{\sigma_0^2 + \sigma_1^2\mu}$ (independent sources adding in variance, linear in the mean)
- n3: Gaussian, $\sigma = \sqrt{\sigma_0^2 + \sigma_1^2\mu^2}$ (constant coefficient of variation at high signal)
- n4: n2 plus an explicitly propagated, time-point-specific vehicle-control variance
- n5: Laplace, $b = \sqrt{b_0^2 + b_1^2\mu}$ (heavy tails)
- n6: log-normal
- n7: n6 plus the propagated control variance.

The log-scale variants n6 and n7 cannot accommodate the two measurements recorded as exactly zero (Section S1.1), and models fitted to different data are not AIC-comparable, so the primary comparison uses the 103 strictly positive measurements for all seven variants. Moreover, for a log-scale observable the optimiser reports a density over $\log_{10} y$, not over $y$, so we converted it using a Jacobian correction.

Table S7 lists the comparison on the topology of model M360. The adopted form n1 ranks first, by a difference of 34.7 in AIC to the next noise model family and 456 to the log-normal family. Because the choice could depend on the model topology, all seven noise model variants were fitted on four models: M360 (42 free parameters), M117 and M64 (44 free parameters, almost disjoint reaction sets) and the full saturated model (71 free parameters). The set of variants being indistinguishable by AIC is $\{\mathrm{n1}, \mathrm{n3}\}$ (Table S8), and n1 is chosen because it wins three of the four model topologies. Adding the measured DMSO variance to the likelihood makes the fit worse in every topology.

We further compared the residual diagnostics for the seven noise model variants. Standardised residuals $(y-\mu)/\sigma$ are $N(0,1)$ only under a Gaussian model and therefore cannot compare a Gaussian against a Laplace against a log-normal. We use randomized quantile residuals, $r_i = \Phi^{-1}(F_i(y_i))$ with $F_i$ the fitted predictive distribution, which are exactly $N(0,1)$ for any correctly specified model [13]. Moreover, residuals are computed per replicate and timepoint

$$r\_ik = (y\_ik - y\_i(t\_k, \theta))/\sigma_i(y\_i(t\_k, \theta), \theta*).$$

n1 and the constant-CV form n3 have no quantile residuals outside the envelope, and their Q–Q $R^2$ of 0.979 and 0.977 reflects a mild S-shape (Figure S10). The variants carrying the propagated control term have 38 (n4) and 31 (n7) of 103 residuals outside the envelope, and the Laplace variant 24. The variance-additive form n2 has the best-behaved residuals of all seven (Q–Q $R^2$ 0.989, Shapiro–Wilk $p = 0.710$) while increasing the AIC by 34.7. Since the residual behavior was overall good, we took the AIC value as the decisive criterion.

#### S1.8 Transcriptional delay

First-order mass action gives a graded lag but no dead time between promoter occupancy and mature transcript. We tested the effect of introducing a transcriptional delay, first on the topology of model M360 under the bounds of Section S1.6, then across the thirteen candidate topologies retained in the original submission.

##### Baseline without delay

Let $P^*_{\mathrm{Cyp}}(t)$ denote the transcriptionally active promoter complex `nAhR_L_ARNT_pCyp1a1` and $P^*_{\mathrm{AhRR}}(t)$ its *AhRR* counterpart. Transcription of each gene proceeds at

$$\beta_{\mathrm{Cyp}}(t) = b^{\mathrm{bas}}_{\mathrm{Cyp}}\, c_{\mathrm{Cyp}}\, \rho_{\mathrm{Cyp}}\, P^*_{\mathrm{Cyp}}(t) \;+\; b^{\mathrm{bas}}_{\mathrm{Cyp}}, \tag{15}$$

$$\beta_{\mathrm{AhRR}}(t) = b_{\mathrm{AhRR}}\, P^*_{\mathrm{AhRR}}(t) \;+\; b^{\mathrm{bas}}_{\mathrm{AhRR}}, \tag{16}$$

where $c_{\mathrm{Cyp}}$ is the ligand-specific fold change (unity for the 3MC reference condition). The mature transcript then obeys

$$\frac{\mathrm{d}[\mathrm{mRNA_g}]}{\mathrm{d}t} = \beta_g(t) \;-\; \tau_{\mathrm{mRNA},g}\,[\mathrm{mRNA_g}], \qquad g \in \{\mathrm{Cyp},\ \mathrm{AhRR}\}. \tag{17}$$

##### Delay by the linear chain trick

Transcription is redirected into a chain of $n$ nuclear intermediates $R_{g,1}, \ldots, R_{g,n}$, each transiting to the next at first-order rate $n/\tau$:

$$\frac{\mathrm{d}R_{g,1}}{\mathrm{d}t} = \beta_g(t) - \frac{n}{\tau}\, R_{g,1}, \tag{18}$$

$$\frac{\mathrm{d}R_{g,i}}{\mathrm{d}t} = \frac{n}{\tau}\, R_{g,i-1} - \frac{n}{\tau}\, R_{g,i}, \qquad i = 2, \ldots, n, \tag{19}$$

$$\frac{\mathrm{d}[\mathrm{mRNA_g}]}{\mathrm{d}t} = \frac{n}{\tau}\, R_{g,n} - \tau_{\mathrm{mRNA},g}\,[\mathrm{mRNA_g}]. \tag{20}$$

Eliminating the intermediates, (18)–(20) are exactly

$$\frac{\mathrm{d}[\mathrm{mRNA_g}]}{\mathrm{d}t} = \int_0^\infty g_{n,\tau}(s)\,\beta_g(t-s)\,\mathrm{d}s \;-\; \tau_{\mathrm{mRNA},g}\,[\mathrm{mRNA_g}], \tag{21}$$

with the Erlang density

$$g_{n,\tau}(s) \;=\; \frac{(n/\tau)^n\, s^{\,n-1}\, e^{-ns/\tau}}{(n-1)!}, \qquad \mathbb{E}[s] = \tau, \quad \mathrm{SD}[s] = \frac{\tau}{\sqrt{n}}, \quad \mathrm{CV} = \frac{1}{\sqrt{n}}. \tag{22}$$

The mean delay is therefore $\tau$ for every $n$, while $n$ controls only the *shape.* The limit $n \to \infty$ at fixed $\tau$ gives $g_{n,\tau} \to \delta(s-\tau)$, a pure dead time; $n = 1$ is the memoryless exponential, which is what unmodified first-order mass action produces [14].

Each intermediate is initialised at its own pre-stimulation steady state,

$$R_{g,i}(0) \;=\; \frac{b_g^{\mathrm{bas}}\,\tau}{n}, \qquad i = 1, \ldots, n, \tag{23}$$

obtained by setting (18) to zero at $\beta_g = b_g^{\mathrm{bas}}$ and propagating along the chain. Substituting (23) into (20) gives

$$[\mathrm{mRNA_g}]_{\mathrm{ss}} = \frac{1}{\tau_{\mathrm{mRNA},g}} \cdot \frac{n}{\tau} \cdot \frac{b_g^{\mathrm{bas}}\tau}{n} = \frac{b_g^{\mathrm{bas}}}{\tau_{\mathrm{mRNA},g}}, \tag{24}$$

which is *identical* to the steady state of (17). The delay therefore introduces no baseline offset, and the unstimulated state is unchanged for every $(n, \tau)$.

We fitted three fixed variants of $(n, \tau)$ sharing the same $n = 3$ and $\tau \in \{600, 1800, 3600\}$. Each was compared against the delay-free model under identical bounds in terms of the AIC (Table S1).

Table S1: **The single-topology transcriptional-delay grid.** The three delay variants of the grid, all fitted on the topology of model M360 under the bounds of Section S1.6. ΔAIC is measured against the delay-free model fitted under identical bounds, a negative value is a lower AIC for the delay variant. The variants add no free parameter, so their ΔAIC reflects the NLLH directly.

| variant | ΔAIC vs. no delay | added parameters |
|---|---:|---:|
| $n = 3$, $\tau = 600\,\mathrm{s}$ | −12.7 | 0 |
| $n = 3$, $\tau = 1800\,\mathrm{s}$ | −2.8 | 0 |
| $n = 3$, $\tau = 3600\,\mathrm{s}$ | −2.5 | 0 |

Only the 10 minute delay shows a significant improvement regarding the AIC criterion for this model. The test was consequently re-run as an invariance array over thirteen candidate model topologies, each fitted with and without a 10 min, three-stage Erlang delay (Figure S12). Across the thirteen topologies the mean AIC gain from the delay is $-0.7 \pm 3.7$, i.e. nothing on average. Nor is the delay a neutral common shift: the per-topology difference $\delta_i = \mathrm{AIC}_i(\mathrm{delay}) - \mathrm{AIC}_i(\mathrm{no\ delay})$, for which a positive value means the delay is worse, runs from $-4.8$ to $+9.8$ (M117 is hurt by 9.8), the Spearman rank correlation between the two arms is 0.70. Delays of 30 and 60 min are worse than 10 min. The delay thus interacts with the ligand-specific parameters, the very quantity the model selection measures, while contributing nothing on average. A delay of the order supported by the literature for transcription and translation (of the order of 10–20 min in total) is not resolvable with a first measurement at 2 h, and a delay of the order of hours is not supported. We retain the first-order formulation and the model selection of Section S1.9 carries no delay.

#### S1.9 Model selection

Model selection was based on AICc as the selection criterion, with AIC and NLLH recorded alongside and used a model space of 544 subspaces (528 distinct models). Each subspace was calibrated in three phases: 20 Fides multistarts, an 8 h saCeSS global search and a per-model refinement of the best candidates (Section S1.5). All 544 subspaces completed all three phases and the median wall time was 8.4 h. The median number of starts converging to the best value of a subspace is 614. Under the cutoff rule, $\Delta$AICc $\leq 10$, 9 models are retained (Table S9). The best model is M343 with 41 estimated parameters, NLLH 212.75, AICc 562.16, an Akaike weight of 0.69 and 312 converged starts. It carries a single ligand-specific parameter, `BpA_c1_dna_ahrr = 9.08`, the fold-change on the forward rate of the binding of the activated receptor complex to the *Ahrr* promoter ($R_{13}$): BpA-activated receptor binds the *Ahrr* promoter about nine times faster than 3MC- or ICZ-activated receptor, and ICZ needs no ligand-specific parameter relative to 3MC. The second best model M384 ($\Delta$AICc 3.27, weight 0.13) estimates `BpA_c_AhRR = 7.85; BpA_c_eL = 1e+05`.

Summing Akaike weights over models (Table S10): models with any BpA-specific parameter carry a weight of 0.97 (models with only BpA-specific parameters: 0.93), models with any ICZ-specific parameter 0.064, models with any 3MC-specific parameter 0.006; no model without a ligand-specific parameter receives weight. Models whose ligand-specific reactions are all at the promoter or transcription level carry 0.73 of the weight, models with any upstream ligand-specific reaction 0.27, and models with upstream ligand-specific reactions only 0.027. Per reaction, the forward binding to the *Ahrr* promoter accumulates 0.70 (BpA 0.70, ICZ 0.014), the extracellular ligand degradation 0.21, the *Ahrr* transcription rate 0.14, binding to and release from the *Cyp1a1* promoter together 0.076, and the *Cyp1a1* transcription rate, the step on which an ER$\alpha$-mediated effect would act, 0.050 (BpA 0.047, ICZ 0.019).

##### Number of ligand-specific reactions

An over-parameterised model can be made to fit any dataset, hence we tested this premise ins our framework. The saturated model, in which all ligand-specific fold-changes are estimated simultaneously (71 estimated parameters), is nested above every candidate of the space, so its optimum bounds the likelihood gain that any further ligand-specific parameter can buy. We performed three optimizations, warm-started from the optima of M76, M384 and M343 embedded in the saturated parameter vector. The best is `s5_full_model_from_M384` at NLLH 205.57 with 1 converged starts. The saturated model improves 14.35 in AIC on the likelihood term over the best model, so a candidate with $k$ additional estimated fold-changes can beat the best model by at most $14.35 - 2k$ AIC: 12.35 for one, 10.35 for two and 8.35 for three. Freeing one further reaction for both ligands costs two parameters, and the corresponding bound, 10.35 AIC, is already below the cutoff. No extension of the model space by further ligand-specific parameters can therefore produce an improvement this calibration would be able to resolve.

This bound by the full model says what is possible and we tested it directly with a forward step. From each of the two leading models we added, one reaction at a time, the ligand-specific fold-changes of every reaction the parent does not already free, for both ligands (+2 parameters), and calibrated each candidate. Table S14 lists all 43 of them. Every one of the 29 three-reaction models is worse than its parent. The best of them is M384+c1_heterodim, still 13.41 AICc behind the best model. No candidate beats the best model and not any does improve on their own parent beyond the AIC cutoff. A third ligand-specific reaction buys nothing on these data, as the saturated bound predicted, and the restriction of the model space to at most two ligand-specific reactions is justified.

#### Convergence of the model selection

The cold multistart is not sufficient for the estimation problem at hand. In none of the models did a single one of the 20 independent Fides starts come within 0.01 of the reported optimum, and the best start of a model misses it by between about two and sixty points in log-likelihood (Figure S6). The global phase is what finds the optimum. For every model of the table the reported value comes from the saCeSS phase or, for one model, from the refinement that follows it, and the merged worker histories then accumulate large numbers of recorded points within 0.01 of that value (Figure S5). To confirm our model selection result. the complete three-phase protocol was repeated from independent seeds for each of the 14 models of the table. And no run of any model reached a lower AICc than M343. In fact, all models reached comparable best likelihood values during the restarts as in the model selection (Table S12).

#### Goodness of fit criterions

The model selection was based on the Akaike information criterion and its corrected version for small data sizes. Information criteria weigh fit against complexity, so they do not order models the way an unpenalised fit statistic does. Table S16 gives both for every selected model: the residual root mean square error and the coefficient of determination next to NLLH, AIC and AICc. M343 is first on AICc but its RMSE is 1.26 times, its $R^2$ 0.74 times, that of the best-fitting candidate M117, which estimates three more parameters. Ranking on RMSE or $R^2$ alone would therefore select a more complex model whose extra likelihood the parameter count does not justify, which is why the selection is made on a penalised criterion rather than on a fit statistic. Also, a root mean squared error is not the quantity the likelihood optimises, because in the heteroskedastic noise model the likelihood weights each residual by its own $\sigma$, whereas a root mean squared error weights every residual equally. It is also, in practice, a single-channel statistic. *Cyp1a1* spans about ten times the range of *Ahrr*, so a pooled error over both observables is essentially the *Cyp1a1* error and carries almost no information about the *Ahrr* response.

#### S1.10 The best model and profile likelihoods

Looking at the fit of the best model M343, we observe that twelve parameter estimate lie within 0.01 decades of a bound (Figure S7). The selected fold-change (BpA_c1_dna_ahrr = 9.08) is not among them and sits well inside its bounds. The parameters that are at a bound are the ones of the unobserved protein layer, as in the bound audit (Section S1.6). Tables S15 carry the same comparison for every model of the selected set. To further investigate the parameter identifiabilty we ran profile likelihoods for the 41 parameters of this model. The confidence region is the one the uncertainty analysis of the main text defines $\mathrm{CR}^{\alpha} = \{\theta \mid \Lambda(\theta) \leq \Delta_{\alpha}\}$ with the likelihood-ratio statistic $\Lambda(\theta) = 2\,(J(\theta) - J(\theta^{*}))$ and $\Delta_{\alpha}$ the $\alpha$-quantile of $\chi^2$ with one degree of freedom. It is applied here one coordinate at a time: the parameter is fixed on a grid, the remaining 40 are re-optimised, and the 95 % interval is where the re-optimised profile crosses $\Delta = \chi^2_1(0.95) = 3.84$, i.e. a rise of 1.92 in the negative log-likelihood (likelihood ratio 0.1465). Interval ends are linearly interpolated between the two profile points that bracket the threshold.

All 41 estimated parameters of M343 are profiled: the ten ligand-relevant ones, the core parameters of the shared topology and the four noise parameters. The optimisation bounds in force for each parameter are given in Table S5 and their provenance in Table S2. As this means many optimisation runs, profiling took approximately 19 hours on 8 processes.

The resulting intervals of the parameter profiles are summarized in Table S17. 13 of the 41 parameters are identifiable at the 95 % level, i.e. their profiles cross the threshold on both sides. The decisive one is the selected fold-change: `BpA_c1_dna_ahrr` has a closed interval [3.38, 36.1] around an estimate of 9.08, 1.03 decades wide, and that interval excludes 1. So, the fold-change is not only the winner of a model ranking, it is a parameter the data bound away from the no-effect value. Three of the ten ligand-relevant parameters are identifiable, and the other two are the association constant of the same promoter reaction, `k_diss_dna_ahrr` (the parameter $d_{13}$) at 0.158 with [$2.64 \times 10^{-4}$, 1.57], and the *Cyp1a1* induction strength $k_{18}$ (`rho_Cyp`) at $2.44 \times 10^{6}$ with [$2.59 \times 10^{4}$, $4.23 \times 10^{7}$]. The remaining ten closed intervals are spread over the core model and the observation model: the nuclear import rate $k_8$ and the production and degradation of cytosolic AhR, $k_5$ and $k_6$, the degradation of nuclear ARNT, $k_{11}$, and of *Cyp1a1* mRNA, $k_{20}$, the two association constants with which AhRR occupies the *Ahrr* and *Cyp1a1* promoters, $k_{27}$ and $k_{28}$, the scaling factor of the *Ahrr* mRNA observable, `s_mRNA_AhRR_1`, and the proportional noise term of each observable, $\sigma_{1,\mathrm{mRNA}_{\mathrm{Cyp1a1}}}$ and $\sigma_{1,\mathrm{mRNA}_{\mathrm{AhRR}}}$. Because every transferable piece of literature information was already used to set the bounds (Section S1.6), the plausibility check on an estimate is where it sits inside its audited interval rather than a separate post-hoc comparison with published values. The cases in which a direct comparison is possible are AhR turnover ($k_6$, $k_9$), the receptor–ligand affinity ($d_7$, anchored on an ICZ $K_D$ of 0.19 nM) and *Cyp1a1* mRNA turnover ($k_{20}$). Each estimate lies inside the interval its source implies.

26 of the 41 parameters are bounded on one side only, 13 with a lower limit and 13 with an upper

limit. In every one of these 26 cases the open side ends at the parameter's own box bound rather than at a threshold crossing. The last 2 parameters are practically non-identifiable. Among the ten ligand-relevant parameters the BpA ligand-degradation constant $k_4$, the extracellular ligand degradation rate $k_1$ and the induced *Ahrr* transcription rate $k_{15}$ are bounded from below only, while the AhR–ligand association and dissociation constants $k_7$ and $d_7$, the association constant of the activator with the *Ahrr* promoter $k_{13}$ and the basal *Ahrr* transcription rate $k_{16}$ are bounded from above only. These are the parameters that act only through species the experiment never measures – receptor and promoter binding constants, protein turnover – and they trade off against each other and against the scaling factors.

#### S1.11 Trajectory ensembles and clustering

The prediction uncertainty and the clustering of model trajectories rest on the 9 retained models of the model selection (Section S1.9).

The ensemble of a model consists of all recorded optimisation end points with NLLH $\leq$ NLLH$^* + \chi^2_1(0.95) = 3.84$ above the model's own optimum, duplicates removed. The list is then sorted by objective value and stride-subsampled to at most 500 vectors per model, a cap only M394 reaches (500 vectors). The cutoff 3.84 is the 95 % quantile of $\chi^2_1$, i.e. the pointwise likelihood-based confidence region, applied per model. Over the 9 models of the ensemble, 377872 recorded optimisation points lie inside their model's region adn they reduce to 781 distinct parameter vectors.

For the clustering we used the two observable trajectories for the three ligand conditions, six series per parameter vector, each affinely min–max normalised over all ensemble members and all time points. The distance between two vectors is the Euclidean distance per series averaged over the six series and we used agglomerative clustering with complete linkage on that precomputed distance. The number of clusters follows the rule largest $k$ for which no cluster holds less than 1 % of the trajectories, which gives $k = 2$ and first fails at $k = 3$.

The Akaike-weight based contribution metric for a cluster is computed by the weights of the contained models. A model $m$ carries the weight $\tilde{w}_m$ and we spread it uniformly over the $n_m$ vectors of the model's own ensemble, so that each vector carries $\tilde{w}_m/n_m$. The weight of cluster $c$ is the sum over its members,

$$W_c = \sum_{(m,i)\,\in\, c} \frac{\tilde{w}_m}{n_m} = \sum_m \tilde{w}_m \, \frac{n_{m,c}}{n_m},$$

where $n_{m,c}$ is the number of vectors of model $m$ in cluster $c$, so it admits $\sum_c W_c = 1$.

Figure S4 shows the envelopes of all 2 clusters against the measured time points. Cluster 1 carries 99.1 % of the Akaike weight and 95.8 % of the 781 vectors, with 748 members contributed by 8 of the 9 models (Table S11). The only other cluster, 33 vectors or 4.2 % of them, carries 0.9 % of the weight and is drawn from the ensembles of M117, M76.

### S2 Supplementary Figures

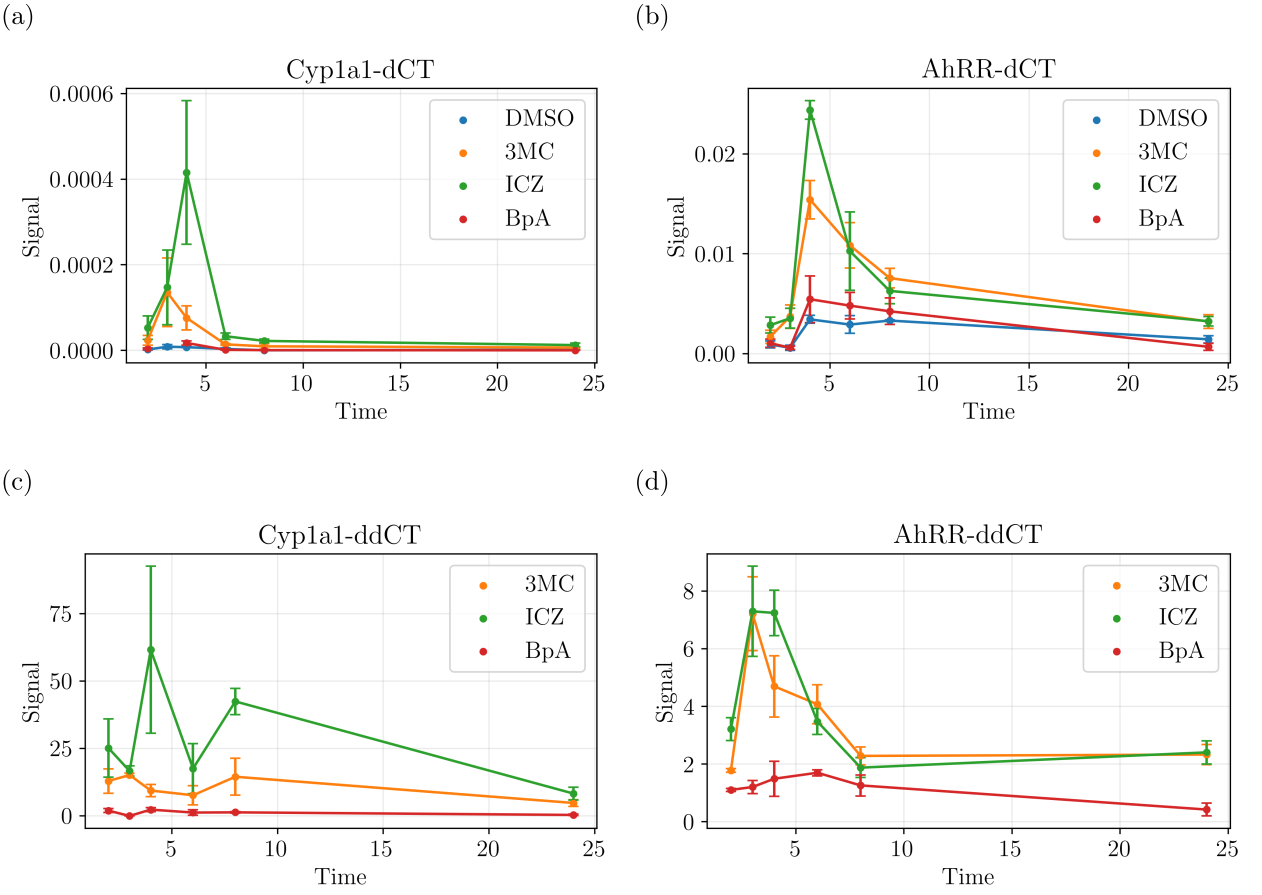

Figure S1: **BMDM kinetics data.** Relative qPCR measurement values for *Cyp1a1* and *Ahrr* measured in BMDMs after treatment with DMSO vehicle control, 3MC (1 $\mu$M), ICZ (10 $\mu$M), or BpA (1 $\mu$M). (a) and (b) $\Delta C_T$ values normalized to the housekeeping gene $\beta$-actin, (c) and (d) $\Delta\Delta C_T$ values formed replicate-wise, dividing the $2^{-\Delta C_T}$ value of each treated replicate by that of the paired DMSO replicate, not by the DMSO mean (Section S1.1).

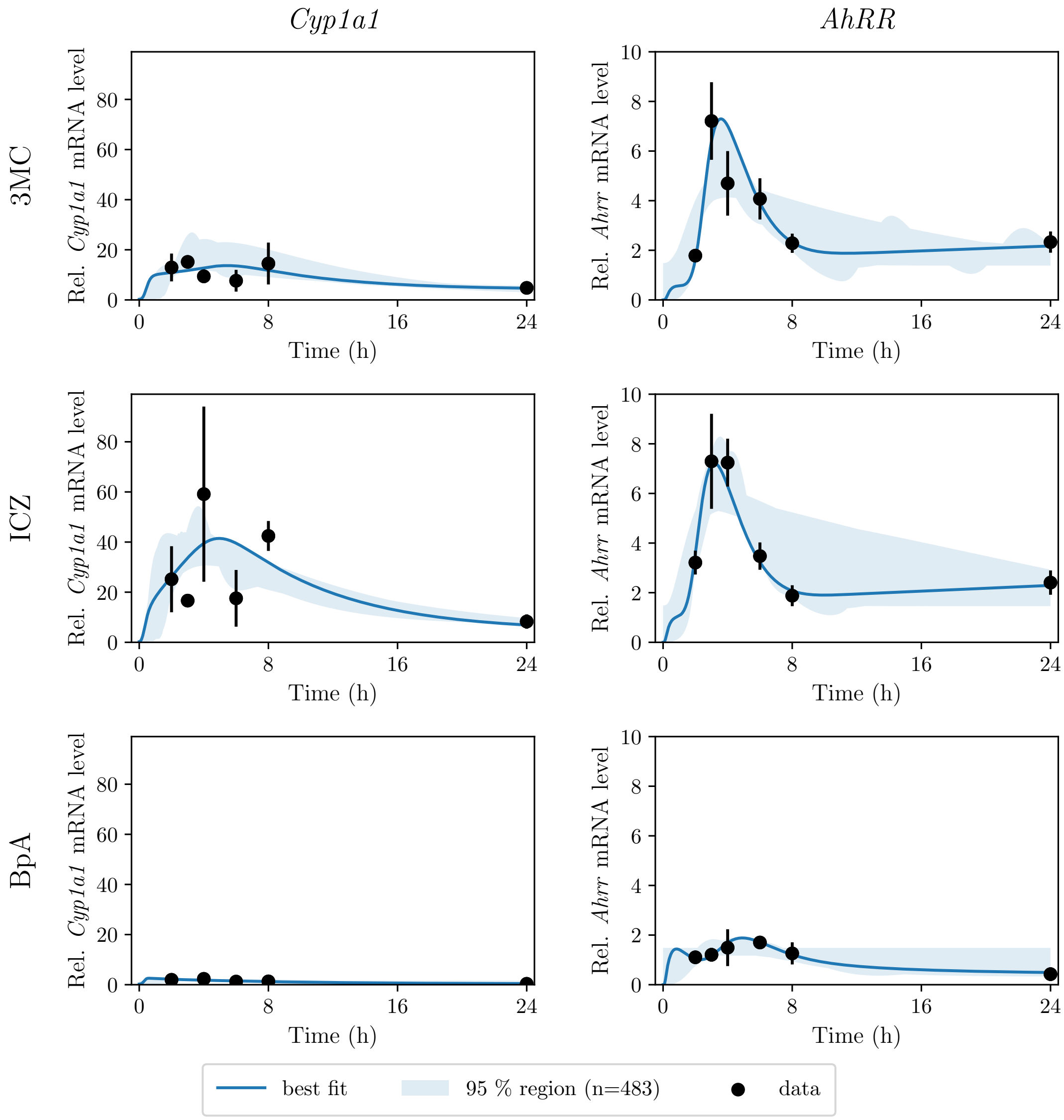

Figure S2: **Prediction uncertainty of the best model.** Simulations of M343 for every parameter vector of its likelihood-based ensemble (the rule is stated in Section S1.11), with the best fit and the *simultaneous* 95 % region, against the measurements. Rows are the ligands and columns the observables, the points are the mean over the biological replicates with error bars std/$\sqrt{n-1}$, at the six measured times 2, 3, 4, 6, 8 and 24 h.

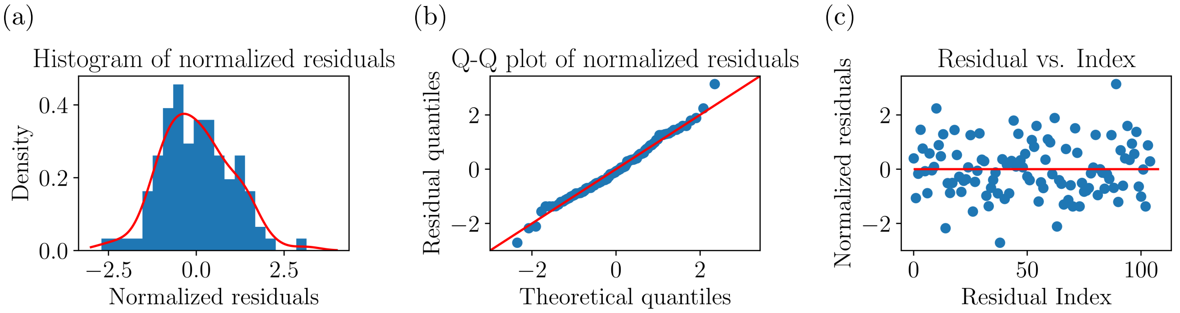

Figure S3: **Residual diagnostics for the heteroskedastic noise model.** Histogram of the residuals of M343 with gaussian kernel density estimate (left), quantile-quantile plot of the residuals (middle), and residual vs. index plot (right). Residuals are computed per individual replicate observation against the model prediction at that time point, $r_{ik} = \big(y_{ik} - y_i(t_k, \theta^*)\big)/\sigma_i\big(y_i(t_k, \theta^*), \theta^*\big)$, and pooled across the three ligand conditions ($n = 105$).

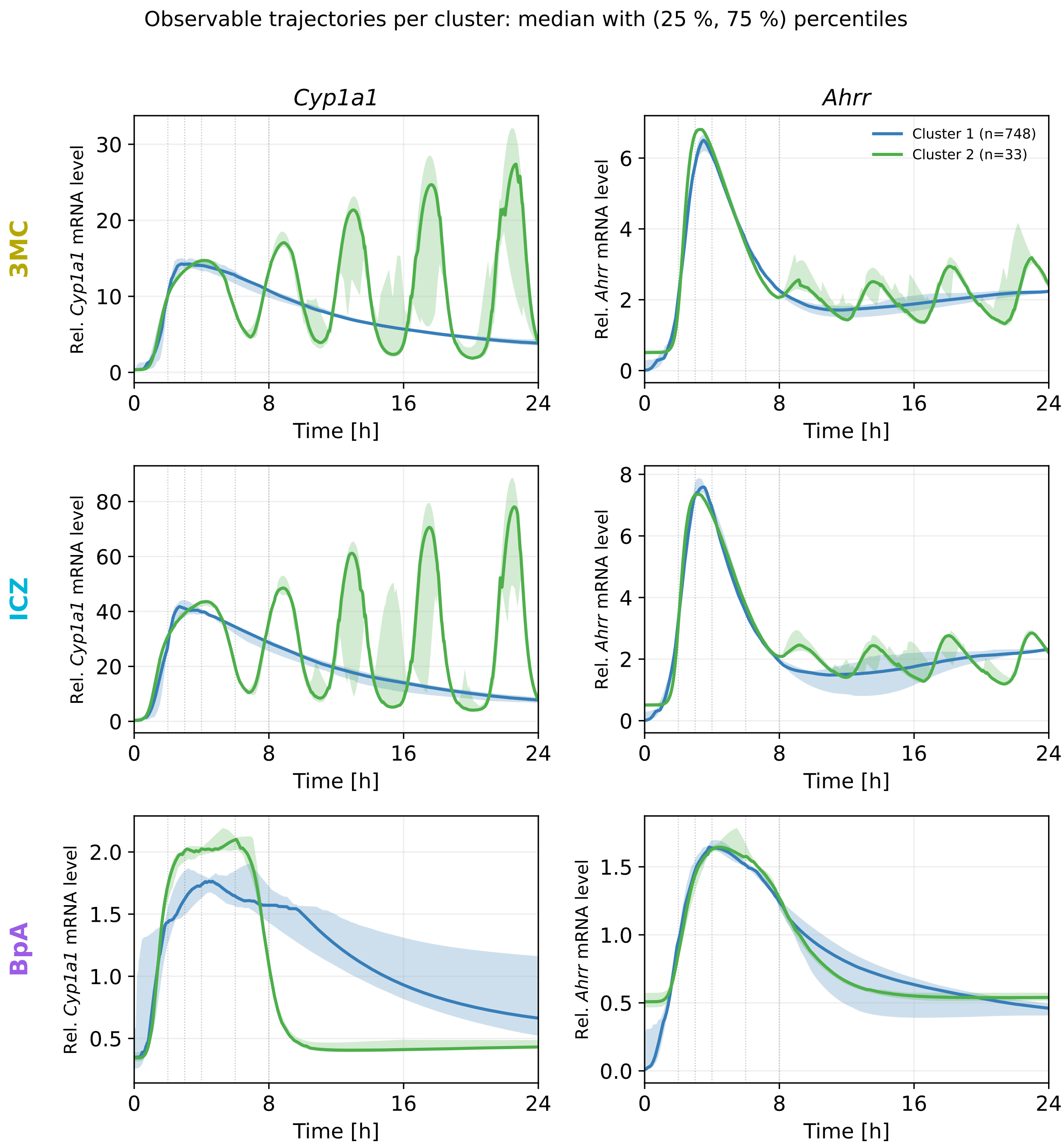

Figure S4: **Cluster envelopes against the measurement schedule.** Observable trajectories with pointwise median and (25 %, 75 %) percentiles for all 2 clusters of the ensemble, with the six measured time points marked.

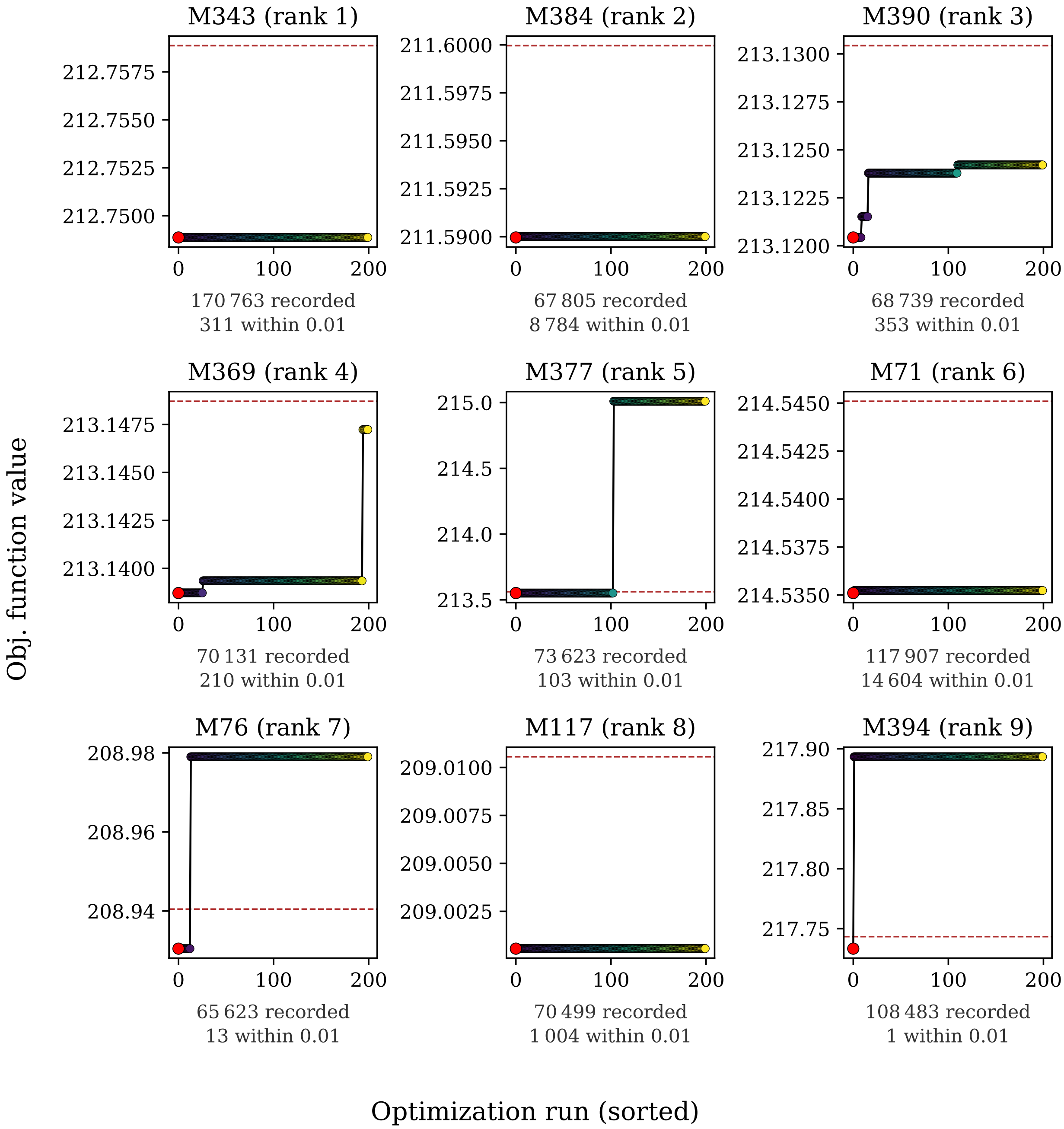

Figure S5: **Waterfall plots of the sweep, all recorded points.** Best objective values recorded for each of the 9 retained models. The recorded negative log-likelihoods in ascending order, the reported optimum in red and the convergence line at the optimum plus 0.01 dashed. Each panel states how many points were recorded and how many lie within 0.01 of the optimum.

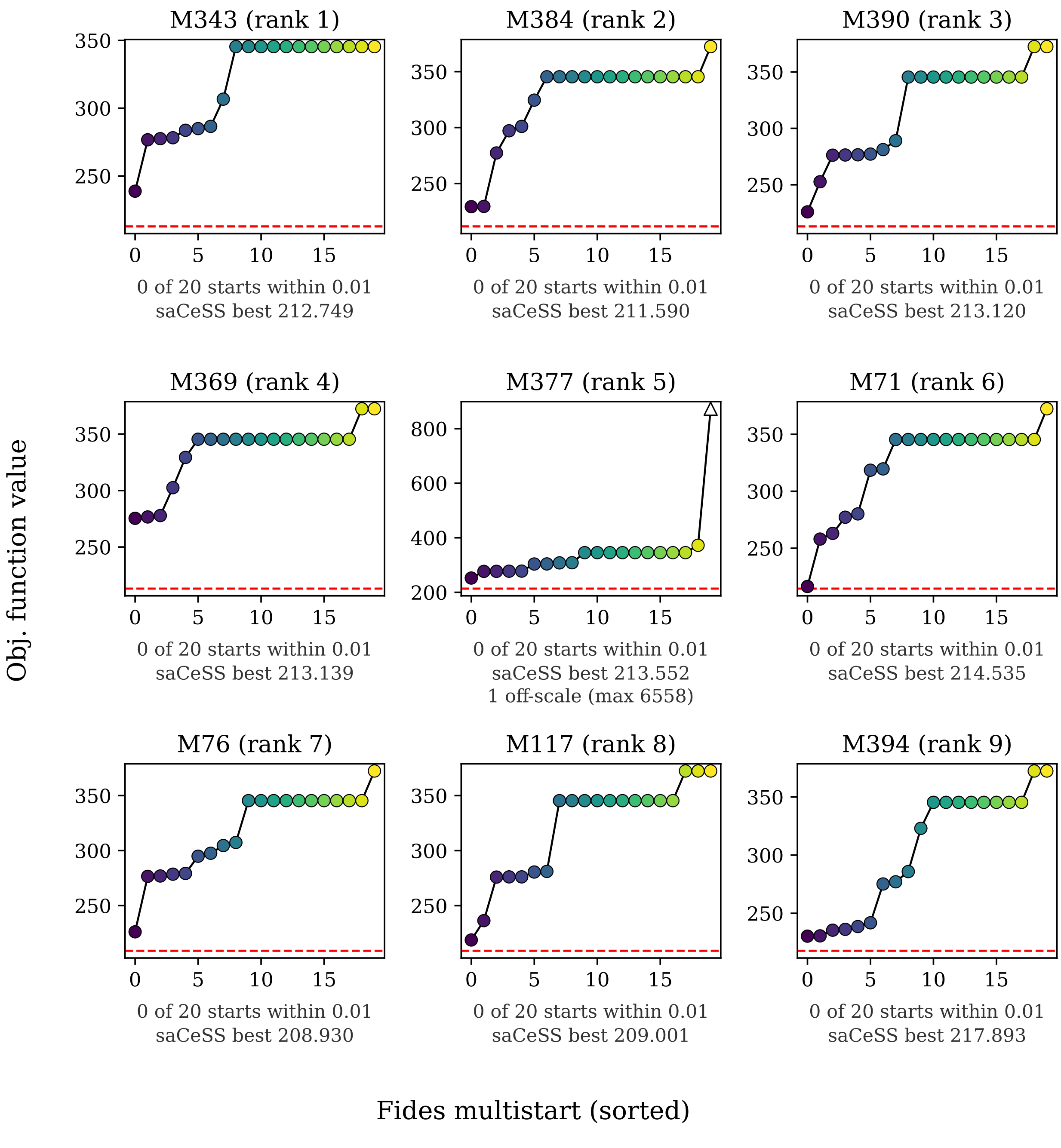

Figure S6: **The twenty independent cold multistarts of each model.** The first phase of the protocol alone, for the 9 retained models. The end points of the 20 independent Fides starts in ascending order, with the reported optimum as a red dashed reference line, and in each panel label how many starts came within 0.01 of it and what the best value of the global phase was.

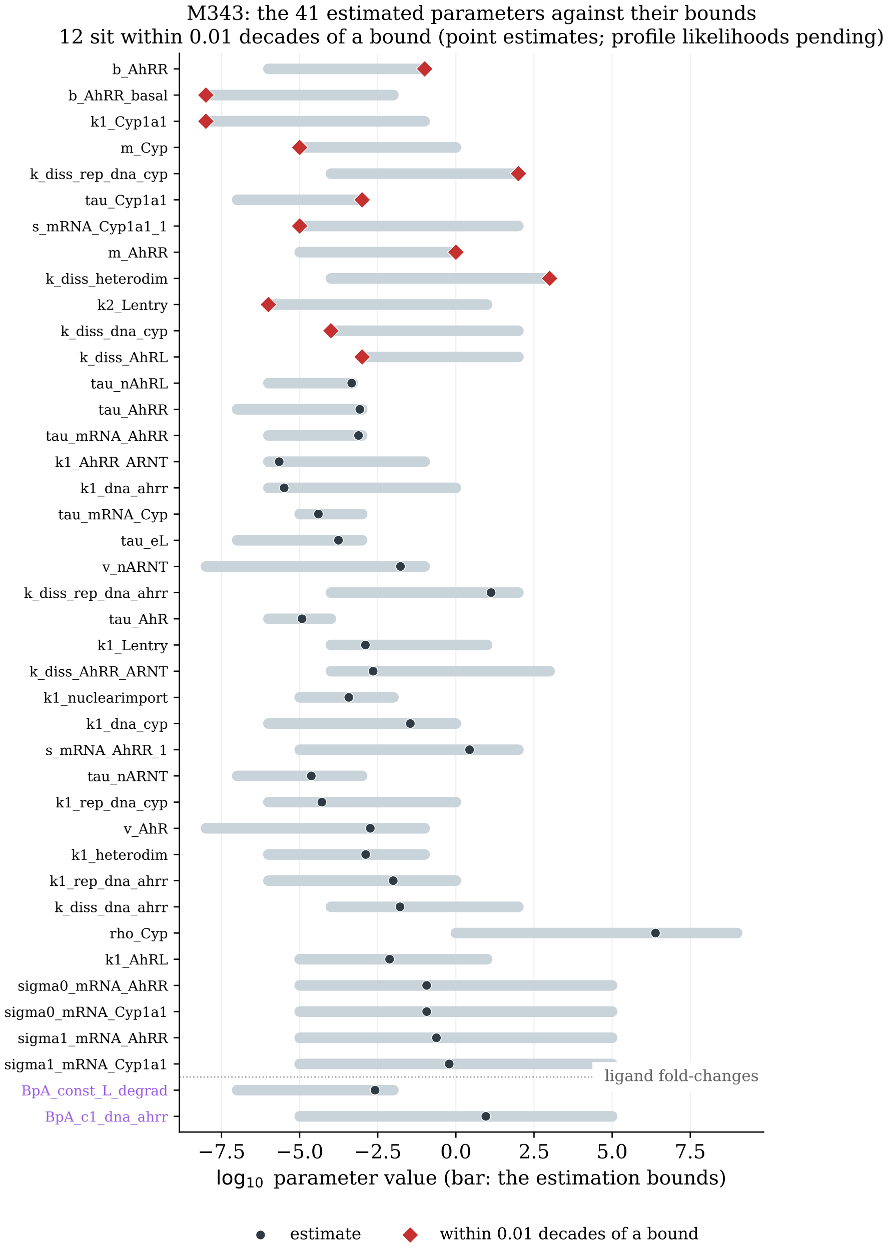

Figure S7: **Estimates of the best model against their bounds.** One row per estimated parameter of M343: the bound interval as a bar on a $\log_{10}$ axis, the estimate as a point, and the estimates within 0.01 decades of a bound marked. Core parameters first, then the ligand-specific fold-changes; rows are ordered by distance to the nearest bound.

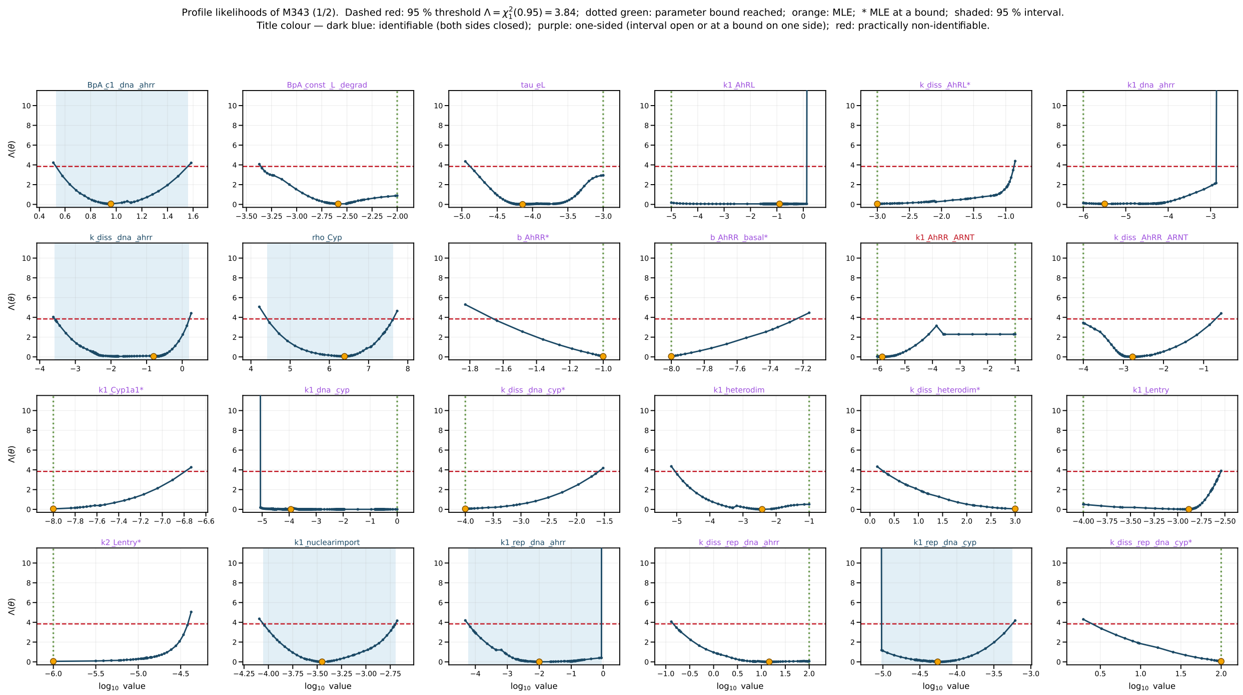

Figure S8: **Profile likelihoods of the best model (1 of 2).** One panel per estimated parameter of M343; all 41 are profiled, the first 24 here and the remaining 17 in Figure S9, in the order of Table S17 (ligand-relevant parameters first, then the core parameters of the shared topology, then the noise parameters). Each panel shows the likelihood-ratio statistic $\Lambda(\theta)$ of the re-optimised profile against the fixed parameter on a $\log_{10}$ axis. Dashed red line: the 95 % threshold $\Lambda = \chi^2_1(0.95) = 3.84$. Dotted green line: a box bound of the parameter reached by the path. Orange point: the maximum-likelihood estimate, with an asterisk in the panel title when it sits on a bound. Shading: the 95 % interval where it is closed on both sides.

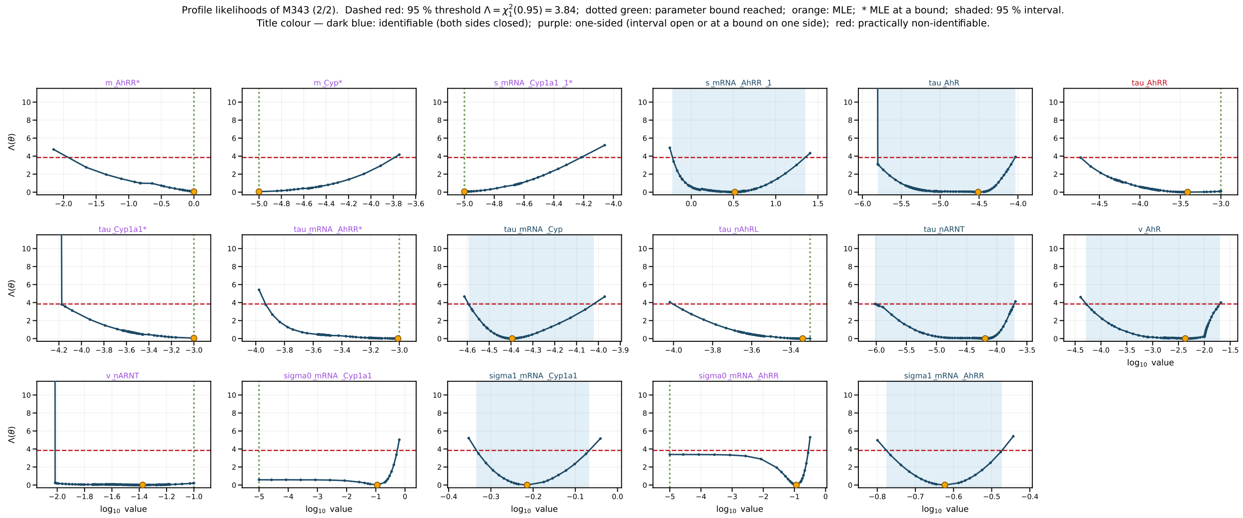

Figure S9: **Profile likelihoods of the best model (2 of 2).** Continued from Figure S8: the remaining 17 estimated parameters of M343, in the order of Table S17, ending with the four noise parameters. Lines, points, shading and title colours are as in Figure S8.

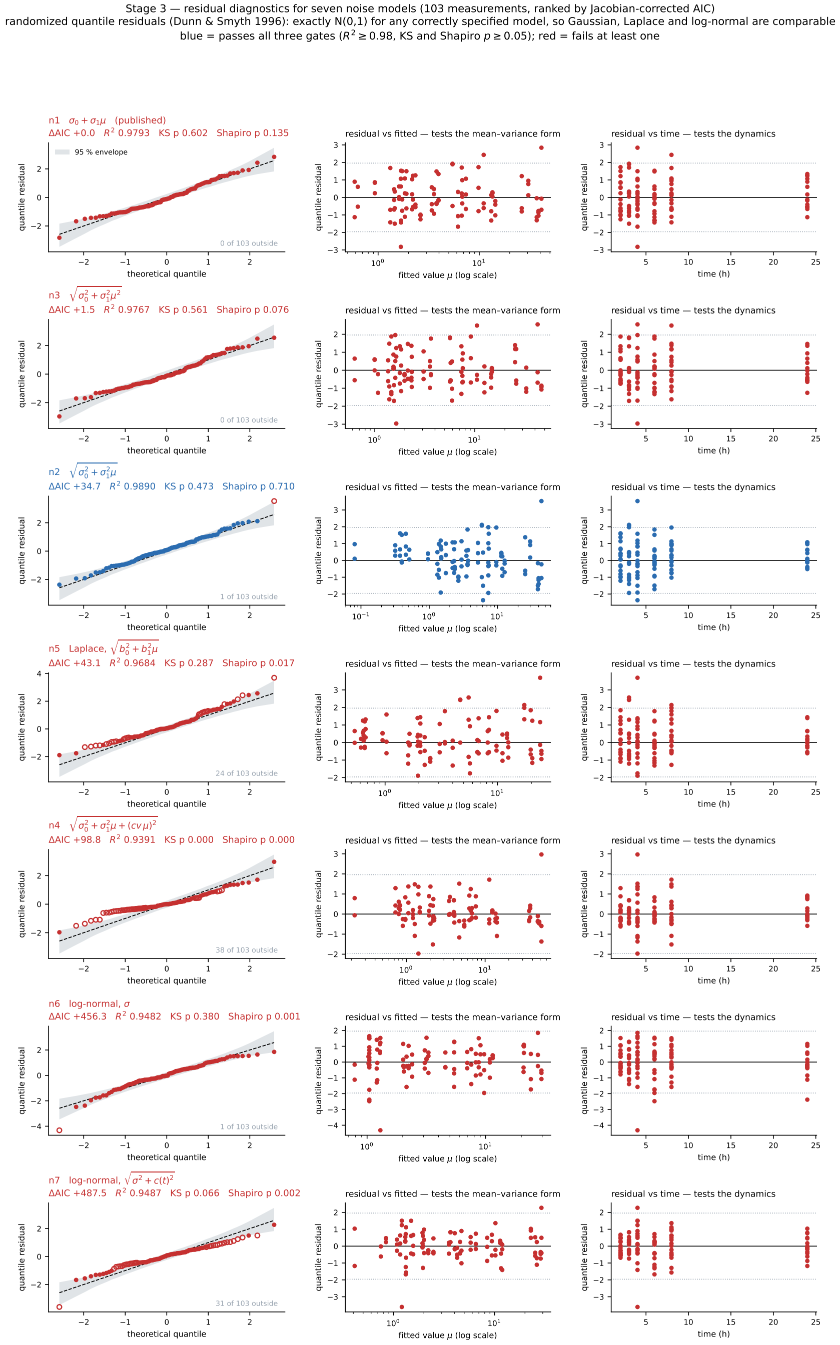

Figure S10: **Residual diagnostics for seven noise models.** Randomized quantile residuals $r_i = \Phi^{-1}(F_i(y_i))$, which are $N(0, 1)$ for any correctly specified model, for the seven error models of Table S7, fitted on the topology of model M360 to the 103 strictly positive measurements and ranked by Jacobian-corrected AIC. Per variant: Q–Q plot with a 95 % pointwise envelope and the number of residuals outside it (left), residual against fitted value, which tests the mean–variance form (middle), and residual against time, which tests the dynamics (right).

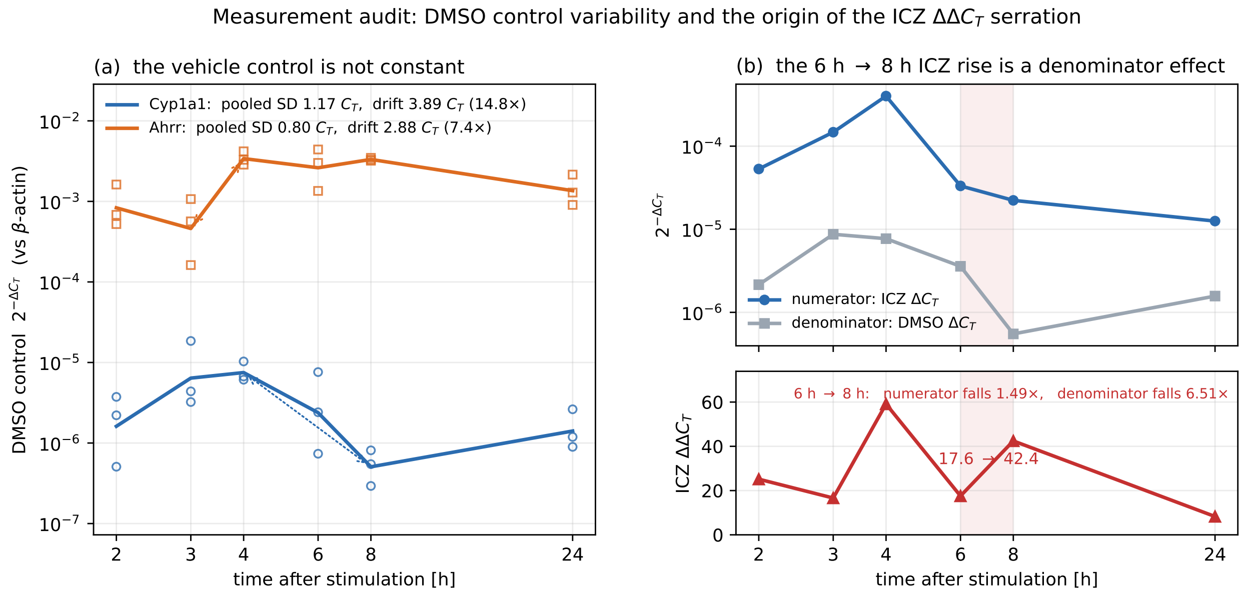

Figure S11: **Measurement audit: control variability and the origin of the ICZ serration.** (a) The DMSO vehicle control $2^{-\Delta C_T}$ (relative to $\beta$-actin) across the time course, all replicates and their geometric mean: pooled within-time-point SD 1.17 $C_T$ and drift 3.89 $C_T$ (14.8-fold) for *Cyp1a1*, 0.80 $C_T$ and 2.88 $C_T$ (7.4-fold) for *Ahrr*. (b) The ICZ *Cyp1a1* numerator and the DMSO denominator on one axis (top) and their ratio, the $\Delta\Delta C_T$ value (bottom).

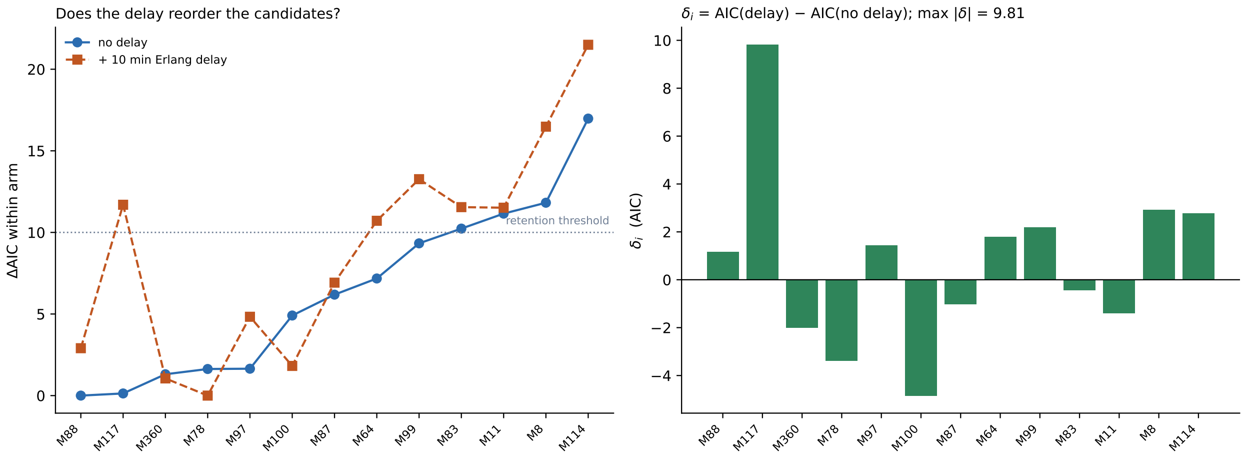

Figure S12: **Transcriptional delay across thirteen candidate topologies.** Left: ΔAIC within each arm (no delay vs. 10 min, three-stage Erlang delay) for the thirteen topologies under the bounds of Section S1.6, with the retention threshold. Right: per-topology difference $\delta_i = \mathrm{AIC}_i(\text{delay}) - \mathrm{AIC}_i(\text{no delay})$, the largest $|\delta_i|$ being 9.8 (M117).

### S3 Supplementary Tables

Table S2: **Literature audit of the parameter bounds.** The 37 numeric bound pairs of the model, each with the source the bound rests on. Evidence tiers: **A**, same molecule, mouse, BMDM or macrophage; **B**, same molecule, mouse, different cell type; **C**, same biochemical process in mammalian cells, or generic biophysics; **D**, qualitative mechanistic evidence only. Sources are given as Digital Object Identifiers (DOIs) where available.

| Par. | PEtab id | lb | ub | Tier | Basis | Source |
|---|---|---|---|---|---|---|
| $k_1$ | `tau_eL` | $10^{-7}$ | $10^{-3}$ | D | No measured medium half-life for 3MC/ICZ/BpA in RPMI + 10 % FCS; the window it replaces was uncited and 0.1 decades wide. | 10.1074/jbc.270.38.22548; 10.1124/jpet.109.162222 |
| $k_2$ | `k1_Lentry` | $10^{-4}$ | $10^{1}$ | D | Effective whole-cell transport, not a molecular rate; $k = PA/V$ gives $\approx 4.8 \cdot 10^3 P$ for a 1 pL cell. | 10.1074/jbc.270.38.22548 |
| $k_3$ | `k2_Lentry` | $10^{-6}$ | $10^{1}$ | D | Efflux of a hydrophobic ligand can be far slower than entry; $k_2/k_3$ is the cell:medium partition coefficient. | – |
| $k_4$ | `BpA_const_L_degrad` | $10^{-7}$ | $10^{-2}$ | D | BpA is cleared by UGT-mediated glucuronidation, not by CYP1A1; mechanism qualitative, no BMDM rate. | 10.1124/dmd.30.11.1180; 10.1248/bpb.24.1064; 10.1042/bj3400405 |
| $k_5$ | `v_AhR` | $10^{-8}$ | $10^{-1}$ | D | No absolute BMDM AhR synthesis rate; bounded by the order of magnitude of the measured abundance. | 10.1038/s41467-022-35095-7; 10.1038/s41590-023-01483-4 |
| $k_6$ | `tau_AhR` | $10^{-6}$ | $10^{-4}$ | B | Unliganded AhR $t_{1/2} = 28\,\mathrm{h}$ in mouse Hepa-1, i.e. $6.9 \cdot 10^{-6}\,\mathrm{s}^{-1}$; the ceiling is already 15-fold faster. | 10.1074/jbc.275.12.8432 |
| $k_7$ | `k1_AhRL` | $10^{-5}$ | $10^{1}$ | C | Small-ligand association typically $10^6$–$10^9\,\mathrm{M}^{-1}\mathrm{s}^{-1}$, Smoluchowski ceiling $\approx 10^{10}$; the buried PAS-B pocket implies the low end. | 10.1006/meth.2001.1171; 10.1016/S0026-895X(25)09356-3; 10.1021/cr800373w |
| $d_7$ | `k_diss_AhRL` | $10^{-3}$ | $10^{2}$ | B | The most direct evidence in the model: ICZ $K_D = 0.19\,\mathrm{nM}$, 3MC in the nM range in C57BL/6 liver cytosol. 3MC is the baseline. | 10.1073/pnas.88.21.9543; 10.1016/0006-2952(84)90303-4 |
| $k_8$ | `k1_nuclearimport` | $10^{-5}$ | $10^{-2}$ | C | AhR accumulates in the nucleus over minutes to tens of minutes; the ceiling corresponds to a 70 s time constant. | 10.1074/jbc.270.38.22548 |
| $k_9$ | `tau_nAhRL`† | $10^{-6}$ | $5 \cdot 10^{-4}$ | B | Ligand binding shortens the AhR half-life from 28 h to $\approx 3\,\mathrm{h}$, i.e. $6.4 \cdot 10^{-5}\,\mathrm{s}^{-1}$. | 10.1074/jbc.275.12.8432; 10.1074/jbc.274.40.28708 |
| $k_{10}$ | `v_nARNT` | $10^{-8}$ | $10^{-1}$ | D | No quantitative BMDM ARNT synthesis rate. | 10.1074/jbc.273.38.24867 |
| $k_{11}$ | `tau_nARNT` | $10^{-7}$ | $10^{-3}$ | D | No reliable value found; deliberately not tightened from generic protein turnover. | 10.1074/jbc.273.38.24867 |
| $k_{12}$ | `k1_heterodim` | $10^{-6}$ | $10^{-1}$ | C | Generic protein–protein association $10^4$–$10^7\,\mathrm{M}^{-1}\mathrm{s}^{-1}$, electrostatically accelerated to $\approx 10^8$. | 10.1002/prot.21679; 10.1002/prot.20489; 10.1021/cr800373w |
| $d_{12}$ | `k_diss_heterodim` | $10^{-4}$ | $10^{3}$ | D | The complex is established structurally; no robust native mouse $K_D$. | 10.1016/j.jmgm.2016.05.012; 10.1038/s41467-025-56574-7 |
| $k_{13}$ | `k1_dna_ahrr` | $10^{-6}$ | 1 | C | Specific protein–DNA association $10^7$–$10^9\,\mathrm{M}^{-1}\mathrm{s}^{-1}$; the upper end requires facilitated 1D diffusion along DNA. | 10.1101/gad.13.1.20; 10.1074/jbc.M011497200; 10.1021/bi00527a028 |
| $d_{13}$ | `k_diss_dna_ahrr` | $10^{-4}$ | $10^{2}$ | D | No quantitative BMDM affinity; an effective single site lumping several AhREs. | 10.1074/jbc.M011497200 |

*Continued on next page*

*Table continued from previous page*

| Par. | PEtab id | lb | ub | Tier | Basis | Source |
|---|---|---|---|---|---|---|
| $k_{14}$ | `k1_dna_cyp` | $10^{-6}$ | 1 | C | As $k_{13}$; the murine *Cyp1a1* promoter carries seven conserved AhREs, lumped here into one effective site. | 10.1016/0006-2952(94)90463-4; 10.3727/000000005783991981; 10.1039/C6MB00795C; 10.3390/ijms15046475 |
| $d_{14}$ | `k_diss_dna_cyp` | $10^{-4}$ | $10^{2}$ | C | Measured AhR:ARNT–DRE affinities span nM to hundreds of nM depending on construct and sequence. | 10.1016/0006-2952(94)90463-4; 10.3727/000000005783991981 |
| $k_{15}$ | `b_AhRR` | $10^{-6}$ | $10^{-1}$ | C | Concentration units cancel, so this is the per-copy initiation rate; a typical gene makes $\approx 2$ transcripts/h/copy, the Pol II re-initiation ceiling is $\approx 0.2\,\mathrm{s}^{-1}$. | 10.1074/jbc.M011497200; 10.1038/msb.2010.13; 10.1038/nbt.1861 |
| $k_{16}$ | `b_AhRR_basal` | $10^{-8}$ | $10^{-2}$ | D | A zeroth-order flux, not a rate: 0.5–5 transcripts/h/gene give $2.3 \cdot 10^{-6}$–$2.3 \cdot 10^{-5}\,\mathrm{nM\,s}^{-1}$ at $V_\mathrm{n} = 10^{-13}\,\mathrm{L}$. | 10.1074/jbc.M011497200 |
| $k_{17}$ | `tau_mRNA_AhRR` | $10^{-6}$ | $10^{-3}$ | C | No *Ahrr*-specific value; mouse dendritic cells under LPS give RNA half-lives of 10–70 min. | 10.1038/nbt.1861; 10.1038/nature10098 |
| $k_{18}$ | `rho_Cyp`* | 1 | $10^{9}$ | C | Not a rate: with $k_{19}$ fixed at 1 this is a ratio in $\mathrm{nM}^{-1}$. The ceiling has to be read at the calibrated promoter occupancy, not at saturation. | 10.1074/jbc.270.38.22548 |
| $k_{19}$ | `b_Cyp_basal` | – | – | fixed | Fixed at 1 to remove a scaling invariance. | – |
| $k_{20}$ | `tau_mRNA_Cyp` | $10^{-5}$ | $10^{-3}$ | B | *Cyp1a1* mRNA $t_{1/2} = 3$–$4\,\mathrm{h}$ in primary C57BL/6 hepatocytes ($4.8$–$6.4 \cdot 10^{-5}\,\mathrm{s}^{-1}$); the human orthologue agrees. | 10.1093/carcin/12.11.2115; 10.1006/abbi.2000.2115; 10.1074/jbc.270.38.22548 |
| $k_{21}$ | `m_Cyp` | $10^{-5}$ | 1 | C | Corrected NIH3T3 median of 140 proteins/mRNA/h, i.e. $3.9 \cdot 10^{-2}\,\mathrm{s}^{-1}$; apparent maximum 1000/h. | 10.1038/nature11848; 10.1038/nature10098; 10.1016/j.cell.2011.10.002 |
| $k_{22}$ | `tau_Cyp1a1` | $10^{-7}$ | $10^{-3}$ | D | No reliable value found; further complicated by substrate-accelerated (suicide) inactivation, so one first-order constant is an approximation. | – |
| $k_{23}$ | `k1_Cyp1a1` | $10^{-8}$ | $10^{-1}$ | D | An effective $k_\mathrm{cat}/K_M$, not elementary binding; the ceiling of $10^{8}\,\mathrm{M}^{-1}\mathrm{s}^{-1}$ respects catalytic-efficiency limits. | 10.1074/jbc.270.38.22548; 10.1002/ijc.2910200414 |
| $k_{24}$ | `m_AhRR` | $10^{-5}$ | 1 | C | No AhRR-specific rate; the generic corrected mouse translation distribution only. | 10.1038/nature11848 |
| $k_{25}$ | `tau_AhRR` | $10^{-7}$ | $10^{-3}$ | D | No measured AhRR protein half-life, consistent with the absence of a working antibody. | 10.1074/jbc.M117.812974 |
| $k_{26}$ | `k1_AhRR_ARNT` | $10^{-6}$ | $10^{-1}$ | C | The complex is structurally established but no $k_\mathrm{on}$ is measured; generic protein–protein association applies. | 10.1074/jbc.M117.812974; 10.1002/prot.21679 |
| $d_{26}$ | `k_diss_AhRR_ARNT` | $10^{-4}$ | $10^{3}$ | D | Structural evidence only; no native $K_D$ located. | 10.1074/jbc.M117.812974 |
| $k_{27}$ | `k1_rep_dna_ahrr` | $10^{-6}$ | 1 | D | AhRR binds AhREs, but repression is not explained by DNA competition and ARNT sequestration alone. | 10.1124/mol.107.040204 |
| $d_{27}$ | `k_diss_rep_dna_ahrr` | $10^{-4}$ | $10^{2}$ | D | No quantitative match found. | 10.1124/mol.107.040204 |
| $k_{28}$ | `k1_rep_dna_cyp` | $10^{-6}$ | 1 | D | Mouse MEF data support AhRR:ARNT occupancy of the *Cyp1a1* XRE; further transrepression mechanisms are documented. | 10.1016/j.bbrc.2007.09.131; 10.1124/mol.107.040204 |
| $d_{28}$ | `k_diss_rep_dna_cyp` | $10^{-4}$ | $10^{2}$ | D | No quantitative native affinity found. | 10.1016/j.bbrc.2007.09.131 |

Table S3: **BMDM kinetics raw data.** Raw expression values ($2^{-\Delta C_T}$ relative to $\beta$-actin) for *Cyp1a1* and *Ahrr* measured in BMDMs at the indicated timepoints after treatment with DMSO vehicle control, 3MC (1 $\mu$M), ICZ (10 $\mu$M), or BpA (1 $\mu$M). Three biological replicates are shown per condition and timepoint. The $\Delta\Delta C_T$ values of the calibration dataset are formed replicate-wise (treated replicate $i$ over DMSO replicate $i$).

| | DMSO | | | 3MC | | |
|---|---|---|---|---|---|---|
| Time (h) | Rep 1 | Rep 2 | Rep 3 | Rep 1 | Rep 2 | Rep 3 |
| *Cyp1a1* | | | | | | |
| 2 | 3.74e-6 | 2.21e-6 | 5.08e-7 | 1.85e-5 | 4.56e-5 | 6.73e-6 |
| 3 | 1.85e-5 | 3.23e-6 | 4.38e-6 | 2.96e-4 | 5.03e-5 | 6.10e-5 |
| 4 | 1.03e-5 | 6.11e-6 | 6.70e-6 | 1.29e-4 | 6.50e-5 | 3.39e-5 |
| 6 | 7.36e-7 | 2.41e-6 | 7.60e-6 | 1.05e-5 | 1.58e-5 | 1.63e-5 |
| 8 | 8.11e-7 | 2.93e-7 | 5.47e-7 | 2.28e-5 | 2.54e-6 | 3.71e-6 |
| 24 | 8.96e-7 | 1.19e-6 | 2.62e-6 | 6.55e-6 | 4.26e-6 | 9.2e-6 |
| *Ahrr* | | | | | | |
| 2 | 5.23e-4 | 1.62e-3 | 6.81e-4 | 9.56e-4 | 3.023e-3 | 1.13e-3 |
| 3 | 1.069e-3 | 5.65e-4 | 1.62e-4 | 4.979e-3 | 4.809e-3 | 1.371e-3 |
| 4 | 2.86e-3 | 4.187e-3 | 3.29e-3 | 1.92e-2 | 1.31e-2 | 1.39e-2 |
| 6 | 1.35e-3 | 3.01e-3 | 4.41e-3 | 7.33e-3 | 1.01e-2 | 1.51e-2 |
| 8 | 3.2e-3 | 3.3e-3 | 3.48e-3 | 9.05e-3 | 5.73e-3 | 7.94e-3 |
| 24 | 2.15e-3 | 1.29e-3 | 9.05e-4 | 4.58e-3 | 2.39e-3 | 2.72e-3 |

| | ICZ | | | BpA | | |
|---|---|---|---|---|---|---|
| Time (h) | Rep 1 | Rep 2 | Rep 3 | Rep 1 | Rep 2 | Rep 3 |
| *Cyp1a1* | | | | | | |
| 2 | 4.82e-5 | 1.03e-4 | 8.12e-6 | 3.34e-6 | 7.47e-6 | 8.13e-7 |
| 3 | 3.22e-4 | 6.32e-5 | 5.66e-5 | | | |
| 4 | 2.34e-4 | 7.05e-4 | 2.63e-4 | 2.22e-5 | 2.27e-5 | 7.68e-6 |
| 6 | 2.64e-5 | 2.52e-5 | 4.81e-5 | 2.44e-6 | 6.48e-7 | 1.17e-6 |
| 8 | 2.67e-5 | 1.33e-5 | 2.68e-5 | 1.06e-6 | 2.36e-7 | 1.02e-6 |
| 24 | 1.11e-5 | 5.1e-6 | 2.14e-5 | 8.24e-7 | 1.98e-7 | 0 |
| *Ahrr* | | | | | | |
| 2 | 1.58e-3 | 4.3e-3 | 2.7e-3 | 5.2e-4 | 1.87e-3 | 7.93e-4 |
| 3 | 4.98e-3 | 4.04e-3 | 1.63e-3 | 8.74e-4 | 6.76e-4 | 2.6e-4 |
| 4 | 2.5e-2 | 2.56e-2 | 2.261e-2 | 8.21e-4 | 8.32e-3 | 7.24e-3 |
| 6 | 5.08e-3 | 7.79e-3 | 1.79e-2 | 2.58e-3 | 4.73e-3 | 7.15e-3 |
| 8 | 3.97e-3 | 6.48e-3 | 8.43e-3 | 2.46e-3 | 3.44e-3 | 6.85e-3 |
| 24 | 4.22e-3 | 2.65e-3 | 2.9e-3 | 1.15e-3 | 9.56e-4 | 0 |

Table S4: **Biochemical reactions included in the AhR-pathway model.** Reactions and parameters are associated with their number in the graphical representation in Figure 1 of the main manuscript. The mapping of these 35 kinetic steps to the 24 reaction elements of the SBML file, to the PEtab parameter identifiers and to the ligand-specific fold-change parameters is given in Table S6.

| No. | Reaction | Description | Rate constants |
|---|---|---|---|
| $R_1$ | $L_e \rightarrow \varnothing$ | $L_e$ degradation | $k_1$ |
| $R_2$ | $L_e \rightarrow L_c$ | Ligand entry | $k_2$ |
| $R_3$ | $L_c \rightarrow L_e$ | Ligand exit | $k_3$ |
| $R_4$ | $L_c \rightarrow \varnothing$ | constant $L_c$ degradation | $k_4$ |
| $R_5$ | $\varnothing \rightarrow AhR_c$ | $AhR_c$ production | $k_5$ |
| $R_6$ | $AhR_c \rightarrow \varnothing$ | $AhR_c$ degradation | $k_6$ |
| $R_7$ | $AhR_c + L_c \rightarrow AhR_c{:}L_c$ | Ligand binding | $k_7$ |
| $R_{-7}$ | $AhR_c + L_c \leftarrow AhR_c{:}L_c$ | Ligand unbinding | $k_{-7}$ |
| $R_8$ | $AhR_c{:}L_c \rightarrow AhR_n{:}L_n$ | Nuclear import | $k_8$ |
| $R_9$ | $AhR_n{:}L_n \rightarrow \varnothing$ | $AhR_n$ degradation | $k_9$ |
| $R_{10}$ | $\varnothing \rightarrow ARNT_n$ | $ARNT_n$ production | $k_{10}$ |
| $R_{11}$ | $ARNT_n \rightarrow \varnothing$ | $ARNT_n$ degradation | $k_{11}$ |
| $R_{12}$ | $AhR_n{:}L_n + ARNT_n \rightarrow AhR_n{:}L_n{:}ARNT_n$ | $AhR_n$ heterodimerization | $k_{12}$ |
| $R_{-12}$ | $AhR_n{:}L_n + ARNT_n \leftarrow AhR_n{:}L_n{:}ARNT_n$ | $AhR_n$ dissociation | $k_{-12}$ |
| $R_{13}$ | $p_{Ahrr} + AhR_n{:}L_n{:}ARNT_n \rightarrow AhR_n{:}L_n{:}ARNT_n{:}p_{Ahrr}$ | $p_{Ahrr}$ activation | $k_{13}$ |
| $R_{-13}$ | $p_{Ahrr} + AhR_n{:}L_n{:}ARNT_n \leftarrow AhR_n{:}L_n{:}ARNT_n{:}p_{Ahrr}$ | $p_{Ahrr}$ inactivation | $k_{-13}$ |
| $R_{14}$ | $p_{Cyp1a1} + AhR_n{:}L_n{:}ARNT_n \rightarrow AhR_n{:}L_n{:}ARNT_n{:}p_{Cyp1a1}$ | $p_{Cyp1a1}$ activation | $k_{14}$ |
| $R_{-14}$ | $p_{Cyp1a1} + AhR_n{:}L_n{:}ARNT_n \leftarrow AhR_n{:}L_n{:}ARNT_n{:}p_{Cyp1a1}$ | $p_{Cyp1a1}$ inactivation | $k_{-14}$ |
| $R_{15}$ | $AhR_n{:}L_n{:}ARNT_n{:}p_{Ahrr} \rightarrow AhR_n{:}L_n{:}ARNT_n{:}p_{Ahrr} + mRNA_{AhRR}$ | $mRNA_{AhRR}$ transcription | $k_{15}$ |
| $R_{16}$ | $\varnothing \rightarrow mRNA_{AhRR}$ | $mRNA_{AhRR}$ basal transcription | $k_{16}$ |
| $R_{17}$ | $mRNA_{AhRR} \rightarrow \varnothing$ | $mRNA_{AhRR}$ degradation | $k_{17}$ |
| $R_{18}$ | $AhR_n{:}L_n{:}ARNT_n{:}p_{Cyp1a1} \rightarrow AhR_n{:}L_n{:}ARNT_n{:}p_{Cyp1a1} + mRNA_{Cyp1a1}$ | $mRNA_{Cyp1a1}$ transcription | $k_{18}$ |
| $R_{19}$ | $\varnothing \rightarrow mRNA_{Cyp1a1}$ | $mRNA_{Cyp1a1}$ basal transcription | $k_{19}$ |
| $R_{20}$ | $mRNA_{Cyp1a1} \rightarrow \varnothing$ | $mRNA_{Cyp1a1}$ degradation | $k_{20}$ |
| $R_{21}$ | $mRNA_{Cyp1a1} \rightarrow mRNA_{Cyp1a1} + Cyp1a1_c$ | $Cyp1a1_c$ translation | $k_{21}$ |
| $R_{22}$ | $Cyp1a1_c \rightarrow \varnothing$ | $Cyp1a1_c$ degradation | $k_{22}$ |
| $R_{23}$ | $L_c + Cyp1a1_c \rightarrow Cyp1a1_c$ | $L_c$ degradation | $k_{23}$ |
| $R_{24}$ | $mRNA_{AhRR} \rightarrow mRNA_{AhRR} + AhRR_n$ | $AhRR_n$ translation | $k_{24}$ |
| $R_{25}$ | $AhRR_n \rightarrow \varnothing$ | $AhRR_n$ degradation | $k_{25}$ |
| $R_{26}$ | $AhRR_n + ARNT_n \rightarrow AhRR_n{:}ARNT_n$ | $AhRR_n$ heterodimerization | $k_{26}$ |
| $R_{-26}$ | $AhRR_n + ARNT_n \leftarrow AhRR_n{:}ARNT_n$ | $AhRR_n$ dissociation | $k_{-26}$ |
| $R_{27}$ | $p_{Ahrr} + AhRR_n{:}ARNT_n \rightarrow AhRR_n{:}ARNT_n{:}p_{Ahrr}$ | $p_{Ahrr}$ repression | $k_{27}$ |
| $R_{-27}$ | $p_{Ahrr} + AhRR_n{:}ARNT_n \leftarrow AhRR_n{:}ARNT_n{:}p_{Ahrr}$ | $p_{Ahrr}$ derepression | $k_{-27}$ |
| $R_{28}$ | $p_{Cyp1a1} + AhRR_n{:}ARNT_n \rightarrow AhRR_n{:}ARNT_n{:}p_{Cyp1a1}$ | $p_{Cyp1a1}$ repression | $k_{28}$ |
| $R_{-28}$ | $p_{Cyp1a1} + AhRR_n{:}ARNT_n \leftarrow AhRR_n{:}ARNT_n{:}p_{Cyp1a1}$ | $p_{Cyp1a1}$ derepression | $k_{-28}$ |

Table S5: **Specification of model parameters.** The table includes the parameter names, the PEtab identifiers, lower bounds (lb) and upper bounds (ub) of the calibration, units ("-"=dimensionless) and fixed value if not estimated. Estimation is on $\log_{10}$ scale with a uniform prior between the bounds. The $d_i$ are association constants ($K_{D,i} = 1/d_i$). $k_4$ is estimated for BpA only and fixed to zero for 3MC and ICZ. The ligand-specific fold-change parameters $c$ multiply the respective rate constant in one ligand condition.

| **Parameter** | **PEtab id** | **lb** | **ub** | **Unit** | **Value** | |
|---|---|---|---|---|---|---|
| $k_1$ | `tau_eL` | 1.0e-07 | 1.0e-03 | $s^{-1}$ | Estimated | |
| $k_2$ | `k1_Lentry` | 1.0e-04 | 1.0e+01 | $s^{-1}$ | Estimated | |
| $k_3$ | `k2_Lentry` | 1.0e-06 | 1.0e+01 | $s^{-1}$ | Estimated | |
| $k_4$ | `BpA_const_L_degrad` | 1.0e-07 | 1.0e-02 | $s^{-1}$ | Estimated (BpA only) | |
| $k_5$ | `v_AhR` | 1.0e-08 | 1.0e-01 | $nM \cdot s^{-1}$ | Estimated | |
| $k_6$ | `tau_AhR` | 1.0e-06 | 1.0e-04 | $s^{-1}$ | Estimated | |
| $k_7$ | `k1_AhRL` | 1.0e-05 | 1.0e+01 | $s^{-1} \cdot nM^{-1}$ | Estimated | |
| $d_7$ | `k_diss_AhRL` | 1.0e-03 | 1.0e+02 | $nM^{-1}$ | Estimated | |
| $k_8$ | `k1_nuclearimport` | 1.0e-05 | 1.0e-02 | $s^{-1}$ | Estimated | |
| $k_9$ | `tau_nAhRL` | 1.0e-06 | 5.0e-04 | $s^{-1}$ | Estimated | |
| $k_{10}$ | `v_nARNT` | 1.0e-08 | 1.0e-01 | $nM \cdot s^{-1}$ | Estimated | |
| $k_{11}$ | `tau_nARNT` | 1.0e-07 | 1.0e-03 | $s^{-1}$ | Estimated | |
| $k_{12}$ | `k1_heterodim` | 1.0e-06 | 1.0e-01 | $s^{-1} \cdot nM^{-1}$ | Estimated | |
| $d_{12}$ | `k_diss_heterodim` | 1.0e-04 | 1.0e+03 | $nM^{-1}$ | Estimated | |
| $k_{13}$ | `k1_dna_ahrr` | 1.0e-06 | 1.0e+00 | $s^{-1} \cdot nM^{-1}$ | Estimated | |
| $d_{13}$ | `k_diss_dna_ahrr` | 1.0e-04 | 1.0e+02 | $nM^{-1}$ | Estimated | |
| $k_{14}$ | `k1_dna_cyp` | 1.0e-06 | 1.0e+00 | $s^{-1} \cdot nM^{-1}$ | Estimated | |
| $d_{14}$ | `k_diss_dna_cyp` | 1.0e-04 | 1.0e+02 | $nM^{-1}$ | Estimated | |
| $k_{15}$ | `b_AhRR` | 1.0e-06 | 1.0e-01 | $s^{-1}$ | Estimated | |
| $k_{16}$ | `b_AhRR_basal` | 1.0e-08 | 1.0e-02 | $nM \cdot s^{-1}$ | Estimated | |
| $k_{17}$ | `tau_mRNA_AhRR` | 1.0e-06 | 1.0e-03 | $s^{-1}$ | Estimated | |
| $k_{18}$ | `rho_Cyp` | 1.0e+00 | 1.0e+09 | $nM^{-1}$ | Estimated | |
| $k_{19}$ | `b_Cyp_basal` | - | - | - | $nM \cdot s^{-1}$ | 1.0 |
| $k_{20}$ | `tau_mRNA_Cyp` | 1.0e-05 | 1.0e-03 | default | $s^{-1}$ | Estimated |
| $k_{21}$ | `m_Cyp` | 1.0e-05 | 1.0e+00 | $s^{-1}$ | Estimated | |
| $k_{22}$ | `tau_Cyp1a1` | 1.0e-07 | 1.0e-03 | $s^{-1}$ | Estimated | |
| $k_{23}$ | `k1_Cyp1a1` | 1.0e-08 | 1.0e-01 | $s^{-1} \cdot nM^{-1}$ | Estimated | |
| $k_{24}$ | `m_AhRR` | 1.0e-05 | 1.0e+00 | $s^{-1}$ | Estimated | |
| $k_{25}$ | `tau_AhRR` | 1.0e-07 | 1.0e-03 | $s^{-1}$ | Estimated | |
| $k_{26}$ | `k1_AhRR_ARNT` | 1.0e-06 | 1.0e-01 | $s^{-1} \cdot nM^{-1}$ | Estimated | |
| $d_{26}$ | `k_diss_AhRR_ARNT` | 1.0e-04 | 1.0e+03 | $nM^{-1}$ | Estimated | |
| $k_{27}$ | `k1_rep_dna_ahrr` | 1.0e-06 | 1.0e+00 | $s^{-1} \cdot nM^{-1}$ | Estimated | |
| $d_{27}$ | `k_diss_rep_dna_ahrr` | 1.0e-04 | 1.0e+02 | $nM^{-1}$ | Estimated | |
| $k_{28}$ | `k1_rep_dna_cyp` | 1.0e-06 | 1.0e+00 | $s^{-1} \cdot nM^{-1}$ | Estimated | |
| $d_{28}$ | `k_diss_rep_dna_cyp` | 1.0e-04 | 1.0e+02 | $nM^{-1}$ | Estimated | |
| $total_{p_{Ahrr}}$ | `tot_pAhRR` | - | - | 2 copies/cell | nM | 3.32e-02 |
| $total_{p_{Cyp1a1}}$ | `tot_pCyp1a1` | - | - | 2 copies/cell | nM | 3.32e-02 |
| $s_{mRNA_{Cyp1a1}}$ | `s_mRNA_Cyp1a1_1` | 1.0e-05 | 1.0e+02 | default | - | Estimated |
| $s_{mRNA_{AhRR}}$ | `s_mRNA_AhRR_1` | 1.0e-05 | 1.0e+02 | default | - | Estimated |
| $\sigma_{0,mRNA_{Cyp1a1}}$ | `sigma0_mRNA_Cyp1a1` | 1.0e-05 | 1.0e+05 | default | - | Estimated |
| $\sigma_{1,mRNA_{Cyp1a1}}$ | `sigma1_mRNA_Cyp1a1` | 1.0e-05 | 1.0e+05 | default | - | Estimated |
| $\sigma_{0,mRNA_{AhRR}}$ | `sigma0_mRNA_AhRR` | 1.0e-05 | 1.0e+05 | default | - | Estimated |

| | | | | | | |
|---|---|---|---|---|---|---|
| $\sigma_{1,\mathrm{mRNA}_{\mathrm{AhRR}}}$ | sigma1_mRNA_AhRR | 1.0e-05 | 1.0e+05 | default | - | Estimated |
| $c$ (fold-changes) | {BpA,ICZ,MC3}_c* | 1.0e-05 | 1.0e+05 | default | - | per model |

Table S6: **Mapping of manuscript reactions to SBML reactions and model parameters.** The table lists the reaction numbering used in the manuscript (Table S4), the corresponding SBML reaction (reversible pairs, the two cytosolic ligand degradation steps and the induced/basal transcription pairs of each gene each share one SBML reaction element, so 24 elements encode the 35 kinetic steps), the associated rate constant(s) in the PEtab parameter table, and the ligand-specific fold-change parameter(s) defined in the PEtab condition table, which carry the ligand as prefix (BpA_, ICZ_, MC3_) and are fixed at one unless estimated by a candidate model.

| **Manuscript** | **SBML reaction** | **Rate constant(s)** | **Fold-change (conditions.tsv)** |
|---|---|---|---|
| $R_1$ | _1_0___eL_degradation | tau_eL | c_eL |
| $R_2$ / $R_3$ | _1_1___Ligand_entry | k1_Lentry / k2_Lentry | c1_Lentry / c2_Lentry |
| $R_4$ | _2_4___cL_degradation (term 1) | BpA_const_L_degrad | BpA-only, fixed elsewhere |
| $R_5$ / $R_6$ | _2_1_AhR_production / _2_3_AhR_degradation | v_AhR / tau_AhR | – |
| $R_7$ / $R_{-7}$ | _2_2_Ligand_binding | k1_AhRL / k_diss_AhRL | c1_AhRL / c2_AhRL |
| $R_8$ | _3_1_Nuclear_Import | k1_nuclearimport | c1_nuclearimport |
| $R_9$ | _3_2_nAhR_degradation | tau_nAhRL | c_nAhRL |
| $R_{10}$ / $R_{11}$ | _3_3_nARNT_production / _6_2_nARNT_degradation | v_nARNT / tau_nARNT | – |
| $R_{12}$ / $R_{-12}$ | _3_4_Heterodimerization | k1_heterodim / k_diss_heterodim | c1_heterodim / c2_heterodim |
| $R_{13}$ / $R_{-13}$ | _5_0___DNA_binding_AhRR | k1_dna_ahrr / k_diss_dna_ahrr | c1_dna_ahrr / c2_dna_ahrr |
| $R_{14}$ / $R_{-14}$ | _4_0___DNA_binding_Cyp1a1 | k1_dna_cyp / k_diss_dna_cyp | c1_dna_cyp / c2_dna_cyp |
| $R_{15}$ / $R_{16}$ | _5_1___mRNA_AhRR_expression | b_AhRR / b_AhRR_basal | c_AhRR |
| $R_{17}$ | _5_6___mRNA_AhRR_degradation | tau_mRNA_AhRR | – |
| $R_{18}$ / $R_{19}$ | _4_1___mRNA_Cyp1a1_expression | rho_Cyp / b_Cyp_basal (=1) | c_Cyp |
| $R_{20}$ | _4_5___mRNA_Cyp1a1_degradation | tau_mRNA_Cyp | – |
| $R_{21}$ / $R_{22}$ | _4_2___Cyp1a1_expression / _4_3___Cyp1a1_degradation | m_Cyp / tau_Cyp1a1 | – |
| $R_{23}$ | _2_4___cL_degradation (term 2) | k1_Cyp1a1 | c1_Cyp1a1 |
| $R_{24}$ / $R_{25}$ | _5_2___AhRR_expression / _5_2_AhRR_degradation | m_AhRR / tau_AhRR | – |
| $R_{26}$ / $R_{-26}$ | _5_2___AhRR_inhibition | k1_AhRR_ARNT / k_diss_AhRR_ARNT | – |
| $R_{27}$ / $R_{-27}$ | _5_4___Inhibition_AhRR_expression | k1_rep_dna_ahrr / k_diss_rep_dna_ahrr | – |
| $R_{28}$ / $R_{-28}$ | _4_3___Inhibition_Cyp1a1_expression | k1_rep_dna_cyp / k_diss_rep_dna_cyp | – |

Table S7: **Noise-model comparison on the topology of model M360.** Seven error models (defined in Section S1.7) fitted to the 103 strictly positive measurements with identical model, bounds and optimisation protocol. $k$ is the number of estimated parameters. Residual diagnostics use randomized quantile residuals.

| variant | $\sigma$ | ΔAIC | Q–Q $R^2$ | KS $p$ | Shapiro $p$ | outside the 95 % envelope |
|---|---|---|---|---|---|---|
| **n1 (original)** | $\sigma_0 + \sigma_1\mu$ | 0 | 0.979 | 0.602 | 0.136 | 0 / 103 |
| n3 | $\sqrt{}(\sigma_0{}^2 + \sigma_1{}^2\mu^2)$ | +1.5 | 0.977 | 0.561 | 0.076 | 0 / 103 |
| n2 | $\sqrt{}(\sigma_0{}^2 + \sigma_1{}^2\mu)$ | +34.7 | 0.989 | 0.473 | 0.710 | 1 / 103 |
| n5 | Laplace, $\sqrt{}(b_0{}^2 + b_1{}^2\mu)$ | +43.1 | 0.968 | 0.287 | 0.017 | 24 / 103 |
| n4 = n2 + control variance | | +98.8 | 0.939 | $1.4 \times 10^{-4}$ | $2.9 \times 10^{-4}$ | 38 / 103 |
| n6 | log-normal | +456.3 | 0.948 | 0.380 | $7.8 \times 10^{-4}$ | 1 / 103 |
| n7 = n6 + control variance | | +487.5 | 0.949 | 0.066 | $1.5 \times 10^{-3}$ | 31 / 103 |

Table S8: **Noise-model ranking across four model topologies.** ΔAIC within each arm (Jacobian-corrected, 103 strictly positive measurements) of the seven error models of Table S7 on the topologies M360 (42 free parameters), M117 and M64 (44 free parameters) and the saturated model (71 free parameters).

| variant | error model | M360 | M117 | M64 | saturated |
|---|---|---|---|---|---|
| n1 | $\sigma_0 + \sigma_1\mu$ (adopted) | 0.0 | 0.0 | 3.5 | 0.0 |
| n3 | $\sqrt{\sigma_0^2 + \sigma_1^2\mu^2}$ | 6.9 | 2.4 | 0.0 | 7.6 |
| n2 | $\sqrt{\sigma_0^2 + \sigma_1^2\mu}$ | 38.9 | 40.9 | 34.8 | 17.2 |
| n5 | Laplace | 63.5 | 49.3 | 87.8 | 49.0 |
| n4 | n2 + control variance | 73.0 | 59.6 | 72.3 | 46.9 |
| n6 | log-normal | 448 | 445 | 453 | 437 |
| n7 | log-normal + control variance | 482 | 480 | 483 | 463 |

Table S9: Models within the cutoff of 10 AICc of the best model.

| rank | model | $k$ | ligand-specific fold-changes | NLLH | AICc | ΔAICc | weight |
|---|---|---|---|---|---|---|---|
| 1 | M343 | 41 | BpA_c1_dna_ahrr | 212.75 | 562.16 | 0.00 | 0.686 |
| 2 | M384 | 42 | BpA_c_AhRR, BpA_c_eL | 211.59 | 565.44 | 3.27 | 0.134 |
| 3 | M390 | 42 | BpA_c_Cyp, BpA_c_eL | 213.12 | 568.50 | 6.33 | 0.029 |
| 4 | M369 | 42 | BpA_c1_dna_cyp, BpA_c_eL | 213.14 | 568.54 | 6.37 | 0.028 |
| 5 | M377 | 42 | BpA_c2_dna_cyp, BpA_c_eL | 213.55 | 569.36 | 7.20 | 0.019 |
| 6 | M71 | 42 | BpA_c1_dna_ahrr, ICZ_c1_dna_ahrr | 214.54 | 571.33 | 9.16 | 0.007 |
| 7 | M76 | 44 | BpA_c1_dna_ahrr, BpA_c_Cyp, ICZ_c1_dna_ahrr, ICZ_c_Cyp | 208.93 | 571.86 | 9.70 | 0.005 |
| 8 | M117 | 44 | BpA_c_Cyp, BpA_c_nAhRL, ICZ_c_Cyp, ICZ_c_nAhRL | 209.00 | 572.00 | 9.84 | 0.005 |
| 9 | M394 | 41 | BpA_c_nAhRL | 217.73 | 572.13 | 9.97 | 0.005 |

Table S10: Accumulated Akaike weight of each ligand-specific reaction over all calibrated models, and its split by the ligand whose fold-change was estimated.

| parameter | reaction | level | accumulated weight | BpA | ICZ | 3MC |
|---|---|---|---|---|---|---|
| c1_dna_ahrr | AhR:L:ARNT binding to the Ahrr promoter | promoter | 0.702 | 0.701 | 0.014 | 0.000 |
| c_eL | extracellular ligand degradation | upstream | 0.213 | 0.211 | 0.001 | 0.001 |
| c_AhRR | Ahrr transcription rate | transcription | 0.143 | 0.139 | 0.004 | 0.002 |
| c_Cyp | Cyp1a1 transcription rate | transcription | 0.050 | 0.047 | 0.019 | 0.001 |
| c1_dna_cyp | AhR:L:ARNT binding to the Cyp1a1 promoter | promoter | 0.038 | 0.031 | 0.008 | 0.001 |
| c2_dna_cyp | AhR:L:ARNT release from the Cyp1a1 promoter | promoter | 0.038 | 0.031 | 0.014 | 0.002 |
| c2_heterodim | heterodimerisation (reverse) | upstream | 0.017 | 0.014 | 0.009 | 0.000 |
| c_nAhRL | nuclear AhR:L degradation | upstream | 0.015 | 0.014 | 0.007 | 0.000 |
| c1_heterodim | heterodimerisation (forward) | upstream | 0.015 | 0.010 | 0.010 | 0.001 |
| c2_dna_ahrr | AhR:L:ARNT release from the Ahrr promoter | promoter | 0.013 | 0.011 | 0.004 | 0.000 |
| c1_nuclearimport | nuclear import | upstream | 0.004 | 0.002 | 0.002 | 0.000 |
| c2_AhRL | ligand binding (reverse) | upstream | 0.004 | 0.002 | 0.002 | 0.000 |
| c1_AhRL | ligand binding (forward) | upstream | 0.004 | 0.002 | 0.001 | 0.000 |
| c1_Cyp1a1 | Cyp1a1-mediated ligand degradation | upstream | 0.003 | 0.000 | 0.002 | 0.001 |
| c1_Lentry | ligand entry (forward) | upstream | 0.003 | 0.001 | 0.002 | 0.000 |
| c2_Lentry | ligand entry (reverse) | upstream | 0.002 | 0.001 | 0.001 | 0.001 |

Table S11: **Cluster membership of the retained models.** Akaike-weighted fraction of the whole ensemble that each model contributes to each cluster (the raw number of ensemble members is given in brackets). Rows sum to the cluster share, columns to the model's renormalised Akaike weight.

| Cluster | M343 | M384 | M390 | M369 | M377 | M71 | M76 | M117 | M394 | $\sum$ |
|---|---|---|---|---|---|---|---|---|---|---|
| 1 | 0.738 (2) | 0.144 (25) | 0.031 (10) | 0.031 (23) | 0.031 (18) | 0.008 (142) | – | 0.004 (28) | 0.005 (500) | 0.991 |
| 2 | – | – | – | – | – | – | 0.006 (9) | 0.003 (24) | – | 0.009 |
| $\sum$ | 0.738 | 0.144 | 0.031 | 0.031 | 0.031 | 0.008 | 0.006 | 0.007 | 0.005 | 1.000 |

Table S12: Independent repetitions of the complete calibration protocol (8 seeds per model) for the selected models. Spread: largest minus smallest NLLH over the model selection run and all restarts.

| model | rank | selection NLLH | best restart | spread (NLLH) | within floor |
|---|---|---|---|---|---|
| M343 | 1 | 212.75 | 218.73 | 11.10 | 1/9 |
| M384 | 2 | 211.59 | 211.83 | 8.73 | 2/9 |
| M390 | 3 | 213.12 | 213.27 | 8.73 | 3/9 |
| M369 | 4 | 213.14 | 213.11 | 9.80 | 3/9 |
| M377 | 5 | 213.55 | 213.13 | 9.29 | 3/9 |
| M71 | 6 | 214.54 | 214.54 | 7.30 | 8/9 |
| M76 | 7 | 208.93 | 213.95 | 5.74 | 3/9 |
| M117 | 8 | 209.00 | 208.70 | 7.98 | 6/9 |
| M394 | 9 | 217.73 | 218.79 | 2.45 | 9/9 |
| M53 | 10 | 209.10 | 209.04 | 6.02 | 6/9 |
| M101 | 11 | 214.98 | 212.56 | 3.14 | 9/9 |
| M228 | 12 | 217.82 | 219.27 | 2.64 | 9/9 |
| M63 | 13 | 209.27 | 208.80 | 5.27 | 9/9 |
| M360 | 43 | 217.10 | 211.60 | 6.45 | 4/9 |

Table S13: **Fits of the saturated model** (all 31 ligand-specific fold-changes estimated, 71 parameters). We did three warm starts embedding different optima of model selection models.

| fit | warm start | wall (h) | NLLH |
|---|---|---|---|
| s5_full_model_from_M384 | embedded M384 optimum | 205.57 | |
| s5_full_model_from_M76 | embedded M76 optimum | 205.66 | |
| s5_full_model_from_M343 | embedded M343 optimum | 206.52 | |

Table S14: **One forward step from the two leading models.** Each model adds the fold-changes of one further reaction, for both ligands where both are free (+2 parameters; +1 where only one ligand can deviate), to its parent (M343, M384). A positive ΔAICc vs. parent means the extension is worse. Sorted by distance to the best model of the selection process (M343, AICc 562.16).

| model | parent | $k$ | NLLH | AICc | ΔAICc vs. parent | ΔAICc vs. best | converged starts |
|---|---|---|---|---|---|---|---|
| M384+c1_heterodim | M384 | 44 | 210.79 | 575.58 | +10.14 | +13.41 | 366 |
| M343+c2_dna_cyp | M343 | 43 | 215.41 | 578.86 | +16.70 | +16.70 | 31 |
| M343+c1_dna_cyp | M343 | 43 | 215.47 | 578.98 | +16.82 | +16.82 | 2206 |
| M384+c1_dna_ahrr | M384 | 44 | 213.76 | 581.52 | +16.08 | +19.36 | 12 |
| M343+c1_heterodim | M343 | 43 | 216.80 | 581.63 | +19.46 | +19.46 | 274 |
| M384+c2_dna_cyp | M384 | 44 | 214.42 | 582.84 | +17.40 | +20.67 | 256 |
| M384+c1_dna_cyp | M384 | 44 | 214.45 | 582.90 | +17.46 | +20.73 | 9373 |
| M384+c1_Cyp1a1 | M384 | 43 | 217.59 | 583.22 | +17.78 | +21.06 | 1 |
| M343+c_Cyp | M343 | 43 | 217.61 | 583.25 | +21.09 | +21.09 | 16 |
| M384+c_Cyp | M384 | 44 | 214.82 | 583.64 | +18.20 | +21.48 | 122 |
| M343+c2_dna_ahrr | M343 | 43 | 217.96 | 583.95 | +21.79 | +21.79 | 434 |
| M384+c2_dna_ahrr | M384 | 44 | 215.14 | 584.29 | +18.85 | +22.12 | 1 |
| M343+c_nAhRL | M343 | 43 | 218.20 | 584.43 | +22.27 | +22.27 | 144 |
| M343+c2_heterodim | M343 | 43 | 218.45 | 584.93 | +22.76 | +22.76 | 116 |
| M343+c_eL | M343 | 43 | 219.13 | 586.28 | +24.12 | +24.12 | 299 |
| M384+c_nAhRL | M384 | 44 | 216.41 | 586.83 | +21.39 | +24.66 | 19 |
| M343+c1_Cyp1a1 | M343 | 42 | 223.24 | 588.73 | +26.57 | +26.57 | 493 |
| M343+c1_nuclearimport | M343 | 43 | 220.51 | 589.05 | +26.89 | +26.89 | 684 |
| M384+c2_AhRL | M384 | 44 | 217.64 | 589.28 | +23.84 | +27.11 | 304 |
| M343+c2_Lentry | M343 | 43 | 220.64 | 589.31 | +27.14 | +27.14 | 11 |
| M384+c1_AhRL | M384 | 44 | 217.69 | 589.38 | +23.94 | +27.22 | 254 |
| M384+c2_heterodim | M384 | 44 | 218.05 | 590.10 | +24.66 | +27.93 | 2831 |
| M343+c2_AhRL | M343 | 43 | 221.24 | 590.51 | +28.34 | +28.34 | 197 |
| M343+c1_AhRL | M343 | 43 | 221.38 | 590.80 | +28.64 | +28.64 | 218 |
| M343+c1_Lentry | M343 | 43 | 221.66 | 591.36 | +29.20 | +29.20 | 1 |
| M343+c_AhRR | M343 | 43 | 221.76 | 591.55 | +29.38 | +29.38 | 1 |
| M384+c1_nuclearimport | M384 | 44 | 219.44 | 592.88 | +27.44 | +30.71 | 342 |
| M384+c1_Lentry | M384 | 44 | 219.51 | 593.01 | +27.57 | +30.85 | 274 |
| M384+c2_Lentry | M384 | 44 | 220.00 | 594.01 | +28.57 | +31.84 | 138 |

Table S15: Estimates at a box bound in the retained set ($\Delta$AICc $\leq$ 10). An estimate counts as sitting at a bound when it is within 0.01 decades of the nearer bound, the rule used for the `n_at_bound` column of the ranking. $k$ is the number of estimated parameters. The classes partition the estimated parameters into the ligand-independent kinetic parameters, the four noise parameters, and the ligand fold-changes.

| rank | model | $k$ | at bound | kinetic | noise | fold-change | fold-change at a bound |
|---|---|---|---|---|---|---|---|
| 1 | M343 | 41 | 12 | 11 | 0 | 0 | – |
| 2 | M384 | 42 | 9 | 7 | 0 | 1 | `BpA_c_eL` |
| 3 | M390 | 42 | 10 | 9 | 0 | 0 | – |
| 4 | M369 | 42 | 9 | 8 | 0 | 0 | – |
| 5 | M377 | 42 | 11 | 9 | 0 | 1 | `BpA_c_eL` |
| 6 | M71 | 42 | 14 | 13 | 0 | 0 | – |
| 7 | M76 | 44 | 10 | 10 | 0 | 0 | – |
| 8 | M117 | 44 | 10 | 10 | 0 | 0 | – |
| 9 | M394 | 41 | 10 | 9 | 0 | 0 | – |
| total | | | 95 | 86 | 0 | 2 | |

Table S16: **Goodness of fit and information criteria for the selected models.** The three qPCR replicates of a condition × observable × time cell disagree with one another, so no deterministic model can do better than predict the cell mean: over the 35 cells that floor is $\mathrm{RMSE}_{\mathrm{min}} = 8.089$ and $R^2_{\mathrm{max}} = 0.6910$. The ratio columns state each model against that floor, a ratio of 1 would be a perfect fit. $R^2 = 1 - \mathrm{SSR}/\mathrm{SST}$ is descriptive only.

| model | $k$ | NLLH | RMSE | $\mathrm{RMSE}/\mathrm{RMSE}_{\mathrm{min}}$ | $R^2$ | $R^2/R^2_{\mathrm{max}}$ | AIC | AICc | ΔAICc | rank AIC/AICc |
|---|---|---|---|---|---|---|---|---|---|---|
| M343 | 41 | 212.75 | 10.206 | 1.262 | 0.5081 | 0.735 | 507.50 | 562.16 | 0.00 | 4/1 |
| M384 | 42 | 211.59 | 10.289 | 1.272 | 0.5001 | 0.724 | 507.18 | 565.44 | 3.27 | 3/2 |
| M390 | 42 | 213.12 | 10.234 | 1.265 | 0.5055 | 0.731 | 510.24 | 568.50 | 6.33 | 5/3 |
| M369 | 42 | 213.14 | 10.212 | 1.262 | 0.5076 | 0.734 | 510.28 | 568.54 | 6.37 | 6/4 |
| M377 | 42 | 213.55 | 10.172 | 1.258 | 0.5114 | 0.740 | 511.10 | 569.36 | 7.20 | 7/5 |
| M71 | 42 | 214.54 | 10.410 | 1.287 | 0.4883 | 0.707 | 513.07 | 571.33 | 9.16 | 8/6 |
| M76 | 44 | 208.93 | 10.234 | 1.265 | 0.5054 | 0.731 | 505.86 | 571.86 | 9.70 | 1/7 |
| M117 | 44 | 209.00 | 9.470 | 1.171 | 0.5765 | 0.834 | 506.00 | 572.00 | 9.84 | 2/8 |
| M394 | 41 | 217.73 | 10.361 | 1.281 | 0.4931 | 0.714 | 517.47 | 572.13 | 9.97 | 9/9 |

Table S17: Profile likelihoods of the best model (M343, all 41 estimated parameters). The 95 % interval is the likelihood-ratio interval $\{\theta \mid \Lambda(\theta) \leq \chi^2_1(0.95)\}$ read off the profile. The *bounds* column gives the calibration box of each parameter, so that each estimate and its interval can be read against the admissible range the literature audit of Section S1.6 set for it. *at bound* means the profile reached the box bound of the parameter before the threshold, *open* that it was still below the threshold where the path ended; both leave the interval one-sided. † marks a maximum-likelihood estimate sitting at a box bound. Ligand-relevant parameters are listed first.

| parameter | unit | estimate | bounds [lb, ub] | 95 % interval | width [dec.] | classification |
|---|---|---|---|---|---|---|
| BpA_c1_dna_ahrr | – | 9.08 | $[10^{-5}, 10^{5}]$ | [3.38, 36.1] | 1.03 | identifiable |
| BpA_const_L_degrad | $s^{-1}$ | 0.00259 | $[10^{-7}, 10^{-2}]$ | $[4.40\times10^{-4}$, at bound] | – | one-sided (lower) |
| tau_eL | $s^{-1}$ | $7.24\times10^{-5}$ | $[10^{-7}, 10^{-3}]$ | $[1.31\times10^{-5}$, at bound] | – | one-sided (lower) |
| k1_AhRL | $nM^{-1}\,s^{-1}$ | 0.126 | $[10^{-5}, 10]$ | [at bound, 1.36] | – | one-sided (upper) |
| k_diss_AhRL† | $nM^{-1}$ | 0.001 | $[10^{-3}, 100]$ | [at bound, 0.134] | – | one-sided (upper) |
| k1_dna_ahrr | $nM^{-1}\,s^{-1}$ | $3.19\times10^{-6}$ | $[10^{-6}, 1]$ | [at bound, 0.00136] | – | one-sided (upper) |
| k_diss_dna_ahrr | $nM^{-1}$ | 0.158 | $[10^{-4}, 100]$ | $[2.64\times10^{-4}, 1.57]$ | 3.78 | identifiable |
| rho_Cyp | $nM^{-1}$ | $2.44\times10^{6}$ | $[1, 10^{9}]$ | $[2.59\times10^{4}, 4.23\times10^{7}]$ | 3.21 | identifiable |
| b_AhRR† | $s^{-1}$ | 0.1 | $[10^{-6}, 0.1]$ | [0.022, at bound] | – | one-sided (lower) |
| b_AhRR_basal† | $nM\,s^{-1}$ | $1.00\times10^{-8}$ | $[10^{-8}, 10^{-2}]$ | [at bound, $5.76\times10^{-8}$] | – | one-sided (upper) |
| k1_AhRR_ARNT | $nM^{-1}\,s^{-1}$ | $1.51\times10^{-6}$ | $[10^{-6}, 0.1]$ | [at bound, at bound] | – | practically non-identifiable |
| k_diss_AhRR_ARNT | $nM^{-1}$ | 0.00169 | $[10^{-4}, 10^{3}]$ | [at bound, 0.199] | – | one-sided (upper) |
| k1_Cyp1a1† | $nM^{-1}\,s^{-1}$ | $1.00\times10^{-8}$ | $[10^{-8}, 0.1]$ | [at bound, $1.62\times10^{-7}$] | – | one-sided (upper) |
| k1_dna_cyp | $nM^{-1}\,s^{-1}$ | $1.16\times10^{-4}$ | $[10^{-6}, 1]$ | $[8.63\times10^{-6}$, at bound] | – | one-sided (lower) |
| k_diss_dna_cyp† | $nM^{-1}$ | $1.00\times10^{-4}$ | $[10^{-4}, 100]$ | [at bound, 0.025] | – | one-sided (upper) |
| k1_heterodim | $nM^{-1}\,s^{-1}$ | 0.00379 | $[10^{-6}, 0.1]$ | $[8.92\times10^{-6}$, at bound] | – | one-sided (lower) |
| k_diss_heterodim† | $nM^{-1}$ | 1000 | $[10^{-4}, 10^{3}]$ | [1.91, at bound] | – | one-sided (lower) |
| k1_Lentry | $s^{-1}$ | 0.00131 | $[10^{-4}, 10]$ | [at bound, 0.00287] | – | one-sided (upper) |
| k2_Lentry† | $s^{-1}$ | $1.00\times10^{-6}$ | $[10^{-6}, 10]$ | [at bound, $3.84\times10^{-5}$] | – | one-sided (upper) |
| k1_nuclearimport | $s^{-1}$ | $3.55\times10^{-4}$ | $[10^{-5}, 10^{-2}]$ | $[8.86\times10^{-5}, 0.00201]$ | 1.35 | identifiable |
| k1_rep_dna_ahrr | $nM^{-1}\,s^{-1}$ | 0.01 | $[10^{-6}, 1]$ | $[5.89\times10^{-5}, 0.884]$ | 4.18 | identifiable |
| k_diss_rep_dna_ahrr | $nM^{-1}$ | 14.6 | $[10^{-4}, 100]$ | [0.143, at bound] | – | one-sided (lower) |
| k1_rep_dna_cyp | $nM^{-1}\,s^{-1}$ | $5.50\times10^{-5}$ | $[10^{-6}, 1]$ | $[9.58\times10^{-6}, 5.62\times10^{-4}]$ | 1.77 | identifiable |
| k_diss_rep_dna_cyp† | $nM^{-1}$ | 100 | $[10^{-4}, 100]$ | [2.51, at bound] | – | one-sided (lower) |
| m_AhRR† | $s^{-1}$ | 1 | $[10^{-5}, 1]$ | [0.0119, at bound] | – | one-sided (lower) |
| m_Cyp† | $s^{-1}$ | $1.00\times10^{-5}$ | $[10^{-5}, 1]$ | [at bound, $1.62\times10^{-4}$] | – | one-sided (upper) |
| s_mRNA_Cyp1a1_1† | – | $1.00\times10^{-5}$ | $[10^{-5}, 100]$ | [at bound, $6.06\times10^{-5}$] | – | one-sided (upper) |
| s_mRNA_AhRR_1 | – | 3.28 | $[10^{-5}, 100]$ | [0.6, 22.3] | 1.57 | identifiable |
| tau_AhR | $s^{-1}$ | $3.08\times10^{-5}$ | $[10^{-6}, 10^{-4}]$ | $[1.57\times10^{-6}, 9.20\times10^{-5}]$ | 1.77 | identifiable |
| tau_AhRR | $s^{-1}$ | $3.88\times10^{-4}$ | $[10^{-7}, 10^{-3}]$ | [open, at bound] | – | practically non-identifiable |
| tau_Cyp1a1† | $s^{-1}$ | 0.001 | $[10^{-7}, 10^{-3}]$ | $[6.69\times10^{-5}$, at bound] | – | one-sided (lower) |
| tau_mRNA_AhRR† | $s^{-1}$ | $9.79\times10^{-4}$ | $[10^{-6}, 10^{-3}]$ | $[1.17\times10^{-4}$, at bound] | – | one-sided (lower) |
| tau_mRNA_Cyp | $s^{-1}$ | $4.03\times10^{-5}$ | $[10^{-5}, 10^{-3}]$ | $[2.55\times10^{-5}, 9.50\times10^{-5}]$ | 0.57 | identifiable |
| tau_nAhRL | $s^{-1}$ | $4.58\times10^{-4}$ | $[10^{-6}, 0.5\times10^{-3}]$ | $[9.92\times10^{-5}$, at bound] | – | one-sided (lower) |
| tau_nARNT | $s^{-1}$ | $6.42\times10^{-5}$ | $[10^{-7}, 10^{-3}]$ | $[9.63\times10^{-7}, 1.96\times10^{-4}]$ | 2.31 | identifiable |
| v_AhR | $nM\,s^{-1}$ | 0.00428 | $[10^{-8}, 0.1]$ | $[5.19\times10^{-5}, 0.02]$ | 2.59 | identifiable |
| v_nARNT | $nM\,s^{-1}$ | 0.0422 | $[10^{-8}, 0.1]$ | [0.00965, at bound] | – | one-sided (lower) |
| sigma0_mRNA_Cyp1a1 | – | 0.112 | $[10^{-5}, 10^{5}]$ | [at bound, 0.542] | – | one-sided (upper) |
| sigma1_mRNA_Cyp1a1 | – | 0.611 | $[10^{-5}, 10^{5}]$ | [0.463, 0.857] | 0.27 | identifiable |
| sigma0_mRNA_AhRR | – | 0.115 | $[10^{-5}, 10^{5}]$ | [at bound, 0.281] | – | one-sided (upper) |
| sigma1_mRNA_AhRR | – | 0.238 | $[10^{-5}, 10^{5}]$ | [0.168, 0.337] | 0.30 | identifiable |
