## Supplementary data for "Mathematical Modeling of the Canonical Aryl Hydrocarbon Receptor Pathway in murine bone-marrow-derived macrophages"

| Timepoint (h) | Relative change (2^-ΔCt) | | | Mean (±SEM) |
|---|---|---|---|---|
| | dmso / control | | | |
| 2 | 0.000523 | 0.00162 | 0.000681 | 9.41E-04 ± 0.00 |
| 3 | 0.001069 | 0.000565 | 0.000162 | 5.99E-04 ± 0.00 |
| 4 | 0.00286 | 0.004187 | 0.003285 | 3.44E-03 ± 0.00 |
| 6 | 0.00135 | 0.003006 | 0.004412 | 2.92E-03 ± 0.00 |
| 8 | 0.003198 | 0.003296 | 0.003475 | 3.32E-03 ± 0.00 |
| 24 | 0.00215216 | 0.00128858 | 0.000904871 | 1.45E-03 ± 0.00 |
| | 3MC | | | |
| 2 | 0.000956 | 0.003023 | 0.00113 | 1.70E-03 ± 0.00 |
| 3 | 0.004979 | 0.004809 | 0.001371 | 3.72E-03 ± 0.00 |
| 4 | 0.019237 | 0.013139 | 0.013888 | 1.54E-02 ± 0.00 |
| 6 | 0.007327 | 0.010127 | 0.01509 | 1.08E-02 ± 0.00 |
| 8 | 0.009053 | 0.005727 | 0.007936 | 7.57E-03 ± 0.00 |
| 24 | 0.00458139 | 0.00238797 | 0.002724109 | 3.23E-03 ± 0.00 |
| | ICZ | | | |
| 2 | 0.001575 | 0.004304 | 0.002705 | 2.86E-03 ± 0.00 |
| 3 | 0.004979 | 0.004044 | 0.001631 | 3.55E-03 ± 0.00 |
| 4 | 0.025033 | 0.025559 | 0.022561 | 2.44E-02 ± 0.00 |
| 6 | 0.005083 | 0.007793 | 0.017941 | 1.03E-02 ± 0.00 |
| 8 | 0.003972 | 0.006481 | 0.008426 | 6.29E-03 ± 0.00 |
| 24 | 0.00421574 | 0.00264962 | 0.00289946 | 3.25E-03 ± 0.00 |
| | BpA | | | |
| 2 | 0.00052 | 0.001874 | 0.000793 | 1.06E-03 ± 0.00 |
| 3 | 0.000874 | 0.000676 | 0.00026 | 6.03E-04 ± 0.00 |
| 4 | 0.000821 | 0.008315 | 0.007239 | 5.46E-03 ± 0.00 |
| 6 | 0.002577 | 0.004729 | 0.007148 | 4.82E-03 ± 0.00 |
| 8 | 0.002457 | 0.003446 | 0.00685 | 4.25E-03 ± 0.00 |
| 24 | 0.00114535 | 0.00095647 | 0 | 7.01E-04 ± 0.00 |

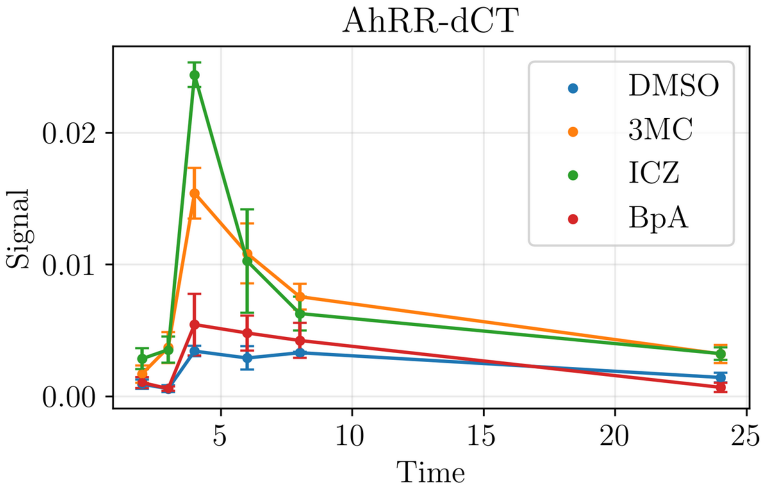

| Timepoint (h) | Normalized Relative change (2^-ΔΔCt) | | | Mean (±SEM) |
|---|---|---|---|---|
| | 3MC | | | |
| 2 | 1.82791587 | 1.86604938 | 1.659324523 | 1.78 ± 0.06 |
| 3 | 4.65762395 | 8.51150442 | 8.462962963 | 7.21 ± 1.28 |
| 4 | 6.72622378 | 3.13804633 | 4.227701674 | 4.70 ± 1.06 |
| 6 | 5.42740741 | 3.36892881 | 3.420217588 | 4.07 ± 0.68 |
| 8 | 2.83083177 | 1.73756068 | 2.283741007 | 2.28 ± 0.32 |
| 24 | 2.12874101 | 1.85317582 | 3.010494314 | 2.33 ± 0.35 |

| | ICZ | | | |
|---|---|---|---|---|
| 2 | 3.01147228 | 2.65679012 | 3.972099853 | 3.21 ± 0.39 |
| 3 | 4.65762395 | 7.15752212 | 10.06790123 | 7.29 ± 1.56 |
| 4 | 8.7527972 | 6.10437067 | 6.867884323 | 7.24 ± 0.79 |
| 6 | 3.76518519 | 2.5924817 | 4.066409791 | 3.47 ± 0.45 |
| 8 | 1.24202627 | 1.96632282 | 2.424748201 | 1.88 ± 0.34 |
| 24 | 1.95884085 | 2.0562277 | 3.204279947 | 2.41 ± 0.40 |

| | BpA | | | |
|---|---|---|---|---|
| 2 | 0.99426386 | 1.15679012 | 1.164464023 | 1.11 ± 0.06 |
| 3 | 0.81758653 | 1.19646018 | 1.604938272 | 1.21 ± 0.23 |
| 4 | 0.28706294 | 1.98590877 | 2.203652968 | 1.49 ± 0.61 |
| 6 | 1.90888889 | 1.57318696 | 1.620126927 | 1.70 ± 0.10 |
| 8 | 0.76829268 | 1.04550971 | 1.971223022 | 1.26 ± 0.36 |
| 24 | 0.53218537 | 0.74226165 | 0 | 0.42 ± 0.22 |

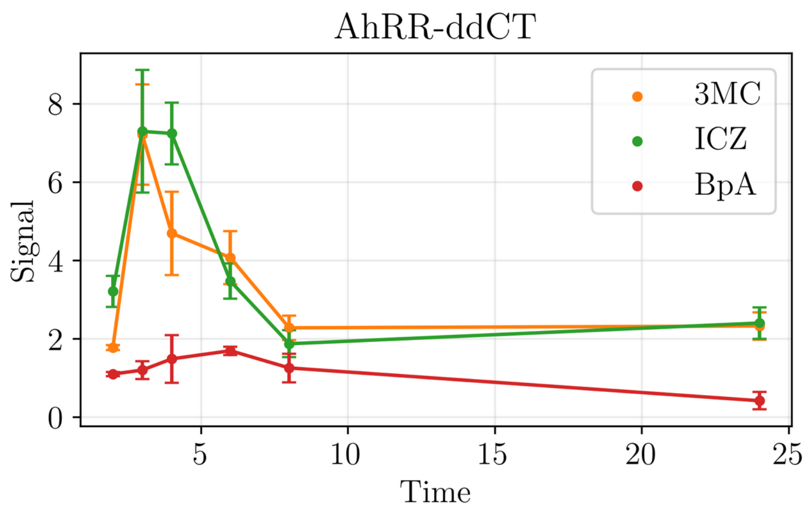

| Timepoint (h) | Relative change (2^-ΔCt) dmso / control | | | Mean (±SEM) |
|---|---|---|---|---|
| 2 | 3.74E-06 | 2.21E-06 | 5.08E-07 | 2.15E-06 ± 9.33E-07 |
| 3 | 1.85E-05 | 3.23E-06 | 4.38E-06 | 8.70E-06 ± 0.00 |
| 4 | 1.03E-05 | 6.11E-06 | 6.70E-06 | 7.70E-06 ± 0.00 |
| 6 | 7.36E-07 | 2.41E-06 | 7.60E-06 | 3.58E-06 ± 0.00 |
| 8 | 8.11E-07 | 2.93E-07 | 5.47E-07 | 5.50E-07 ± 1.50E-07 |
| 24 | 8.96E-07 | 1.19E-06 | 2.62E-06 | 1.57E-06 ± 5.34E-07 |

| | 3MC | | | |
|---|---|---|---|---|
| 2 | 1.85E-05 | 4.56E-05 | 6.73E-06 | 2.36E-05 ± 0.00 |
| 3 | 0.000296 | 5.03E-05 | 6.10E-05 | 1.36E-04 ± 0.00 |
| 4 | 0.000129 | 0.000065 | 3.39E-05 | 7.60E-05 ± 0.00 |
| 6 | 1.05E-05 | 1.58E-05 | 1.63E-05 | 1.42E-05 ± 0.00 |
| 8 | 2.28E-05 | 2.54E-06 | 3.71E-06 | 9.68E-06 ± 0.00 |
| 24 | 6.55E-06 | 4.26E-06 | 9.20E-06 | 6.67E-06 ± 0.00 |

| | ICZ | | | |
|---|---|---|---|---|
| 2 | 4.82E-05 | 0.000103 | 8.12E-06 | 5.31E-05 ± 0.00 |
| 3 | 0.000322 | 6.32E-05 | 5.66E-05 | 1.47E-04 ± 0.00 |
| 4 | 0.000234 | 0.000705 | 2.63E-04 | 4.01E-04 ± 0.00 |
| 6 | 2.64E-05 | 2.52E-05 | 4.81E-05 | 3.32E-05 ± 0.00 |
| 8 | 2.67E-05 | 1.33E-05 | 2.68E-05 | 2.23E-05 ± 0.00 |
| 24 | 1.11E-05 | 5.1E-06 | 2.14E-05 | 1.25E-05 ± 0.00 |

| | BpA | | | |
|---|---|---|---|---|
| 2 | 3.34E-06 | 7.47E-06 | 8.13E-07 | 3.87E-06 ± 0.00 |
| 3 | | | | |
| 4 | 2.22E-05 | 2.27E-05 | 7.68E-06 | 1.75E-05 ± 0.00 |
| 6 | 2.44E-06 | 6.48E-07 | 1.17E-06 | 1.42E-06 ± 5.32E-07 |
| 8 | 1.06E-06 | 2.36E-07 | 1.02E-06 | 7.72E-07 ± 2.68E-07 |
| 24 | 8.24E-07 | 1.98E-07 | 0.00E+00 | 3.41E-07 ± 2.49E-07 |

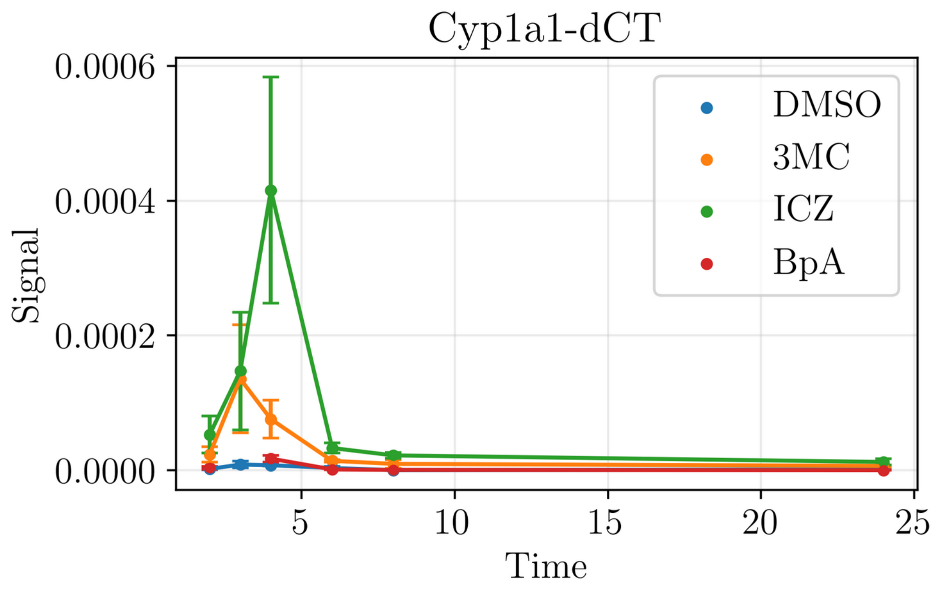

| | Normalized Relative change (2^-ΔΔCt) | | | Mean (±SEM) |
|---|---|---|---|---|
| Timepoint (h) | 3MC | | | |
| 2 | 4.946524 | 20.63348 | 13.2480315 | 12.94 ± 4.53 |
| 3 | 16 | 15.57276 | 13.9269406 | 15.17 ± 0.63 |
| 4 | 12.52427 | 10.6383 | 5.05970149 | 9.41 ± 2.24 |
| 6 | 14.2663 | 6.556017 | 2.14473684 | 7.66 ± 3.54 |
| 8 | 28.11344 | 8.668942 | 6.78244973 | 14.52 ± 6.82 |
| 24 | 7.310644 | 3.580101 | 3.50641856 | 4.80 ± 1.26 |
| | ICZ | | | |
| 2 | 12.8877 | 46.60633 | 15.984252 | 25.16 ± 10.76 |
| 3 | 17.40541 | 19.56656 | 12.9223744 | 16.63 ± 1.96 |
| 4 | 22.71845 | 115.3846 | 39.2537313 | 59.12 ± 28.53 |
| 6 | 35.86957 | 10.45643 | 6.32894737 | 17.55 ± 9.24 |
| 8 | 32.92232 | 45.39249 | 48.9945155 | 42.44 ± 4.87 |
| 24 | 12.38048 | 4.287098 | 8.16807946 | 8.28 ± 2.34 |
| | BpA | | | |
| 2 | 0.893048 | 3.38009 | 1.6003937 | 1.96 ± 0.74 |
| 3 | | | | |
| 4 | 2.15534 | 3.715221 | 1.14626866 | 2.34 ± 0.75 |
| 6 | 3.315217 | 0.26888 | 0.15394737 | 1.25 ± 1.04 |
| 8 | 1.307028 | 0.805461 | 1.86471664 | 1.33 ± 0.31 |
| 24 | 0.920187 | 0.166086 | 0 | 0.36 ± 0.28 |

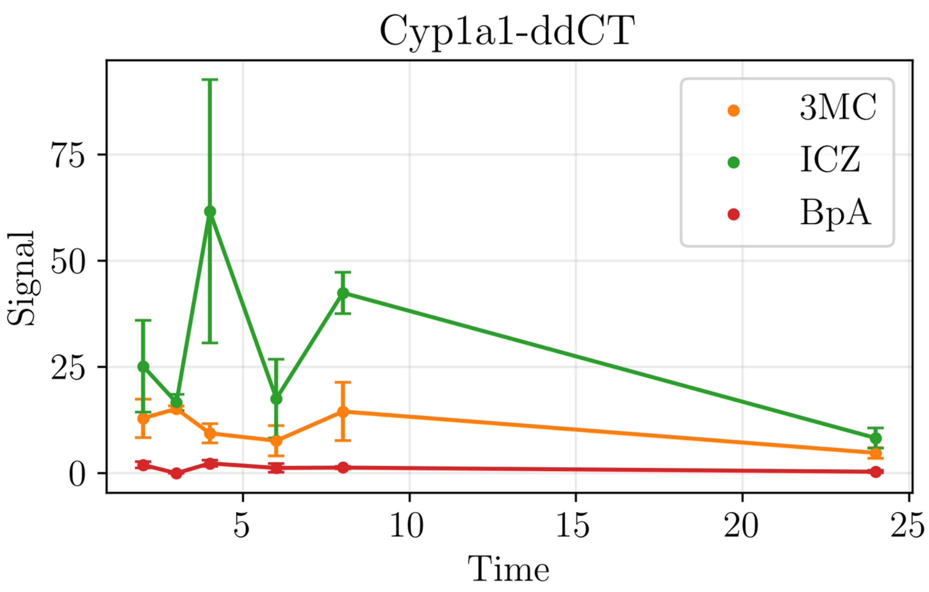

| *Cyp1a1* | | | | | | |
|---|---|---|---|---|---|---|
| | dmso | | | 3MC | | |
| Timepoint (h) | Replicate 1 | Replicate 2 | Replicate 3 | Replicate 1 | Replicate 2 | Replicate 3 |
| 2 | 0.00000374 | 0.00000221 | 0.000000508 | 0.0000185 | 0.0000456 | 0.00000673 |
| 3 | 0.0000185 | 0.00000323 | 0.00000438 | 0.000296 | 0.0000503 | 0.000061 |
| 4 | 0.0000103 | 0.00000611 | 0.0000067 | 0.000129 | 0.000065 | 0.0000339 |
| 6 | 0.000000736 | 0.00000241 | 0.0000076 | 0.0000105 | 0.0000158 | 0.0000163 |
| 8 | 0.000000811 | 0.000000293 | 0.000000547 | 0.0000228 | 0.00000254 | 0.00000371 |
| 24 | 0.000000895999 | 0.0000011905 | 0.00000262364 | 0.00000655033 | 0.00000426211 | 0.00000919958 |
| | ICZ | | | BpA | | |
| Timepoint (h) | Replicate 1 | Replicate 2 | Replicate 3 | Replicate 1 | Replicate 2 | Replicate 3 |
| 2 | 0.0000482 | 0.000103 | 0.00000812 | 0.00000334 | 0.00000747 | 0.000000813 |
| 3 | 0.000322 | 0.0000632 | 0.0000566 | 0.0000174 | 0.00000105 | 0.00000213 |
| 4 | 0.000234 | 0.000705 | 0.000263 | 0.0000222 | 0.0000227 | 0.00000768 |
| 6 | 0.0000264 | 0.0000252 | 0.0000481 | 0.00000244 | 0.000000648 | 0.00000117 |
| 8 | 0.0000267 | 0.0000133 | 0.0000268 | 0.00000106 | 0.000000236 | 0.00000102 |
| 24 | 0.0000110929 | 0.00000510379 | 0.0000214301 | 0.000000824487 | 0.000000197725 | 0 |

| *AhRR* | | | | | | |
|---|---|---|---|---|---|---|
| | dmso | | | 3MC | | |
| Timepoint (h) | Replicate 1 | Replicate 2 | Replicate 3 | Replicate 1 | Replicate 2 | Replicate 3 |
| 2 | 0.000523 | 0.00162 | 0.000681 | 0.000956 | 0.003023 | 0.00113 |
| 3 | 0.001069 | 0.000565 | 0.000162 | 0.004979 | 0.004809 | 0.001371 |
| 4 | 0.00286 | 0.004187 | 0.003285 | 0.019237 | 0.013139 | 0.013888 |
| 6 | 0.00135 | 0.003006 | 0.004412 | 0.007327 | 0.010127 | 0.01509 |
| 8 | 0.003198 | 0.003296 | 0.003475 | 0.009053 | 0.005727 | 0.007936 |
| 24 | 0.002152158 | 0.001288582 | 0.000904871 | 0.004581387 | 0.002387969 | 0.002724109 |
| | ICZ | | | BpA | | |
| Timepoint (h) | Replicate 1 | Replicate 2 | Replicate 3 | Replicate 1 | Replicate 2 | Replicate 3 |
| 2 | 0.001575 | 0.004304 | 0.002705 | 0.00052 | 0.001874 | 0.000793 |
| 3 | 0.004979 | 0.004044 | 0.001631 | | | |
| 4 | 0.025033 | 0.025559 | 0.022561 | 0.000821 | 0.008315 | 0.007239 |
| 6 | 0.005083 | 0.007793 | 0.017941 | 0.002577 | 0.004729 | 0.007148 |
| 8 | 0.003972 | 0.006481 | 0.008426 | 0.002457 | 0.003446 | 0.00685 |
| 24 | 0.004215735 | 0.002649618 | 0.00289946 | 0.001145347 | 0.000956465 | 0 |

| **Conc. substances** |
|---|
| 1 µM DMSO |
| 1 µM 3MC |
| 10 µM ICZ |
| 1µM BpA |
